# Dysproteostasis primes pancreatic epithelial state changes in *KRAS*-mediated oncogenesis

**DOI:** 10.1101/2025.03.24.644918

**Authors:** Carla Salomo Coll, Marisa di Monaco, Jocelyn Holkham, Matt Smith, Morwenna Muir, Philippe Gautier, Xiaozhong Zheng, Roopesh Krishnankutty, Alain J. Kemp, Katie Winnington-Ingram, Alex von Kriegsheim, Jennifer P. Morton, Natalia Jimenez-Moreno, Damian Mole, Simon Wilkinson

## Abstract

Pre-malignant transformation of pancreatic acinar cells by oncogenic *Kras* is dependent upon stochastic emergence of metaplastic cell states, through unknown mechanisms. We reveal that an early, transcriptionally-mediated effect of *Kras* is sporadic failure of proteostatic ER-phagy. Genetically-altered mice deficient in ER-phagy demonstrate that this event co-operates with *Kras* to drive acinar-ductal metaplasia (ADM) and subsequent cancer. Mechanistically, proteomics and high-resolution imaging uncover pathologic aggregation of a subset of ER proteins, including the injury marker REG3B, resulting from failure to physically interact with the ER-phagy receptor CCPG1. Spatial transcriptomics demonstrate that the appearance of sporadic intracellular aggregates upon *Kras* activation marks rare acinar cells existing in an injured, ADM-primed state. Importantly, engineered mutants of REG3B establish that aggregate formation is sufficient to directly engender this epithelial cell state. Pancreatic cancer can thus arise from stochastic pathologic protein aggregates that are influenced by and co-operate with an oncogene.

## Introduction

Oncogenic mutations can be tolerated indefinitely by epithelial cells within some tissues, without apparent phenotypic consequence (Martincorena et al., 2015; Neuhofer et al., 2021). Such cells exhibit only a low probability of transformation to cancer precursor states. It is widely presumed that additional stochastic events co-operate with oncogene mutation to facilitate initial progression towards cancer. Identification of these events is a necessary step in the assembly of a molecular mechanistic description of cancer risk. Their discovery would greatly enhance the development of preventative strategies. Pancreatic ductal adenocarcinoma (PDAC) is a notable exemplar of this concept. PDAC can originate in the exocrine pancreatic epithelium, from constituent acinar or ductal cells (Guerra et al., 2007; Habbe et al., 2008; Gidekel Friedlander et al., 2009; Guerra et al., 2011; Kopp et al., 2012; Bailey et al., 2016; Ferreira et al., 2017; Storz, 2017; Lee et al., 2019; Neuhofer et al., 2021; Braxton et al., 2024). In greater than 90 % of cases this occurs after initial mutation of the *Kras* oncogene. However, acinar cells can harbour the *Kras* oncogene long prior to morphological state changes, overt hyperproliferation or other canonical responses such as senescence (Guerra et al., 2007; Habbe et al., 2008; Guerra et al., 2011; Ardito et al., 2012; Navas et al., 2012; Alonso-Curbelo et al., 2021; Li et al., 2021; Neuhofer et al., 2021).

Externally-imposed, pancreas-wide injury triggers acinar cells to undergo a reversible state change termed acinar-ductal metaplasia (ADM) (Storz, 2017). In mouse models, oncogenic *Kras* diminishes resolution of injury-induced ADM, resulting in persistent pseudo-ductal cell states that can seed premalignant ductal lesions (pancreatic intraepithelial neoplasia, PanIN) (Morris et al., 2010; Kopp et al., 2012; Alonso-Curbelo et al., 2021; Li et al., 2021). Furthermore, previous resolution of experimental injury primes wild-type cells for ADM upon future *Kras* mutation (Del Poggetto et al., 2021; Falvo et al., 2023). These injury models thus reflect the role of chronic pancreatitis - which prefigures a small minority of human PDAC cases (Kirkegard et al., 2017) – as a co-operating event in oncogene-mediated evolution of ADM and PanIN. However, in the majority of PDAC that are not preceded by overt injury and inflammation, the stochastic co-operating event(s) remain unknown. Modelling this, only a small minority of acinar cells bearing oncogenic *Kras* will spontaneously undergo irreversible ADM in the absence of externally-imposed injury (Hingorani et al., 2003; Habbe et al., 2008). This raises the question of the identity of molecular events that substitute for externally-imposed injury in order to co-operate with oncogenic *Kras*.

A hypothetical co-operating event in the absence of external injury could be sporadic impairment of acinar cell homeostasis, which could in turn generate pathologic cell state(s) primed for ADM. In this regard, it has been observed that experimental deletion of autophagy genes – which compromises bulk lysosomal catabolism of cytoplasm - drives catastrophic inflammation and sensitisation to ADM (Rosenfeldt et al., 2013). There is scant evidence of autophagy suppression as a foundational event in *Kras*-driven PDAC; indeed, oncogenic *Kras* is a known stimulant for bulk autophagy (Guo et al., 2011; Lock et al., 2011) and at least some genetic subtypes of established PDAC rely on autophagy for survival (Yang et al., 2011; Yang et al., 2014; Perera et al., 2015). However, it is striking that the role of selective autophagy has not been addressed. Selective autophagy refers to the emerging conceptualisation of autophagy as multiple, discrete pathways that target different organelles for degradation (Vargas et al., 2023). *In vivo*, this is expected to maintain cellular and organ homeostasis in a substantially tissue-specific manner (McWilliams et al., 2018). In particular, there is growing evidence for the importance in pancreatic acinar cells of ER-phagy, the selective autophagic degradation of parts of the endoplasmic reticulum (ER) network (Smith et al., 2018). Mechanistically, ER-phagy is specified by “cargo receptor” proteins in the ER membrane, including CCPG1, FAM134B, TEX264 and SEC62, which bind to cytoplasmic autophagy proteins (Khaminets et al., 2015; Fumagalli et al., 2016; Grumati et al., 2017; Smith et al., 2018; An et al., 2019; Chino et al., 2019; Nthiga et al., 2020; Stephani et al., 2020). Indeed, loss of *Ccpg1* function by targeted gene disruption results in protein granule accumulation within the pancreatic acinar ER, suggesting a primary or secondary proteostatic failure (Smith et al., 2018). However, the molecular basis of such defects is unclear, as are their physiological sequelae and, crucially, whether or not these aberrations occur pathologically.

Here, we generate a reporter for ER-phagy in mouse pancreas, demonstrating heterogenous basal activity across the acinar epithelial compartment. However, ER-phagy suppression occurs as an early transcriptional response to *Kras* mutation and - in isolated regions where this is most penetrant - correlates spatially with the stochastic emergence of ADM. Furthermore, genetic ablation of ER-phagy shows that suppression of this selective autophagy pathway has a causal role in premalignant lesion evolution, co-operating with *Kras* in sequential generation of microinflammation and ADM, PanIN, and, ultimately, cancer. Mechanistically, proteomic analysis identifies a cohort of tissue-specifically expressed ER proteins (“clients”) that require ER-phagy for turnover and maintenance of solubility. A notable exemplar is the stereotypical injury marker REG3B, which we find physically binds the CCPG1 receptor in the ER lumen. In cells where ER-phagy is suppressed by *Kras*, compositionally- and morphologically-distinct luminal protein aggregates form, composed of these ER-phagy clients. Notably, this readout of proteostatic failure is detected sporadically in morphologically-normal acinar cells proximal to ADM, in both mouse and human pancreata. This observation is further consistent with the reduced ER-phagy in these regions. Finally, using spatial transcriptomics, we reveal that protein aggregates mark a rare acinar cell state that is indicative of localised, low-level injury and is primed for ADM. Importantly, by direct generation of REG3B aggregates upon ectopic expression of insoluble protein *in vivo*, we show that protein aggregates are sufficient to directly drive the state changes and ADM observed within this subpopulation of cells, downstream of *Kras*.

Remarkably, thus, pancreatic cancer is initiated by the co-operation of the *Kras* oncogene with stochastic deficiency in ER-phagy and proteostasis. Furthermore, oncogenic signalling greatly enhances the probability of pathologic protein aggregate formation via this route in the first instance.

## Results

### ER-phagy is heterogeneously suppressed by oncogenic *Kras* in pancreatic acinar cells

To determine whether oncogene activation might affect ER-phagy, cargo receptor expression was analysed in *Pdx1-Cre Kras^LSL-G12D/+^* (*KC*) mice, which have conditional pancreatic epithelial expression of the most common mutant of *Kras* found in pancreatic cancer, G12D (Fig. 1A-C). We observed that CCPG1, SEC62 and FAM134B protein abundances were reduced relative to *Pdx1-Cre Kras^+/+^* (*C*) controls (Fig. 1A-B). This reduction is consistent with the canonical view that *Kras* activation increases generalised autophagy flux and thus cargo receptor degradation. However, suppression of cognate transcript levels was also observed, highlighting an opposing possibility of transcriptionally-mediated inhibition of ER-phagy (Fig. 1C). Although ER-phagy is proposed to have a role in pancreatic acinar homeostasis (Smith et al., 2018), it has not previously been possible to directly measure this *in vivo*. Thus, we developed a technique employing recombinant adeno-associated virus (rAAV)-mediated delivery of a state-of-the-art ER-targeted fluorescent reporter, ss-TOLLES-YPet-KDEL (Supp. Fig. 1A-B). YPet fluorescence is suppressed and TOLLES is enhanced when the ER is delivered to acidic lysosomes (Katayama et al., 2020; Jimenez-Moreno et al., 2023). TOLLES-only foci are thus indicative of ER-phagy flux and can be counted to provide individual acinar cell ER-phagy indices. Notably, such TOLLES-only foci were readily detected within acinar cells in control *C* mice (Fig. 1D-E, Supp. Fig. 1B), indicating basal ER-phagy flux. When quantified, substantial inter- and intra-lobular spatial heterogeneity was evident (Fig. 1F-G). In *KC* animals, heterogeneity of basal ER-phagy across normal lobules remained evident but an overall reduction was detected (Fig. 1D-G). Furthermore, in rare lobules encompassing sporadic ADM (peri-ADM), ER-phagy within the morphologically-normal acinar cells therein was homogenously and substantially suppressed (Fig. 1D-G). In line with the previously observed loss of cargo receptor expression, this deficiency in ER-phagy was attributable to defects in autophagy initiation events, i.e., formation of ER-containing autophagosomes (Supp. Fig. 1C).

**Fig. 1.**
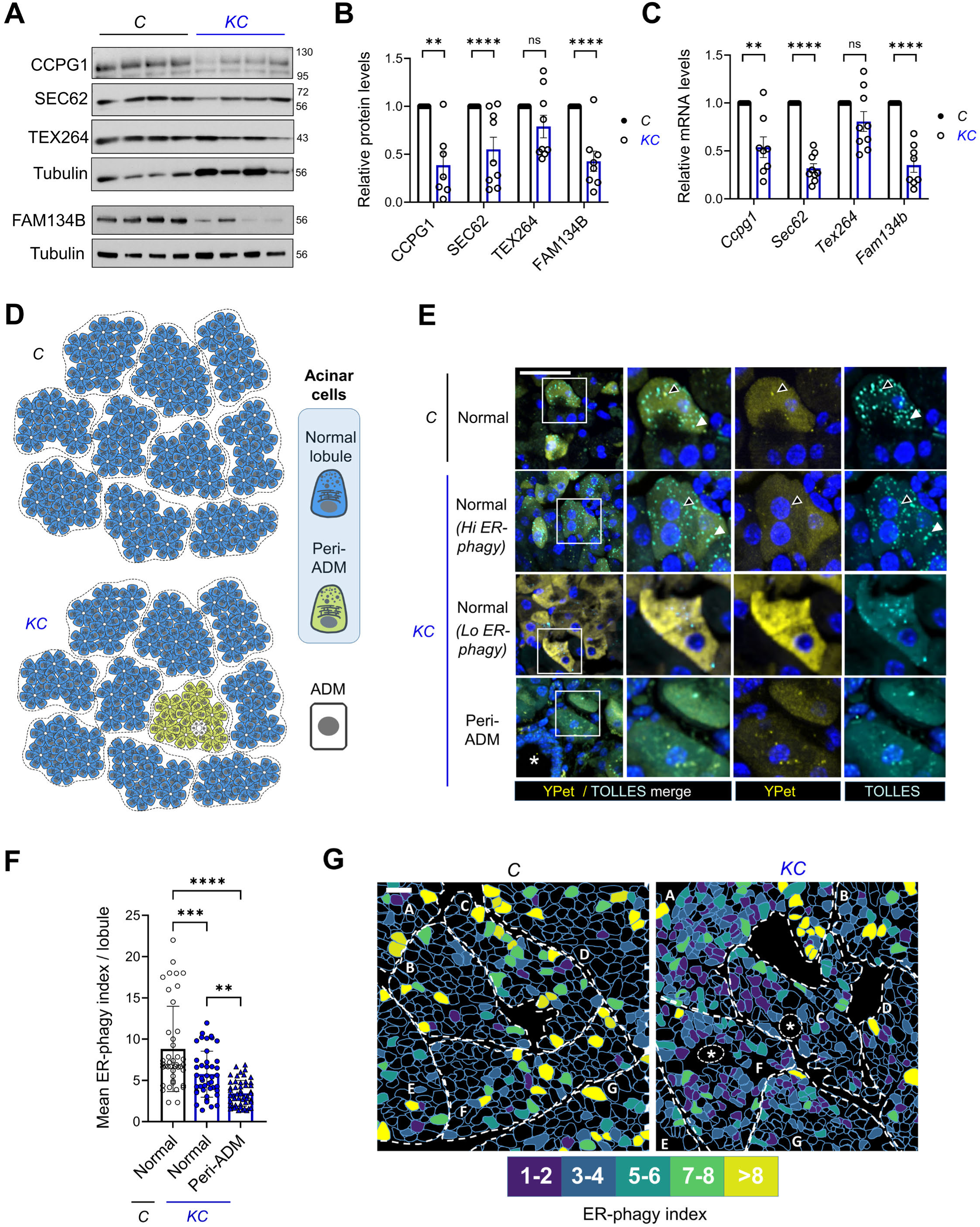
ER-phagy is heterogeneously suppressed by oncogenic *Kras* in pancreatic acinar cells. **A, B)** Immunoblotting of whole pancreatic lysates reveals reduced abundance of ER-phagy receptors in 10-week-old *KC* (*Pdx1-Cre Kras^LSL-G12D/+^)* mice compared with *C* (*Pdx1-Cre Kras ^+/+^*) controls (n = 7-9). **C)** qRT-PCR of whole pancreatic RNA reveals the transcriptional basis of reduced ER-phagy receptor abundance (n= 8-9). (Values normalised to *C* mice, ± S.E.M., 1-sample t-tests, ** = p ≤ 0.01, **** = p ≤ 0.0001, ns = p > 0.05). **D)** Schematic of pancreatic acinar lobules in control *C* mice and in *KC* mice. The latter is divided into a majority of normal lobules and minority of lobules that exhibit sporadic ADM embedded amongst morphologically-normal acinar cells (“peri-ADM” lobules). **E-F)** Representative spinning-disk confocal microscopy images and quantitative analyses of ER-phagy flux in acinar cells of 18-week-old *C* and *KC* mice, two weeks post-injection with rAAV expressing the ER-phagy flux reporter ss-YPet-TOLLES-KDEL (black arrowhead: bifluorescent YPet-TOLLES focus, white arrowhead: autolysosomal TOLLES-only focus, asterisk: ADM). (total n = 135 lobules from 5 pairs of mice, ± S.D., 1-way ANOVA and Holm-Šidák post-hoc test, ** = p ≤ 0.01, *** = p ≤ 0.001, **** = p ≤ 0.0001). **G)** Schematic maps of representative cross-sectional images of pancreata analysed in **E-F**. Lobules (labelled A-G) are circumscribed by broken white lines. Individual reporter-expressing acinar cells are colour coded according to ER-phagy index (TOLLES-only focus number on a per cell basis). The representative *KC* section demonstrates normal lobules (A,B,D,E) and “peri-ADM” lobules harbouring sporadic ADM (C,F; ADM represented by encircled asterisks). Scale bars = 50 μm.

Taken together, the above data show that an early effect of oncogenic *Kras* activation in pancreatic acinar cells is ER-phagy suppression. Furthermore, spatial correlation of the magnitude of ER-phagy deficiency with ADM highlights a possible causal link, wherein *Kras*- driven defects in ER homeostasis may promote sporadic ADM.

### ER-phagy function suppresses ADM and tumorigenesis

As a first step toward testing whether ER-phagy suppresses *Kras*-driven ADM and cancer, we first used our ER-phagy flux reporter to establish that basal activity was indeed suppressed upon whole animal loss of *Ccpg1* function (Fig. 2A-B, Supp. Fig. 2A; germline *Ccpg1* deficiency, *Ccpg1^GT/GT^*). Then we bred these *Ccpg1*-deficient mice with the *KPC* model of pancreatic cancer (*KC p53^LSL-R172H/+^*), observing acceleration of PDAC development (Fig. 2C). Having established acceleration of cancer in the absence of ER-phagy, we then dissected the stage of tumorigenesis affected. For these subsequent experiments, we also generated a new conditional (*loxP*) allele of *Ccpg1* (*Ccpg1^flox^*, Supp. Fig. 2B-D). This enabled us to discern whether early events in tumorigenesis were dependent upon pancreatic epithelial ER-phagy, as opposed to disruption of this pathway in other cell lineages. We observed that pancreatic epithelial deletion of *Ccpg1* combined with oncogenic *Kras* (*KC Ccpg1^flox/flox^,* hereafter *KC Ccpg1^ΔPANC^*) promoted early onset of ADM, followed by progression to PanIN at later time points (Fig. 2D-F). Inflammation promotes ADM and, while no evidence of gross inflammation was observed in younger *Ccpg1-*deficient *KC* mice, prior to the appearance of substantial metaplasia (10 weeks-old), increased macrophage abundance was already detectable immunohistologically, suggesting the existence of microinflammation (Fig. 2G-H). Finally, to gain insight into whether increased ADM abundance upon *Ccpg1* deletion was consequent from either supernumerary ductal metaplastic events or decreased resolution of ADM (back to morphologically-normal acinar cells), we provoked metaplasia synchronously, via intraperitoneal caerulein injection. In untreated animals, no ADM nor inflammation was detected (Fig. 2I-L). However, while initial responses (day 2) in both control groups and in *Ccpg1*-deficient animals were similar, resolution of ADM and inflammation at day 7 did not occur in the absence of *Ccpg1* (Fig. 2I-L).

**Fig. 2.**
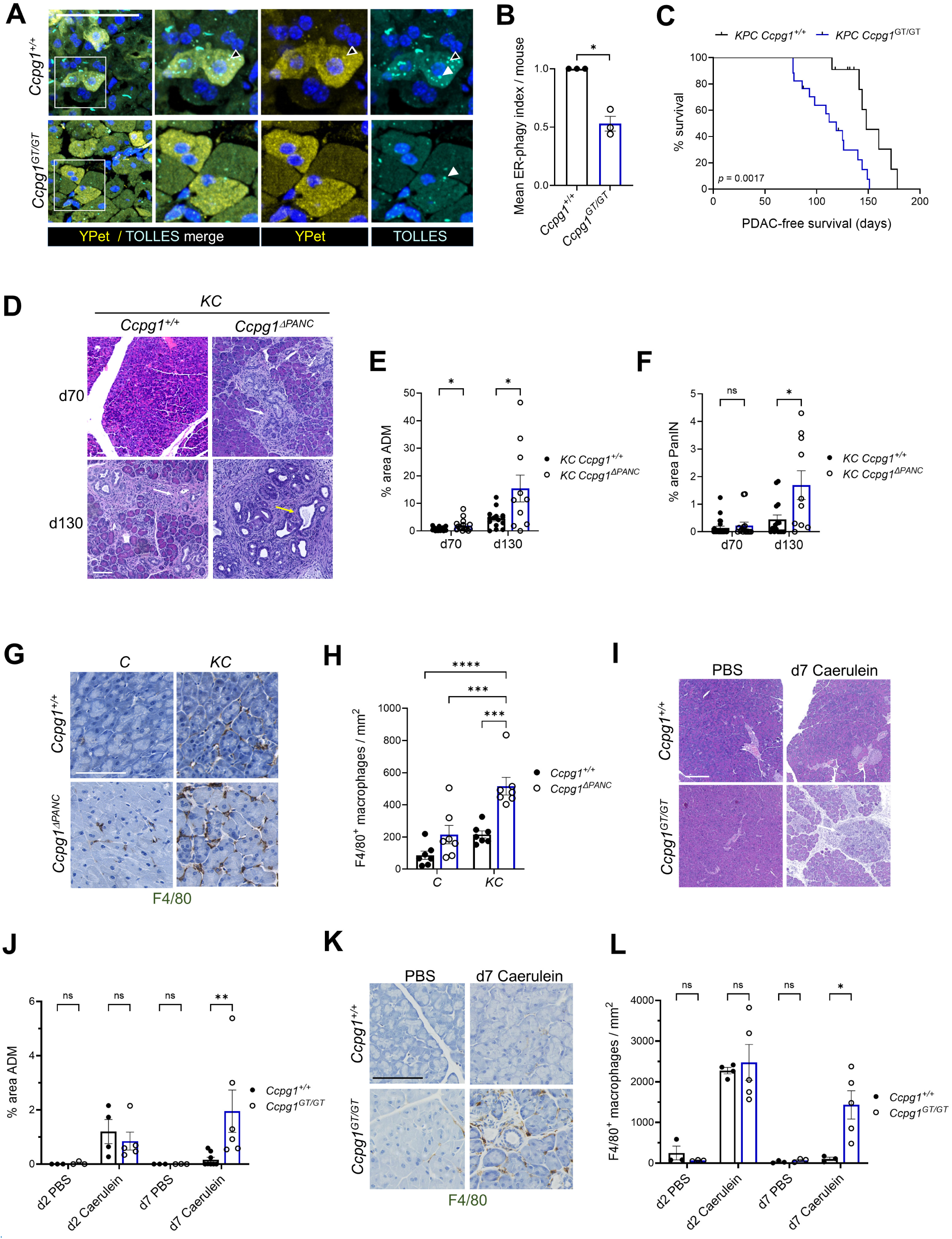
ER-phagy function suppresses ADM and tumorigenesis. **A-B)** ER-phagy flux in acinar cells of 8-week-old *Ccpg1*-deficient mice (*Ccpg1^GT/GT^*), as detected with ER-phagy reporter ss-YPet-TOLLES-KDEL (as per Fig. 1E-G, black arrowhead: bifluorescent YPet-TOLLES focus, white arrowheads: autolysosomal TOLLES-only foci). (n = 3 mice, 53 total microscopic fields, mean TOLLES-only foci per acinar cell, normalised to sibling *Ccpg1^+/+^*mice, ± S.E.M., Student’s t-test, * = p ≤ 0.05). **C)** Morbidity due to PDAC upon *Ccpg1* deficiency, shown by Kaplan-Meier survival plot of *KPC* mice *(Pdx1-Cre Kras^LSL-G12D/+^ Tp53^LSL-R172H/+^)*, comparing controls (*Ccpg1^+/+^*) with germline *Ccpg1* loss-of-function (*Ccpg1^GT/GT^*) (n = 11 and 17, respectively, upticks = censored mice). **D-F)** Pre-malignant lesions in ageing *KC* mice with pancreatic epithelial loss of *Ccpg1* function (*Ccpg1*^Δ*PANC*^), representative microscopic fields exhibiting accelerated ADM and PanIN are shown as H & E images in **D** (d70/130 = day 70 or 130 of age, white arrows: example ADM, yellow arrow: example low-grade PanIN) and quantified in **E** and **F** (d70: n = 23 and 16, d130: n = 15 and 10, ± S.E.M., Student’s t-tests, * = p ≤ 0.05, ns = p > 0.05). **G-H)** Microinflammation detection across morphologically-normal pancreatic regions, via staining for macrophages (F4/80^+^) in 10-week-old *C* (control) and *KC* (*Kras* mutant) mice, wild-type or deficient for *Ccpg1* (*Ccpg1*^Δ*PANC*^) in the pancreatic epithelium, shown in representative images in **G** and quantified in **H** (n = 7, ± S.E.M., 2-way ANOVA and Holm-Šidák post-hoc tests, *** = p ≤ 0.001, **** = p ≤ 0.0001, not shown: p > 0.05). **I-L)** Persistent ADM and inflammation in 7-week-old *Ccpg1*-deficient (*Ccpg1^GT/GT^*) mice (no *Kras* mutation) after 6 hourly i.p. caerulein injections, compared with PBS sham. Representative H & E images of ADM in **I** (arrow: region of multiple ADM), quantified in **J** (n = 3 PBS, n =5 caerulein at 2 days post-injection (d2); n = 3 PBS, n = 7 caerulein at d7) and IHC for macrophages in **K**, quantified in **L** (n = 3 PBS, n =5 caerulein at d2; n = 3 PBS, n = 5 caerulein at d7). All quantifications expressed ± S.E.M. (2-way ANOVA and Holm-Šidák post-hoc tests, * = p ≤ 0.05, ns = P > 0.05). Scale bars = 100 μm.

Taken together, the above data strongly suggest that ER-phagy suppresses *Kras*-driven microinflammation, stochastic and irreversible ADM, and eventual cancer formation.

### ER-phagy suppresses accumulation of specific ER luminal proteins in insoluble form

Given previous observations suggesting links between ER-phagy and acinar cell proteostasis, we hypothesised that defective proteostasis facilitates *Kras-*driven ADM. Thus, we first surveyed the proteomic consequences of ER-phagy disruption in the presence of oncogenic *Kras*. We isolated acini from *KC Ccpg1^+/+^* control and *KC Ccpg1^ΔPANC^* animals, and subjected detergent-soluble and -insoluble fractions to label-free mass spectrometric quantification (Supp. Table 1). When *Ccpg1* was ablated, a subset of ER luminal proteins increased in abundance across both fractions, implying a subproteome dependent upon ER-phagy for clearance and solubility (Fig. 3A, Supp. Fig. 3A). This finding was cross-validated by immunoblot of differentially-soluble fractions of fresh whole pancreas (Fig. 3B-C, Supp. Fig. 3B).

**Fig. 3.**
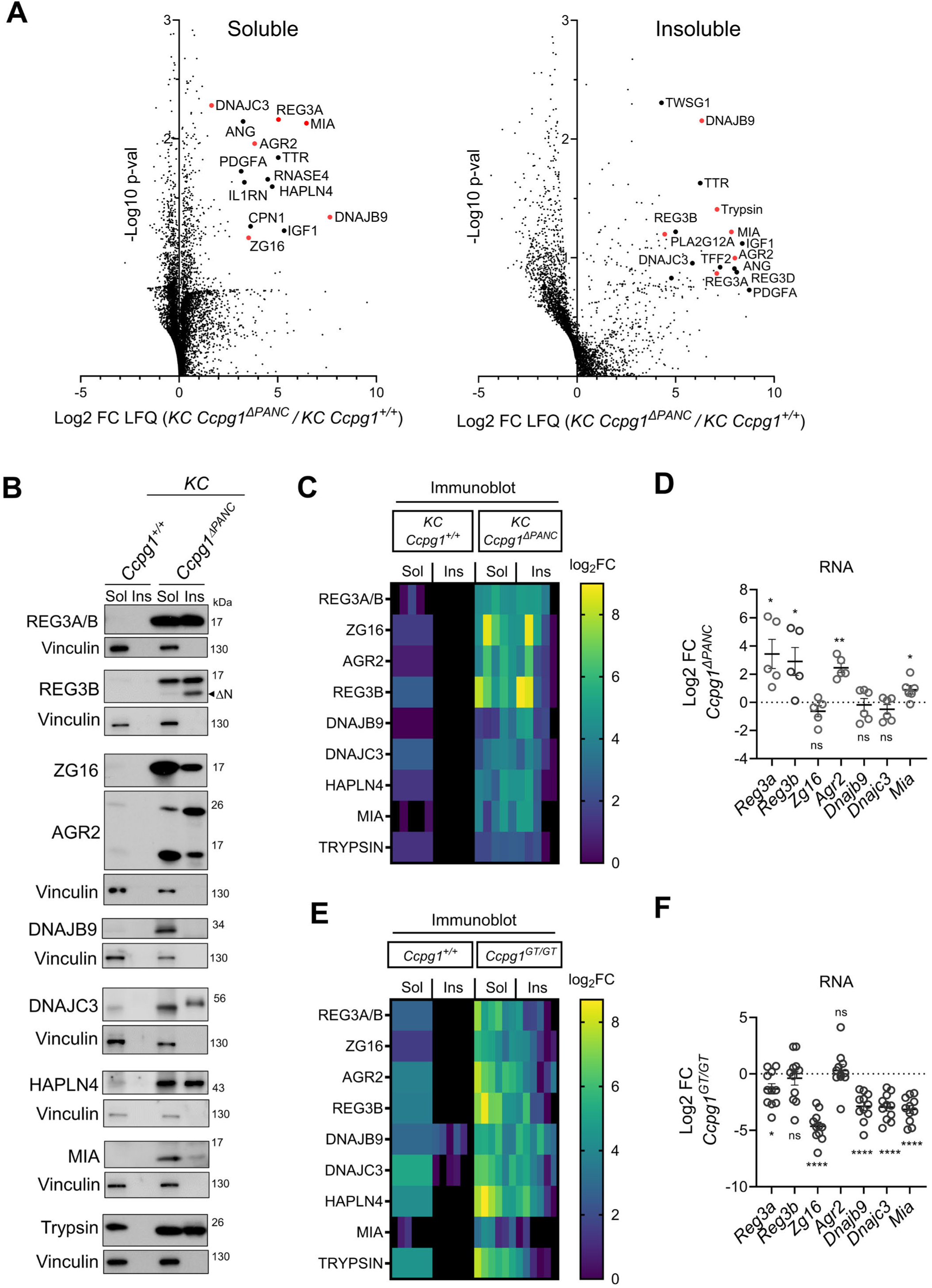
ER-phagy suppresses accumulation of specific ER luminal proteins in insoluble form. **A)** Volcano plots showing differential abundance of detergent (SDS)-soluble and -insoluble proteins in acini isolated from pancreata of 10-week-old *KC Ccpg1*^ΔPANC^ and *KC Ccpg1*^+/+^ mice (n = 3, FC = fold change, LFQ = label-free quantification, p-val = p value, 1 sample t-test on log2 ratios of LFQ values, red coloration highlights proteins amenable to immunoblot validation in subsequent panels). **B-C)** Orthogonal validation of differential protein abundance observed in **A** by immunoblot of whole pancreatic extracts from pancreata of 10-week-old *KC Ccpg1*^Δ^*^PANC^* and *KC Ccpg1^+/+^* mice. Representative blot shown in **B**, normalised quantifications of replicates summarised in **C** (n = 5, Sol/Insol = detergent (SDS)-soluble/insoluble, ΔN = N-terminally processed REG3B, FC = fold change). **D)** qRT-PCR analysis of whole pancreatic RNA from 10-week-old *KC Ccpg1*^Δ^*^PANC^* mice, expressed normalised to *KC Ccpg1^+/+^* mice (n = 5-6, ± S.E.M., 1-sample t-tests on untransformed values, * = p ≤ 0.05, ** = p ≤ 0.01, ns = p > 0.05). **E-F)** Normalised quantification of immunoblot analyses and qRT-PCR analyses of indicated protein (n = 6) and RNA species (n = 11) from whole pancreata of 16-week-old *Ccpg1^GT/GT^* mice, in reference to *Ccpg1^+/+^* controls (± S.E.M., 1-sample t-tests on untransformed values, * = p ≤ 0.05, **** = p ≤ 0.0001, ns = p > 0.05). Representative immunoblot shown in Supp. Fig. 3D.

The biggest increases were seen for REG3A and REG3B, small (<20 kDa) proteins with highly tissue-restricted expression (gut and pancreas), and, strikingly, that have human orthologues upregulated during pancreatic injury (Chen et al., 2019). REG3B is also already known to be required, via an unknown mechanism, for *Kras*-driven ADM (Loncle et al., 2015). Notably, REG3B accumulated partly in a truncated, highly insoluble form (ΔN, Fig. 3B, Supp. Fig. 3C); N-terminal trypsin-processed forms of orthologous REG3 protein isoforms, for example, human REG3γ, represent particularly insoluble, polymeric subspecies (Graf et al., 2001; Mukherjee et al., 2009; Mukherjee et al., 2014). Other proteins accumulating with impaired solubility in *KC Ccpg1^ΔPANC^* pancreata included ZG16, a tissue-specifically expressed lectin-like protein, and AGR2, a protein known to prefigure pre-malignant state changes in the pancreatic acinar epithelium (Dumartin et al., 2017).

We next performed qRT-PCR on cognate whole pancreatic RNA to dissect any transcriptional contribution to the increased overall protein abundances in ER-phagy deficient *KC* pancreata. A subset (*Reg3*a/b, *Agr2*) exhibited transcriptional upregulation in response to *Ccpg1* deficiency whereas others (*Zg16*, *Dnajb9*, *Dnajc3*) did not (Fig. 3D). However, further analyses were then conducted in the absence of oncogenic *Kras* mutation, where upon comparison of protein and RNA abundances it was seen that *Ccpg1* deficiency alone is sufficient to provoke post-transcriptional accumulation and insolubility of all proteins (including the REG3BΔN subspecies), but insufficient to provoke transcriptional responses (Fig. 3E-F, Supp. Fig. 3D-E). Overall, these data show that proteostatic defects due to ER-phagy suppression manifest in impaired turnover and maintenance of solubility of a specific cohort of ER luminal proteins. Importantly, the data also provide a first suggestion that, in the presence of oncogenic *Kras*, primary proteostatic failure may also be exacerbated by feed-forward transcription of genes encoding aggregation-prone proteins.

### REG3B is a physiologically-relevant client of *Ccpg1*-mediated ER-phagy in pancreas

It is unclear how ER-phagy mediates proteostasis in pancreas. It has been proposed that binding of aggregation-prone protein “clients” to the luminal domain(s) of CCPG1 (Supp. Fig. 4A) may facilitate their post-transcriptional clearance via ER-phagy (Ishii et al., 2023). Nonetheless, physiologic client proteins of CCPG1 have not been identified. Given the known importance of REG3B in *Kras-*driven ADM, along with the multi-layered regulation of solubility and abundance of this protein in *Kras-*mutant pancreas, we thus addressed whether REG3 species fulfilled the criteria of physiologic client proteins. Indeed, we found that REG3 family proteins bound to CCPG1, as assayed via co-immunoprecipitation (Fig. 4A). Furthermore, interaction was dependent upon a small, C-terminal helical bundle domain of CCPG1 present within the ER lumen (L2 domain) (Supp. Fig. 4A, Fig. 4B-D). Further underscoring the role of ER-phagy in regulating client protein solubility, we also observed striking accumulation of CCPG1 and REG3A/B together in detergent-insoluble fractions from pancreas deleted for the core autophagy gene *Atg5* (Supp. Fig. 4B, Fig. 4E, *Atg5^ΔPANC^*). Taken together, these data thus strengthen our prior conclusion (Fig. 3) that REG3 proteins are post-transcriptionally regulated by ER-phagy.

**Fig. 4.**
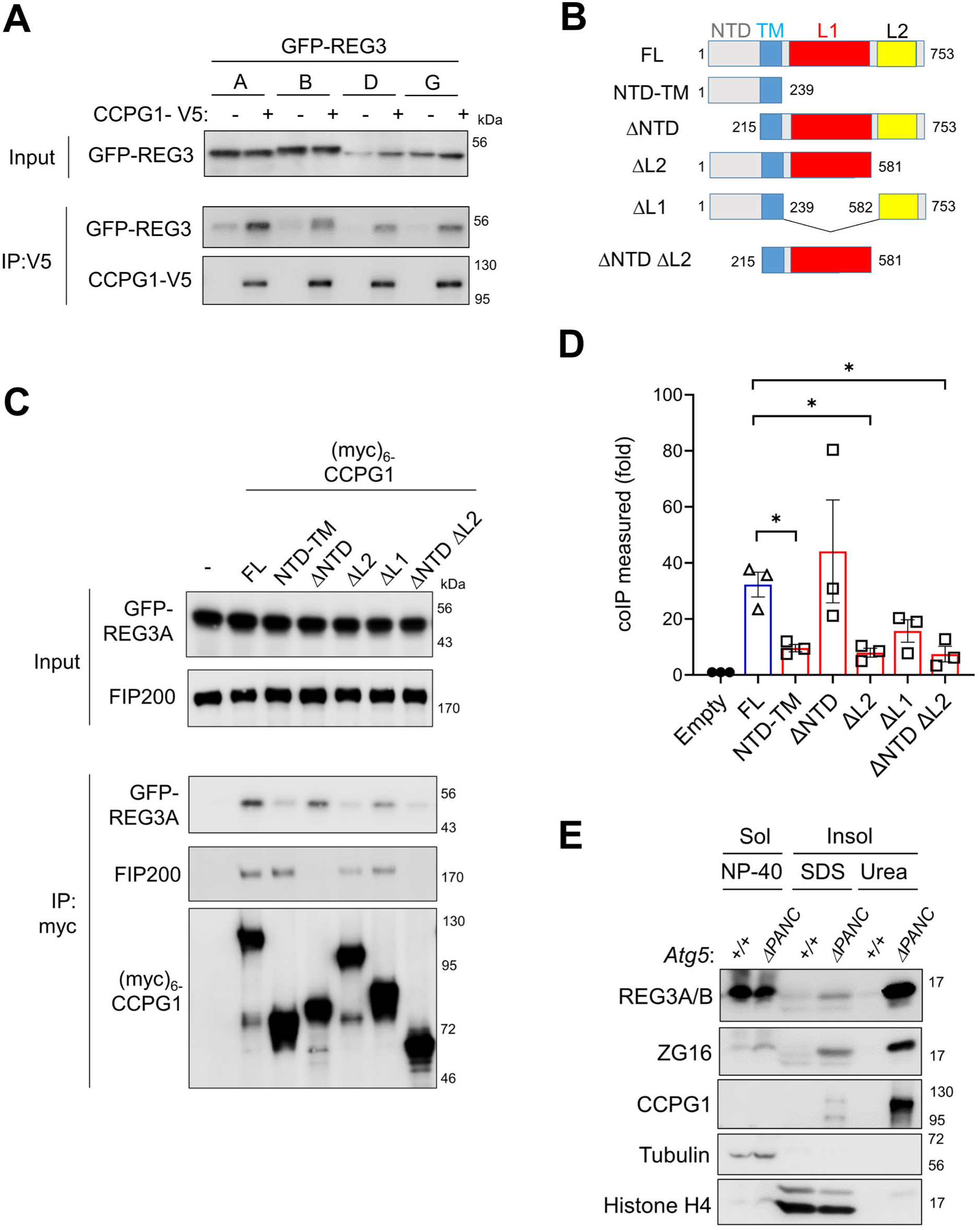
REG3 is a physiologically-relevant client of *Ccpg1*-mediated ER-phagy in pancreas. **A)** Co-immunoprecipitation (IP) of GFP-fused murine REG3-family members (-A, -B, -D, and -G) with C-terminally V5-tagged CCPG1 was assessed by immunoblot. **B)** Full-length (FL) and deletion mutants of murine CCPG1 (NTD = N-terminal domain, TM = transmembrane segment, L1 and L2 = ER luminal domains 1 and 2). **C-D)** HEK293FT cells stably expressing GFP-REG3A were transfected with empty vector (-) or N-terminally myc-tagged FL or deletion mutants of CCPG1, and binding assessed by co-immunoprecipitation, representative immunoblot in **C**, quantification of replicates in **D** (n = 3, 1-way ANOVA and Holm-Šidák post-hoc test of each mutant vs. FL, * = p ≤ 0.05, not shown = p > 0.05). Endogenous FIP200 immunoprecipitation included a positive control for functional FL and NTD-containing deletion mutants of CPCG1. **E)** Immunoblot of whole pancreata from 8-week-old *Atg5^+/+^* (*Pdx1-Cre Atg5^+/+^*) or *Atg5*^Δ^*^PANC^* (*Pdx1-Cre Atg5^flox/flox^*) mice, sequentially extracted in non-denaturing detergent (Sol, NP-40 detergent-soluble), then denaturing detergent (SDS), then Urea (Insol, detergent-insoluble fractions).

### *Kras* drives stochastic proteostatic failure in pancreatic acinar cells

Oncogenic *Kras* itself is sufficient to inhibit ER-phagy function in a stochastic, spatially inhomogeneous manner (Fig. 1). We hypothesised that *Kras* would thus also be sufficient to drive sporadic proteostatic failure. Additionally, our prior hypothesis that ER-phagy defects drive ADM would dictate that a substantial proportion of any such proteostatic defect would be detected proximal to ADM. Indeed, detergent-insoluble forms of REG3A/B, including REG3BΔN, are detected in whole pancreatic extracts from *KC* mice (Supp. Fig. 5A-B). However, this finding provides no spatial information. We thus decided to establish a phenotypic assay for proteostatic failure, first using *KC Ccpg1^ΔPANC^*tissues as a tool, as herein ER-phagy is ubiquitously compromised throughout the pancreatic epithelium. Accumulated REG3A/B was detected in a punctate staining pattern across the entire acinar compartment in such tissue (Fig. 5A-B). Super-resolution microscopy (Fig. 5C) combined with immunogold transmission electron microscopy (Fig. 5D) revealed this staining to arise from focal REG3A/B accumulation within tubulovesicular networks corresponding to the ER lumen. In line with the ER-phagy origin of this phenotype, the same punctate REG3A/B aggregation was detected in *Atg5*- deficient pancreata (Supp. Fig 5C).

**Fig. 5.**
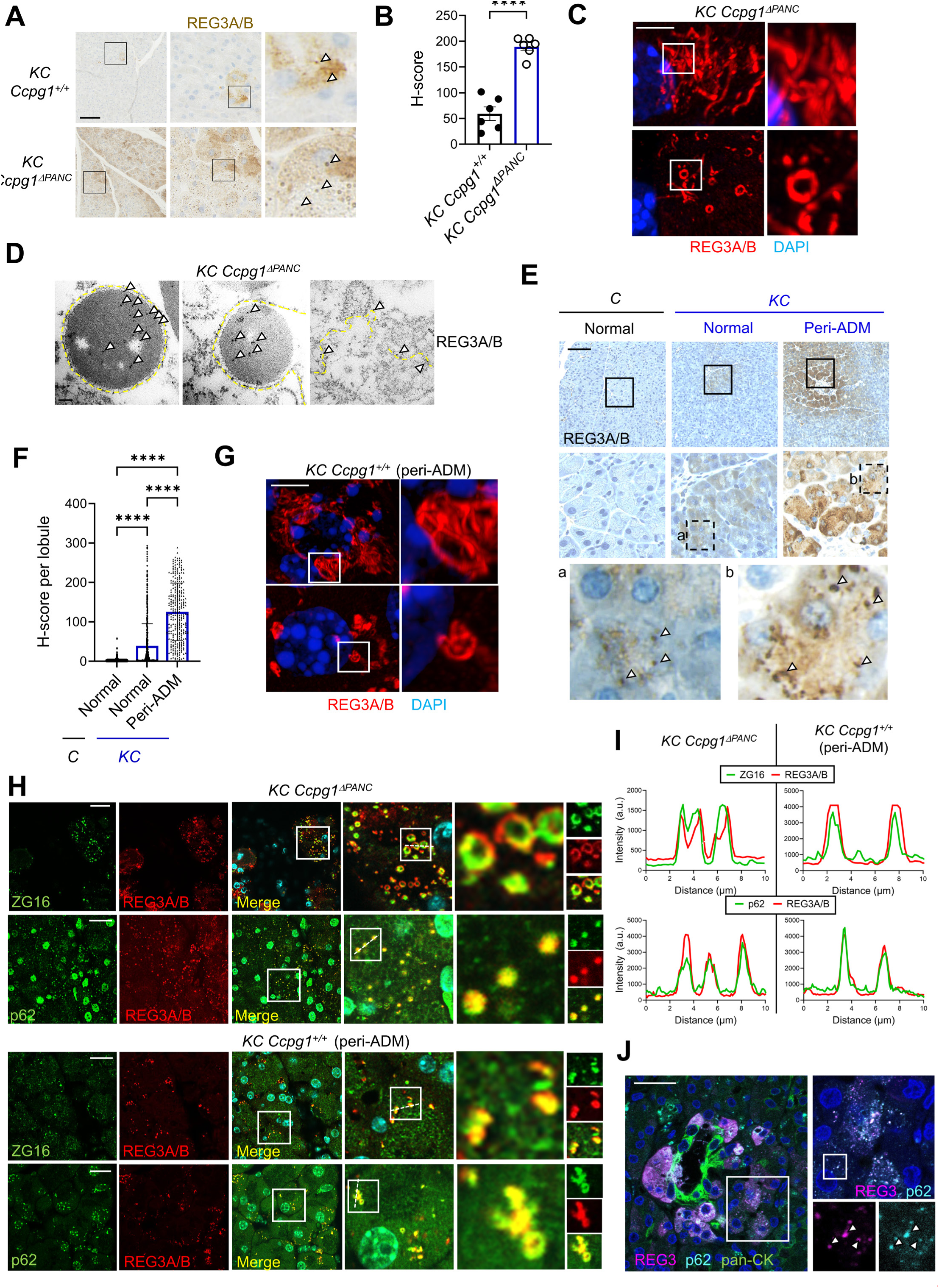
*Kras* drives stochastic proteostatic failure in pancreatic acinar cells. **A-B)** Immunohistochemistry for REG3A/B on formalin-fixed paraffin embedded (FFPE) pancreata from 10-week-old *KC Ccpg1*^Δ^*^PANC^* (*Pdx1-Cre Kras^LSL-G12D/+^ Ccpg1^flox/flox^*) mice, compared with *KC Ccpg1*^+/+^ (*Pdx1-Cre Kras^LSL-G12D/+^*) mice, representative images in **A**, signal quantified in **B** (n = 6, ± S.E.M., Student’s t-test, **** = p ≤ 0.0001). Arrowheads indicate exemplar focal signals. Scale bar = 100 μm. **C)** SORA super-resolution microscopy analysis of REG3A/B in acinar cells within FFPE pancreata of 10-week-old *KC Ccpg1*^Δ^*^PANC^* mice. Scale bar = 5 μm. **D)** Immunogold transmission electron microscopy of REG3A/B in acinar cells within pancreata of 16-week-old *KC Ccpg1*^Δ^*^PANC^* mice. Arrowheads indicate gold particles marking accumulation within intra-ER granular aggregates (left panels) and distributed within the tubular ER network (rightmost panel). Broken yellow lines indicate some of the ER limiting membrane in these images. Scale bar = 100 nm. **E-F)** Immunohistochemistry for REG3A/B on FFPE pancreata from 10-week-old *KC* (*Pdx1-Cre Kras^LSL-G12D/^*^+^) mice, compared with *C* (*Pdx1-Cre Kras^+/+^*) mice. Representative images in **E**, for *KC* pancreata these include fields of acinar cells within both normal lobules and lobules harbouring sporadic ADM (peri-ADM) (see Fig. 1D). Arrowheads indicate exemplar focal signals. Signal quantified in **F** (n = 2487 cells from 10 mice total, ± S.D., 1-way ANOVA and Holm-Šidák post-hoc test, **** = p ≤ 0.0001). Scale bar = 100 μm. **G)** SORA super-resolution microscopy analysis of sporadic acinar REG3A/B signals within FFPE pancreata from 10-week-old *KC* mice. Scale bar = 5 μm. **H-I)** Confocal immunofluorescence microscopy for REG3A/B costaining proteins, ZG16 and p62, in acinar cells within FFPE pancreata from 10-week-old *KC Ccpg1*^Δ^*^PANC^* mice and 18-week-old *KC Ccpg1^+/+^*mice (focussing here on sporadic REG3A/B punctate signals in peri-ADM lobules). Representative images in **H,** signal intensity traces demonstrate colocalization in **I**. Scale bar = 20 μm. **J)** Confocal immunofluorescence microscopy of ADM and surrounding acinar cells in a non-cancerous donor human pancreas (panCK = pan-cytokeratin, here marking ADM). Arrowheads indicate exemplar focally colocalised signals for REG3A/G (human orthologues of REG3A/B) and p62. Scale bar = 15 μm.

Importantly, oncogenic *Kras* was in itself sufficient to engender sporadic, punctate REG3A/B aggregation within acinar cells, as demonstrated by comparison of *KC* pancreatic tissue with that of control *C* mice (Fig. 5E-F). This REG3A/B staining in *KC* mice is detected in both acinar cells within normal lobules and, at much greater frequency, within peri-ADM lobules, i.e., acinar cells in the vicinity of ADM (Fig. 5E-F). This mirrors the spatial pattern of oncogenic *Kras*-driven ER-phagy suppression (Fig. 1D-G). These stochastic aggregates also exhibit the same distinctive tubulovesicular morphology (Fig. 5G) of those induced directly by genetic ER-phagy deficiency (Fig. 5C) and - further underscoring the similarity between the two - in both settings the observed aggregates counterstain for ZG16 (discovered as an ER-phagy target in Fig. 3) and the cytoplasmic early autophagy marker p62 (Fig. 5H-I, Supp. Fig. 5D). Taken together, the above data support two key findings. First, distinctive focal REG3/p62 staining identifies ER-phagy related proteostatic failure in pancreatic acinar cells. Second, oncogenic *Kras* is sufficient to sporadically drive this acinar phenotype, and this is tightly associated with ADM. Importantly, acinar cells surrounding ADM in healthy human pancreata and tumour-adjacent normal tissue also exhibit a REG3/p62 aggregation phenotype, indicating cross-species conservation of the association between proteostatic defects and ductal metaplasia (Fig. 5J, Supp. Fig. 5E).

### ER aggregates mark an intermediate acinar cell state of injury and partial ductal identity

ADM is a cell state generated by transcriptional rewiring of acinar cells. Thus, to gain mechanistic insight, we next aimed to uncover the transcriptional basis of the acinar cell propensity for ADM upon ER-phagy defects. We first analysed the whole pancreatic transcriptome using mRNA-Seq (Supp. Table 2, Supp. Fig. 6A), after homogenous abrogation of ER-phagy in *Kras* mutant animals (*KC Ccpg1^ΔPANC^* versus *KC* control). We initially theorised that the unfolded protein response (UPR) would be engaged, as a canonical reaction to ER dysproteostasis (Hetz, 2012). Surprisingly, no consistent elevation of UPR target gene expression was observed (Supp. Fig. 6B-C). We then alternatively hypothesised that loss of ER-phagy function leads to cellular injury in a small number of acinar cells, which would also be in line with the previous observation of microinflammation (Fig. 2G-H). We analysed recently published transcriptomic data (Del Poggetto et al., 2021; Ma et al., 2022) to compile eight gene sets describing the transcriptional signature of acinar cells upon acute or chronic injury, and during the subsequent acute recovery phase (Supp. Table 3). Gene set enrichment analysis (GSEA) was used to perform an unbiased ranking of these acinar gene signatures versus all molecular signatures in the MSigDB database (Fig. 6A, Supp. Table 3). This revealed enriched mesenchymal, fibroblastic and immunological stromal signatures, in line with more ADM and inflammation upon *Ccpg1* deletion. However, the top-ranking acinar specific gene set was one of the compiled signatures of acutely-resolving injury (INJURY_D7_TOP_UPREGULATED POGGETTO_2021 in red, Fig. 6A, Supp. Fig. 6D). Furthermore, the other seven gene sets signifying current or recent injury were also prominent (in red, Fig. 6A, Supp. Fig. 6D).

**Fig. 6.**
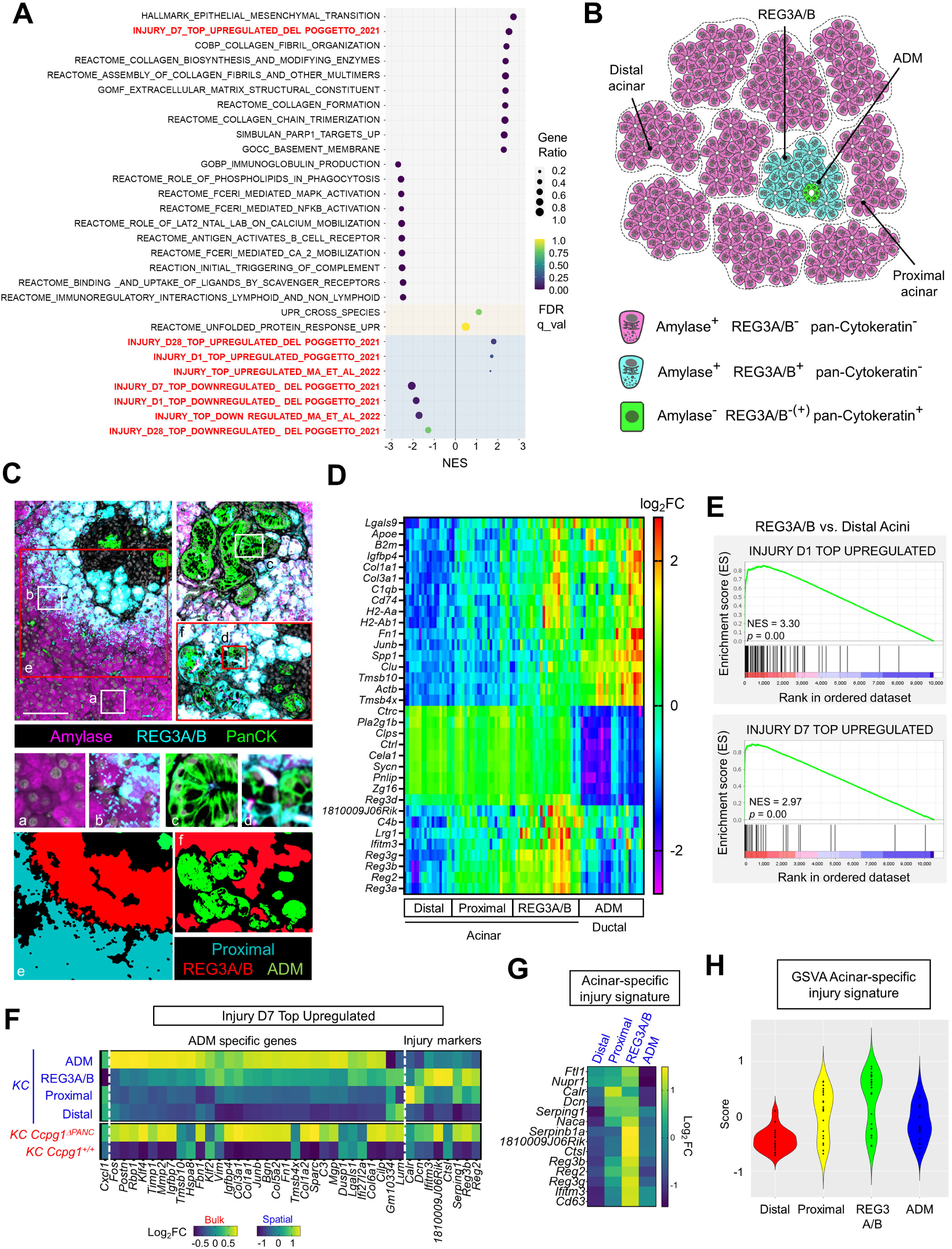
ER aggregates mark an intermediate acinar cell state of injury and part ductal identity. **A)** Gene-set enrichment analysis (GSEA) was performed after whole pancreas mRNA-Seq on 10-week-old *KC Ccpg1^+/+^* (*Pdx1-Cre Kras^LSL-G12D/+^*) or *KC Ccpg1*^Δ^*^PANC^* (*Pdx1-Cre Kras^LSL-G12D/+^ Ccpg1^flox/flox^*) mice. 20 top-ranking signatures are ordered here by NES (normalised enrichment score), and additionally ranked alongside UPR signatures and collated injury signatures (emboldened in red). FDR = false-discovery rate. **B)** Diagram of the 4 different cell populations isolated for spatial transcriptomics from FFPE of 10-week-old *KC* mice **C)** Representative images of immunofluorescence staining of different populations found in vicinity of REG3A/B Acinar cells (top subpanel). *a* is an example of Proximal Acinar, *b* of REG3A/B Acinar, *c* of majority ADM, *d* of rare ADM that also contains some ductal cells exhibiting REG3A/B signal. Bottom subpanel shows some cognate examples (*e* and f) of subsequent segmentation of the different populations for spatial transcriptomics. Scale bar = 100 μm. **D)** Heatmap of top differentially-expressed genes across different cell subpopulations from spatial transcriptomics of *KC* mice. **E)** Example of GSEA using two of the collated injury signatures derived from Del Poggetto et al (2021), comparing REG3A/B^+^ acinar cells with Distal (healthy) acinar cells. For complete scoring of 20 most-enriched gene sets and all of the injury gene sets see the summary in Supp. Fig. 7A. **F)** Heatmap comparing transcript-by-transcript changes within an example injury gene set derived from Del Poggetto et al (2021), both within bulk mRNA-Seq data from wild-type and *Ccpg1-*deficient *KC* mice as in **A** (lower rows, labels in red) and across the different cell populations isolated by spatial transcriptomics from *KC* mice as in **C-E** (upper rows, labels in blue). Further examples for other injury gene sets are provided in Supp. Fig. 7B-D. See text for rationale of subclassification of genes as ADM specific and unique acinar injury markers **G-H)** A 14-gene acinar cell specific injury signature was derived by collating the unique acinar injury markers derived in analyses in **E** and Supp. Fig. 7B-D (excluding ribosomal genes). In **H,** this signature was tested across the cell populations from the spatial transcriptomic analysis using gene-set variation analysis (GSVA).

The above data suggest that genetic loss of ER-phagy provokes a low-level injury state across the *Kras* mutant acinar epithelium. Thus, we reasoned that ER-phagy failure driven more stochastically by oncogenic *Kras* alone would similarly be sufficient to lead to acinar injury. To identify sporadic dysproteostatic acinar cells in this setting, we performed spatial transcriptomics in *KC* mice (Fig. 6B-C, Supp. Table 4). Three morphologically-normal acinar populations were defined via immunophenotyping (all Amylase^+^ pan-Cytokeratin^-^): cells bearing distinctive REG3A/B aggregates, which were frequently observed adjacent to ADM (termed *REG3A/B*); REG3A/B^-^ acinar cells immediately adjacent to the REG3A/B^+^ cells (termed *Proximal*); and control acinar cells (REG3A/B^-^) not in the proximity of ADM or REG3A/B acinar cells (termed *Distal*). A fourth population of ADM itself was identified morphologically and immunologically (termed *ADM*; Amylase^-^ pan-Cytokeratin^+^, predominantly REG3A/B^-^ - a minor population of REG3A/B^+^ ductal cells is detected within a minority of these ADM) (Fig. 6C). As expected, differential gene expression (DGE) analysis revealed transcripts specific to all acinar cells (Fig. 6D, e.g., *Zg16, Pnlip, Cela1*). However, REG3A/B^+^ cells, and, to a lesser extent, Proximal cells, were discriminated from healthy Distal cells by expression of some apparent ADM markers (e.g., *Fn1, Clu, Junb, Col1a1, Spp1, Igfbp4*) as well as unique transcripts that were not enriched for in ADM (e.g., *Reg3a, Reg3b, Ifitm3, 18100009J06Rik*). Furthermore, acinar injury gene sets were strongly enriched in REG3A/B^+^ cells, and, again, less so in Proximal cells, relative to healthy Distal cells (Fig. 6E, Supp. Fig 7A). Detailed examination of expression trajectories of the individual transcripts mapping to these gene sets revealed that injury signatures can be divided into genes uniquely upregulated in REG3A/B^+^ acinar cells (we term “Injury Markers”) and those that are markers of ADM and remain dysregulated upon metaplasia (Fig. 6F, Supp. Fig. 7B-D). This parallels the bifurcating expression trajectories identified in the DGE analysis (Fig. 6D). We collated the REG3A/B^+^ cell- specific injury markers identified within the eight acinar injury gene sets to generate a consolidated “acinar-specific injury signature” (Fig. 6G). This signature was confirmed to be strongly enriched in, specifically, REG3A/B acinar cells by gene set variation analysis (GSVA, Fig. 6H).

In summary, transcriptomic data reveal a non-UPR injury signature that uniquely describes the transcriptional state of cells evidencing the sporadic protein aggregation phenotype downstream of oncogenic *Kras*. This transcriptional state is associated with wider changes that indicate priming for ductal metaplasia, in line with the emergence of ADM from such acinar cells.

### Acinar protein aggregates are causal for injury and ADM

To determine whether protein aggregates generated by ER-phagy deficiency are sufficient to cause ADM, we used rAAV to transduce truncated REG3B (ΔN) - equivalent to the particularly aggregation-prone, proteolytically-processed endogenous subspecies - into *KC* mouse pancreas, on the assumption that ER-phagy would be insufficient to prevent aggregation. We found that REG3B FL (full-length) control does not accumulate within pancreatic tissue upon expression (Fig. 7A-C) but instead is detected in plasma, indicative of secretion (Fig. 7D, Supp. Fig. 8A). In stark contrast, REG3B protein levels do increase in pancreatic tissue upon expression of REG3BΔN, partly in a detergent-insoluble fraction indicative of aggregation, but no secretion is detectable (Fig. 7A-D, Supp. Fig. 8A). The aggregation-prone, secretion- disabled nature of REG3BΔN was corroborated *in vitro*, using transfection of HEK293FT cells (Supp. Fig. 8B-C). Importantly, the REG3B accumulated in acinar cells upon transduction with REG3BΔN is detected by immunofluorescence within distinct p62-positive inclusions (Fig. 7E-F). Thus REG3BΔN, but not REG3B FL, drives protein aggregation formation.

**Fig. 7.**
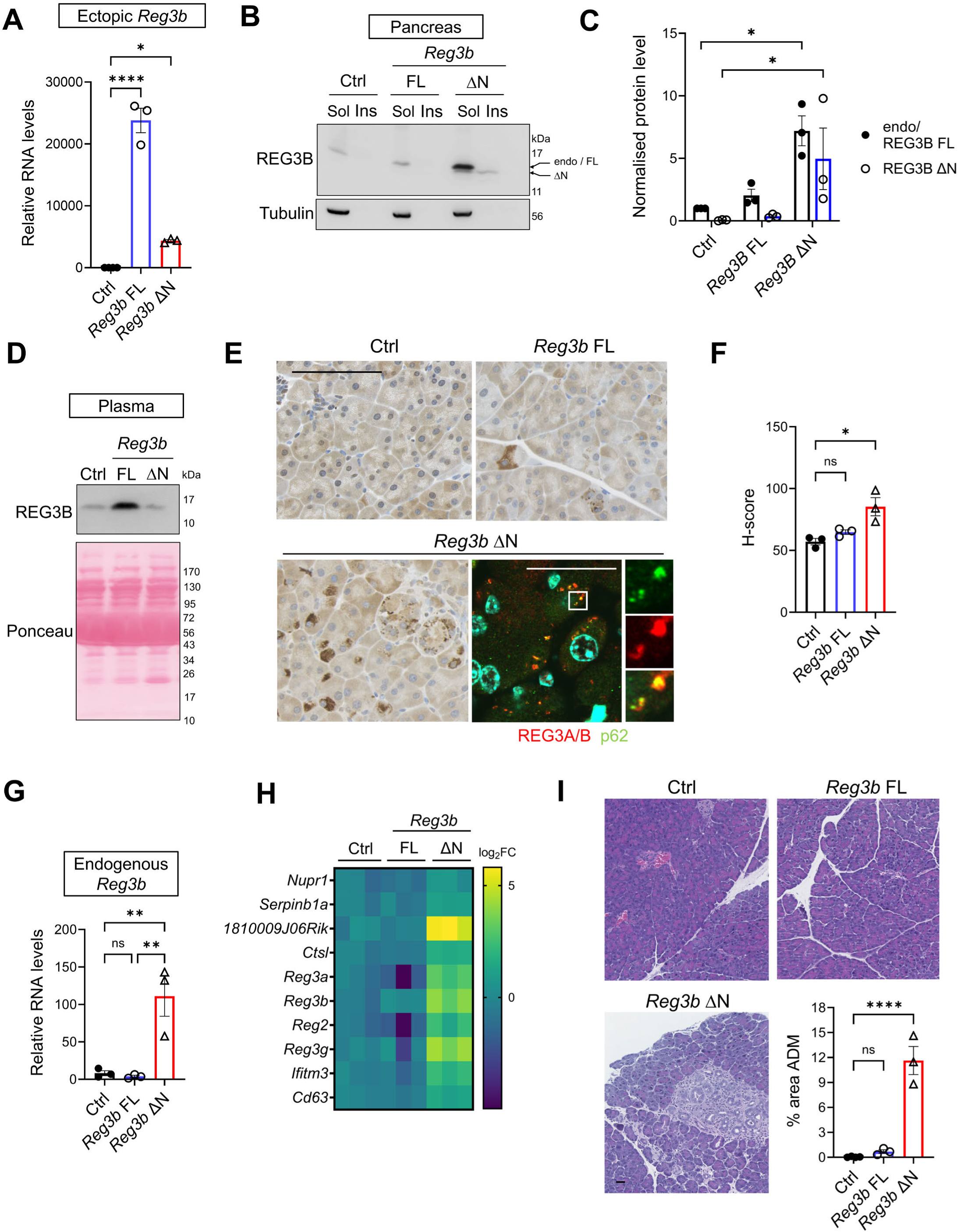
Acinar protein aggregates are causal for injury and ADM. 5-week-old *KC* mice were transduced with control rAAV (Ctrl) expressing luciferase or ectopic *Reg3b* forms (n = 3, FL = full-length, ΔN = N-terminally processed REG3B) then pancreatic tissue and plasma analysed at 10 weeks of age. **A)** qRT-PCR of whole pancreatic RNA to assesses expression of ectopic *Reg3b* (n = 3, ± S.E.M., 1-way ANOVA and Holm-Šidák post-hoc tests versus Ctrl, **** = p ≤ 0.0001, * = p ≤ 0.05). **B-C)** Immunoblotting for endogenous (endo) and ectopic (FL and ΔN) REG3B in whole pancreatic detergent (NP-40)-soluble (sol) and-insoluble (insol) extracts. **B** shows a representative immunoblot, **C** quantifications (n = 3, normalised to endo/FL REG3B in Ctrl group, ± S.E.M., 1-sample t-tests for endo/FL vs. Ctrl, Student’s t-tests on log fold changes for ΔN vs. Ctrl, * = p ≤ 0.05, not shown = p > 0.05). **D)** Representative immunoblotting of REG3B in plasma, quantified in Supp. Fig. 8A. **E-F)** Immunohistochemistry (IHC) of REG3B signal in FFPE, representative images in **E** (lower right panel: representativ confocal immunofluorescence colocalization of punctate REG3 signal with p62 in *Reg3b*ΔN transduced mice). IHC quantifications in **F** (n = 3, ± S.E.M., 1-way ANOVA and Holm-Šidák post-hoc tests vs. Ctrl, * = p ≤ 0.05, ns = p > 0.05). **G)** qRT-PCR of whole pancreatic RNA to assesses expression of endogenous *Reg3b* (n = 3, ± S.E.M., 1-way ANOVA and Holm-Šidák post-hoc test, ** = p ≤ 0.01, ns = p > 0.05). **H)** Heatmap summary of qRT-PCR of whole pancreatic RNA for a subset acinar cell injury signature transcripts (n = 3). **I)** Representative H & E images quantified for ADM (n =3, ± S.E.M., 1-way ANOVA and Holm-Šidák post-hoc tests vs. Ctrl, **** = p ≤ 0.0001, ns = p > 0.05). Scale bars = 30 μm.

We next addressed the direct sequelae of protein aggregate formation. First, we noted that the increased signal for REG3B in pancreata partly reflected increased expression of endogenous (full-length, FL) REG3B, in addition to the transduced REG3BΔN (Fig. 7B-C). qRT-PCR analysis demonstrated that this was due to upregulation of endogenous *Reg3b* transcription (Fig. 7G). This observation is in line with the previous proposal of feed-forward transcription of genes encoding aggregation-prone proteins driven by initial loss of proteostasis in combination with oncogenic *Kras* (Fig. 3C-F). Importantly, expression of REG3BΔN drove the acinar-specific injury signature (Fig. 7H), as well as microinflammation (Supp. Fig. 8D) and ADM (Fig. 7I). Taken together, the above data show that protein aggregates seeded by an insoluble client of ER-phagy are indeed sufficient to drive ADM.

Notably, a prior study has demonstrated that *Reg3b* knockout prevents *Kras*-driven ADM and PanIN in the *KC* mouse model (Loncle et al., 2015). The biological function of Reg3 family proteins is not well understood, with the exception of convincing data that their aggregation-prone nature permits them to polymerise and encapsulate bacteria during host defence responses (Mukherjee et al., 2014). Nevertheless, in the context of pancreatic cancer initiation, it was speculated that REG3B might have a role as a cytokine, triggering JAK/STAT3 or MAP kinase (ERK1/2) signalling (Loncle et al., 2015; Zhang et al., 2021). However, it is the non-secreted, aggregating form of REG3B that is sufficient here to co-operate with *Kras* in promoting ADM. Importantly, this does not preclude a role for the feed-forward upregulation of endogenous *Reg3* isoform expression, resulting in secreted growth factor being produced. Indeed, reconciling these observations, we find that the insoluble, non-secreted form of REG3B (REG3BΔN) does in fact provoke JAK/STAT3 activation at a tissue level, in contrast to the full-length, secretion-competent control (Supp. Fig 8E-F).

Taken together, the above data show that protein aggregation is sufficient to account for sporadic injury and ADM triggered by ER-phagy deficiency in acinar cells bearing oncogenic *Kras*.

## Discussion

Collectively, our data support the following model for early pancreatic tumorigenesis (Fig. 8). Sporadic failure of ER-phagy in acinar cells leads to intraluminal protein aggregation. Aggregates co-operate with oncogenic *Kras* to trigger localised tissue injury, transcriptional state changes and microinflammation. Our results show that cancer joins the grouping of serious diseases that are promoted by longitudinal dysproteostasis and protein aggregation, alongside neurodegenerative disorders and myopathies (Klaips et al., 2018).

**Fig. 8.**
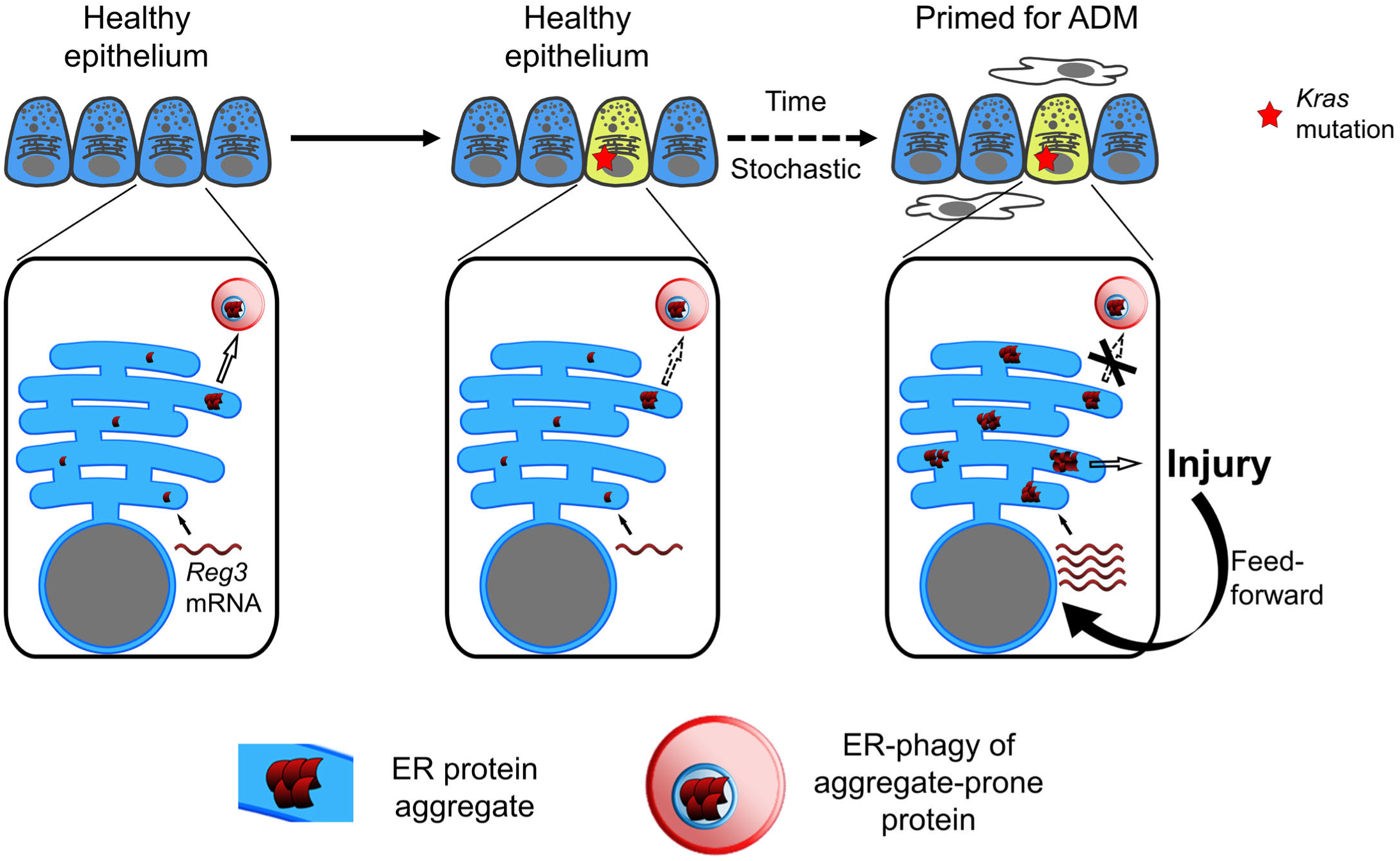
Model of acinar dysproteostasis underpinning early *Kras-*driven tumorigenesis. See text in Discussion for details.

Our findings also reveal a new function for *Kras* in tumorigenesis. It has been firmly established that oncogenic *Kras*-mediated transcription drives the cell-cycle (Klomp et al., 2024) and consequent senescence (Morton et al., 2010), as well as engaging general autophagy for cancer cell survival. Notably, recent studies have established that oncogenic *Kras* “locks in” the transcriptional changes underpinning ADM that occur upon experimental injury (Alonso-Curbelo et al., 2021; Li et al., 2021; Burdziak et al., 2023). However, transcriptional events before this, in pre-dysplastic epithelia, have been underexplored. We show that oncogenic *Kras* transcriptionally inhibits ER-phagy in pancreatic acinar cells and that the probability of sporadic proteostatic failure is accordingly elevated. Thus, *Kras* not only co-operates with proteostatic failure in generating irreversible ADM and cancer, but greatly elevates the likelihood of dysproteostasis in the first instance. This observation highlights an unexpected dual nature of the relationship between oncogene action and proteostatic failure.

Beyond early transcriptional changes yet prior to ADM, *Kras*-induced proteostatic defects also define a sporadic pathologic acinar cell state associated with acinar-specific injury transcripts and partial ductal identity. Notably, rare acinar cells with decreased acinar identity and/or characterised by *Reg3*-family member expression have been observed as minority populations within recent single human and mouse cell transcriptomic datasets, and in some of these studies were detected at greater frequency after oncogenic *Kras* mutation or inflammation (Muraro et al., 2016; Schlesinger et al., 2020; Gopalan et al., 2021; Tosti et al., 2021; Chondronasiou et al., 2022; Cui Zhou et al., 2022; Ma et al., 2022). This suggests the existence of conserved core features marking minority acinar cell states that - in the case of proteostatic failure as described here, at least - facilitate ADM. From a mechanistic standpoint, we reveal that REG3B protein, along with a select cohort of other ER luminal proteins, forms injurious aggregates, driving the distinctive transcriptional changes observed in these cells. In the case of aggregated REG3 proteins, we also identify the origin of dysregulated proteostasis; REG3 proteins are revealed here as the first physiologic “client” proteins for pancreatic ER-phagy, i.e., targets of autophagic degradation via interaction in the ER lumen with the CCPG1 ER-phagy receptor. Importantly, molecular changes triggered by failure to prevent protein aggregates containing REG3 proteins also include feed-forward increases in *Reg3*-family transcription, in line with the identification of these transcripts as signature features of the pathologic acinar cell state. This probable dysproteostasis amplification loop appears to occur specifically when *Kras* is mutated, again underscoring the dual nature of the relationship between oncogenic *Kras* and proteostasis.

*KRAS^G12D^* mutations accumulate in the human acinar epithelium with age (Neuhofer et al., 2021), potentially underpinning longitudinal accumulation of premalignant PanIN (Braxton et al., 2024). Building a molecular mechanistic description of cancer risk, framed around co-operating events that trigger step-wise progression toward PDAC, will facilitate future preventative strategies. In this regard, this work suggests future questions at varying scales that should be addressed to understand pancreatic acinar proteostasis (or lack thereof) during ageing. These include: detailed mechanistic description of ER-phagy regulation and molecular mechanism; the role of background genetics and environmental factors in modifying proteostatic function; and the role of ageing in modifying the proteostatic outputs of oncogenic *Kras*. Similarly, interventions that ameliorate proteostatic failure - as proposed for age-related conditions such as neurodegeneration (Klaips et al., 2018; Aman et al., 2021; Lee et al., 2022) - should be investigated for their impact on cancer development.

### Limitations of this study

The *KC* mouse and variants thereof are widely-used model for early pancreatic tumorigenesis from acinar cells, particularly ADM and PanIN. However, other routes to PDAC including ductal cell-of-origin and non-ADM pre-malignant lesions - such as intraductal papillary mucinous neoplasia – are not possible to interrogate using the KC mouse model. While beyond the scope of this study, they may also be influenced by the proteostatic health of the acinar epithelium. This should be investigated in future. Additionally, while acinar proteostatic failure is detected in humans and does associate with ADM, intervention studies cannot currently be performed here. Thus, causality remains inferred from the mouse model.

## Author contributions

Conceptualization, SW, NJM, MDM and CSC; Methodology, SW, MS and CSC; Software, AvK, RK and CSC; Validation, CSC, MS, JH, JM; Formal Analysis, AvK and CSC; Investigation, CSC, MS, MDM, JH, RK and NJM; Resources, SW, DM, XZ, JM, MM; Data Curation, AvK, SW, and CSC; Writing – Original Draft Preparation, SW; Writing – Review & Editing, SW and CSC; Visualization, SW and CSC; Supervision, SW, NJM; Project Administration, SW; Funding Acquisition, SW.

## Declaration of interests

The authors declare no competing interests.

## Acknowledgements

We thank Valerie J. Brunton and Ian Jackson (Institute of Genetics and Cancer, University of Edinburgh) for the gift of *KC* and *KPC* mouse strains, and Deleter mouse strains, respectively. We thank the Biomedical Research Facility (University of Edinburgh) for additional assistance with mouse husbandry. We thank Charles David (Tsinghua University) for the gift of the pAAV2/8 vector. We thank Steven Mitchell (Wellcome Centre for Cell Biology, University of Edinburgh) for electron microscopy assistance. We acknowledge Alison Munro of HTPU Microarray Services at the Institute of Genetics and Cancer, University of Edinburgh, for their technical support with the NanoString GeoMx Digital Spatial Profiler. We thank Matt Pearson and Laura Murphy from the Advanced Imaging Resource, Institute of Genetics and Cancer, University of Edinburgh for technical support. SW, NJM, CSC and KWI were funded by a Senior Cancer Research UK Fellowship awarded to SW (A29576). AJK was funded by a Cancer Research UK Career Development Fellowship awarded to SW (A12825). JH was funded by a Wellcome Trust clinical academic track scholarship. MS was funded by a BBSRC project grant (BB/N000315/1). JPM is funded by CRUK core funding to the CRUK Scotland Institute (A17196 and A31287), to JPM (A29996), and by a CRUK Precision-Panc grant (A25233). MM is funded by a CRUK Centre award (CRUK Scotland Centre; CTRQQR-2021\100006). DM was funded by a UKRI MRC Senior Clinical Fellowship (MR/P008887/1). The Dragonfly spinning-disk confocal was funded by the MRC (MC_PC_MR/X012212/1). SORA microscope was funded by an MRC Capital Equipment Award. The Orbitrap Fusion Lumos mass spectrometer was funded by the Wellcome Trust (Multiuser Equipment Grant 208402/Z/17/Z). Electron microscopy was supported by the Wellcome Trust (Multi User Equipment Grant WT104915MA).

## Methods

### Tissue culture

Cells were cultured in Dulbecco’s modified Eagle’s medium (DMEM), supplemented with 10% Fetal Bovine Serum (FBS), 1% L-Glutamine and 0.1% Penicillin/Streptomycin at 37 °C in 5 % CO_2_. HEK293FT were from laboratory stocks and AAVpro 293T were from Takara Bio (#623273). Murine PDAC (pancreatic ductal adenocarcinoma) cells were an established line generated from a tumour dissected from the KPC mouse model on a C57/BL6J pure background. To make REG3-GFP isoform and CCPG1-V5 expressing cell derivatives, HEK293FT cells were used to package lentivirus, via transfection in 6 cm dishes in antibiotic-free culture medium, using 2 μg plasmid DNA complexed with Lipofectamine 2000 (Invitrogen 11668019) in Opti-MEM (Gibco 31985062), as per manufacturer’s instructions (1:0.75:0.25 ratio lentivirus: psPAX2: pMD2.G). Lentivirus was collected in fresh medium from 24 h – 48 h post-transfection, 0.45 μm filtered, and used to infect target cells for 24 h in 8 μg/ml polybrene, prior to selection in 10 μg/ml blasticidin (Fisher BP264725). All cell lines were routinely screened for mycoplasma contamination. Human cell lines were confirmed by microsatellite genotyping.

### Mouse strains and breeding

All animal work was approved by a University of Edinburgh internal ethics committee and was performed in accordance with institutional guidelines under license from the UK Home Office (PP7280430 and PP7510272). Specifically, mice were bred in i.v.c cages at a constant ambient temperature of 21 °C with 12 h light-dark cycles. Pathogen status was monitored by environmental interceptors and sentinel testing. All genotyping was outsourced to Transnetyx (Memphis, USA). Individual genetically-altered alleles used in this study that were: *Pdx1-Cre (*MGI:3032531); *Ccpg1* gene-trap (*Ccpg1*^GT^ MGI:5000356 also denoted as *Ccpg1^tm1a^*); conditional oncogenic *Kras (Kras^LSL-G12D^* MGI:2429948); conditional mutant p53 (*Tp53^LSL-R172H^*^;^MGI:3039263) and conditional deletion *Atg5* (*Atg5^flox^* MGI: 3663625). Additionally, *Ccpg1^flox^*alleles (*Ccpg1^tm1c^*) were generated by breeding *Ccpg1^GT/GT^*parental animals with CAG-FLPe (FLPe deleter) animals (MGI: 3850329) and then selection of germline FLPe^-^ *Ccpg1^flox/+^* mice from within F2 offspring. Compound genetically-altered mice - *KC* and *KPC* - were generated by interbreeding of *Pdx1-Cre, Kras^LSL-G12D^* and *Tp53^LSL-R172H^*mice. ER-phagy and autophagy-deficient mice for experimentation were generated by breeding *KC* and *KPC* mice heterozygous for *Ccpg1* gene-trap or flox alleles, or *Atg5^flox^*, to generate sibling *K(P)C* animals both wild-type and homozygous for the cognate allele. In general, pairs of comparator siblings of both sexes were randomly selected from litters to populate data sets, allowing pair-wise normalisation of quantifications to control animals to minimise variance, where appropriate. For Kaplan-Meier survival analyses, mice were aged until symptoms of pancreatic lesion-associated morbidity were evident, whereupon they were euthanised by cervical dislocation and pancreas examined at necropsy to confirm morbidity due to evident PDAC (for *KPC* mice) or superabundant PanIN (*KC* mice). With the exception of experiments where plasma was harvested, mice were terminated by cervical dislocation. Dissected pancreata (head and tail together) were either processed for RNA or protein extraction, or fixed by immediate immersion in 10% neutral buffered formalin and incubation overnight at RT. The following day samples were washed 2 x 70% EtOH and embedded in paraffin.

### Protein extraction from pancreas

PBS-washed pancreas was snap frozen in liquid nitrogen and protein extracted using a modification of (Smith et al., 2022). Briefly, frozen pancreas was wrapped in tinfoil and pulverised under liquid nitrogen. Powdered pancreas was then lysed in 400 μl - per 50 mg of initial mass - of extraction buffer, either NP-40 buffer (50 mM Tris-HCl, pH 7.5, 150 mM NaCl, 0.5% IGEPAL, 2 mM activated Na_3_VO_4_, 20 mM NaF, 10 mM sodium pyrophosphate, 1 mM PMSF and cOmplete protease inhibitor cocktail, Roche, 11697498001) or SDS lysis buffer (4% SDS, 150 mM NaCl, 50 mM Tris-HCl pH 7.5), depending upon experiment. NP-40 lysates were homogenised with a Dounce homogeniser, transferred to a microcentrifuge tube, left on ice for 5 min and sonicated for a few seconds. After 10 mins on ice, NP-40 lysates were then centrifuged at 17000 x g for 15 min at 4 °C, supernatant collected as NP-40-soluble and pellets were washed with NP40 buffer and resolubilised in SDS lysis buffer (400 μl per 50 mg of initial weight). SDS Lysates were vortexed, homogenised with a 21G needle, heated at 98 °C for 5 min and sonicated for a few seconds. Samples were centrifuged at 17000 x g for 15 min and supernatant was collected. Initial-detergent insoluble pellets were washed in the same amount of initial buffer and then further extracted, either in SDS lysis buffer, as described above, or, for SDS-insoluble pellets, solubilised in 400 μl - per 50 mg of initial mass - of urea lysis buffer (8 M Urea, 1% SDS, 10 mM DTT, 10 mM Tris-HCl pH 8.0). Samples were vortexed, boiled for 2 min and sonicated briefly. Protein concentration was measured either using a Nanodrop 2000 Spectrophotometer (SDS lysates, Thermo Fisher Scientific) or BCA protein assay (NP-40 lysates, Thermofisher, 23225).

### Immunoblotting

Samples were denatured at 98 °C in Laemmli buffer for 5 min, then separated by electrophoresis on 4-20% polyacrylamide Tris-Glycine gels (Novex, Invitrogen), or, for ER-phagy receptors, 3-8 % Tris-Acetate gels (NuPAGE, Invitrogen). For whole cell and pancreas lysates, 30 μg of detergent-soluble extract and equivalent volumes of other fractions were typically loaded. Proteins were transferred to 0.45 µm nitrocellulose membranes by wet-transfer, washed in TBST (TBS, 0.05% Tween-20) and incubated in 5% BSA in TBST for 1 hour (or 5% non-fat dry milk in TBST for CCPG1 probing). Membranes were then incubated with primary antibody diluted in 2.5 % BSA in TBST overnight (or 5% non-fat dry milk for CCPG1). Membranes were washed 3 x TBST for 10 min and incubated with HRP-conjugated secondary antibody diluted 1:4000 (Cell Signalling Technology). Membranes were then washed 3 x TBST for 10 min and 1 x PBS. Enhanced chemiluminescence was used to detect HRP, typically visualising on an Amersham ImageQuant 800 imaging system (Cytiva). Quantitative analyses were performed using ImageJ 2.9.0 (Gel analyzer plugin, NIH, Bethesda, USA).

### RNA extraction from pancreas

Pancreata were diced into RNAlater (Sigma-Aldrich, R0901) and stored at 4 °C for 2 h prior to snap freezing in liquid nitrogen. Subsequent RNA extraction was performed as described in (Smith et al., 2022). Briefly, frozen pancreas was wrapped in tinfoil and pulverised under liquid nitrogen, the resultant powder was homogenised using a Dounce homogenizer containing 1 ml of TRIzol (Invitrogen, 15596026) per 50 mg of initial pancreas mass. Samples were then clarified by centrifugation at 12 000 x g for 10 min at 4 °C. 200 μl of chloroform was added per 1 ml of TRIzol and samples were slowly vortexed to homogeneity, incubated at RT for 2 min then centrifuged at 12 000 x g for 15 min at 4 °C. The aqueous phase was precipitated in isopropanol and pellets resuspended in RLT buffer for further purification by the Qiagen RNeasy kit (Qiagen, 74106), as per manufacturer’s instructions. RNA concentration was measured using a Nanodrop 2000 Spectrophotometer (Thermo Fisher Scientific). RNA quality and integrity was assessed by RNA ScreenTape analysis in an Agilent 4200 TapeStation system (Agilent).

### qRT-PCR

1st strand cDNA was synthesised from total RNA using qScript cDNA SuperMix (Quantabio) and random priming, according to manufacturer’s instructions. qPCR was performed on an Applied Biosystems StepOne Plus Real-Time PCR System using SYBR green select master mix (Applied Biosystems) as per manufacturer’s protocol. Normalisation was performed to *18S rRNA* or *Gapdh* mRNA by standard curve method or ΔΔCt method, respectively.

### Recombinant adeno-associated virus generation

Adeno-associated virus (AAV) serotype 2/8 was prepared as previously described (Chen et al., 2017; Li et al., 2021). Briefly, AAVpro 293T cells in 15 cm dishes were transfected using Opti-MEM (Gibco 31985062) and PEI-Max (Polysciences 24765-1), at a ratio of 1:1:1 AAV plasmid, pAAV2/8 (gift from C. David, Tsinghua University) and pHelper (a gift from L.Boulter, University of Edinburgh). Viral particles were collected from culture medium and cell lysates. Briefly, cells were scraped into 5 ml lysis buffer (10 mM Tris-HCl pH 8.5, 150 mM NaCl) and then freeze-thawed 3 times in dry ice, alternating with heating at 37 °C, then centrifuged at 3,000 x g for 10 min. The resultant supernatant lysate was combined with original cell culture medium. 1 vol of concentrator solution was then added (40 % PEG-8000, 1.2 M NaCl, PBS pH 7.4) to 3 vol of viral particles and rocked overnight at 4 °C, then centrifuged at 1600 x g for 60 min at 4 °C. The resultant viral pellet was resuspended in 1 ml cold PBS, incubated overnight at 4 °C and then centrifuged at 13,000 g for 3 min at 4 °C. The final AAV-containing supernatant was titered as described using qPCR (Aurnhammer et al., 2012), then aliquoted and frozen prior to use.

### ER-phagy reporter mice

Delivery of rAAV expressing ss-Ypet-TOLLES-KDEL to mouse pancreas was performed via a single intraperitoneal (i.p.) injection of neat virus preparation (5 ml/kg of 5 x 10^7^ viral genome / μL). Fluorescence was preserved upon tissue fixation via adaptation of a previously described method (McWilliams and Ganley, 2019). Briefly, 2 weeks post injection with rAAV expressing ss-YPet-TOLLES-KDEL, perfusion fixation was performed with freshly-prepared 3.7% paraformaldehyde in 200 mM HEPES pH 7.0. The pancreas was post-fixed in the dark for 24 h in fresh paraformaldehyde solution. Tissue was subsequently cryoprotected by washing 3 x cold PBS followed by immersion into 30% sucrose in PBS at 4 °C overnight. Tissue was blotted to remove excess fluid then placed into circular tinfoil moulds and covered completely with O.C.T mounting medium (VWR, Leicestershire, UK) and equilibrated for 15 mins at RT. Moulds were then floated on liquid nitrogen until frozen, followed by submersion for 2 mins. Frozen tissue blocks were stored at −70 °C then cryosectioned.

### ER-phagy reporter microscopy

Cryosections were thawed at RT for 1 h, washed in PBS, incubated with DAPI in PBS for 10 mins, then washed in PBS. In some instances, additional immunostaining was performed at this step (in the absence of Triton-X-100 treatment, which ablates TOLLES fluorescence). Briefly, tissue was blocked for 1 h in PBS + 1 % BSA, and then incubated in primary antibody (1:50) in fresh block solution for 1 h at RT, then washed 3 x PBS, then incubated in Alexa Fluor 647 conjugated secondary antibody (Invitrogen) in block solution for 1 h. Coverslips were mounted onto slides using DAKO mounting medium (CS70330-2, Agilent Technologies) and sealed with clear nail varnish. Fluorescence imaging of z-stacks was performed on a spinning-disk confocal microscope (Andor Dragonfly+ Mosaic) equipped with a 20x (Plan Apo VC 0.75 N.A) and 60x objective lens (Plan Apo VC 1.4 N.A), HiRes camera and z drive. TOLLES was excited by a 445-nm diode laser and fluorescence was detected within the wavelength range of 460–500 nm. YPet was excited by a 514-nm diode laser and fluorescence was detected within the wavelength range of 530–580 nm. Where additional staining was performed, Alexa Fluor 647 was excited by a 647-nm diode laser and fluorescence was detected within the wavelength range of 651-667 nm.

### ER-phagy reporter analysis

All images were processed with Fiji v2.1.0 software (ImageJ, NIH). Quantification of TOLLES- only foci was carried out with the automated mito-QC Counter tool (Montava-Garriga et al., 2020), with the following adjustments made when merging stacked images on ImageJ: the “red channel” was assigned to CFP/Channel1 (TOLLES), the “green channel” to YPet/Channel 2 and the “blue channel” used for DAPI/Channel 0. Z-projections were generated using a standard deviation projection, with all AAV-infected acinar cells manually selected as regions of interest (ROIs) within each lobule of acinar cells, or, for comparison of *Ccpg1*^GT/GT^ mice, simply within each field of view. Per lobule or per field ER-phagy indices were yielded by division of the sum of TOLLES-only foci across all ROIs in the lobule/field by the total number of ROIs, thus yielding the mean number of TOLLES-only puncta per cell for that lobule/field. In the supplementary quantification method, focussing upon determining relative abundances of YPet^+^ TOLLES^+^ foci and YPet^-^ TOLLES^+^ foci - in order to interrogate the initiation versus completion stages of autophagy - Z-stacks were reconstructed into multi-channel 3D models by Imaris software (Oxford Instruments, v10.0). Acinar cells were designated as ROIs using the Surface creation tool, and the Spots creation tool was used to detect fluorescent foci (settings: “Object – Object Statistics”, estimated XY diameter 1 μm, with consistent background subtraction across all images), and using the Imaris XTension: Spot colocalization tool, using a centroid displacement of < 1 μm.

### Mouse pathology

Formalin-fixed pancreata were cut into pieces at both head and tail and these were distributed around a mould before embedding in paraffin. 4 μm sections were then cut and stained with haematoxylin and eosin. Stained slides were imaged using a Nanozoomer slide scanner (Hamamatsu Photonics) and viewed using NDP.view2 software. ADM (isolated and within tubular complexes) and PanINs were quantified manually by defining areas occupied by the lesions, excluding any surrounding inflammation not core to the lesion (Gopinathan et al., 2015). The area of these was expressed as a quotient of total pancreatic area (normal and lesion, excluding large regions of inflammatory cells occupying large interlobular spaces).

### Immunohistochemistry

4 μm sections were cut from formalin-fixed, paraffin-embedded samples (FFPE) onto charged slides then placed in a 37°C oven overnight. Slides were de-waxed in xylene, then rehydrated with graded ethanol. Antigen retrieval was performed by immersing the slides in 10 mM sodium citrate, pH 6.0 and boiling them for 10 min using a microwave, or, for F4/80 staining, via incubation in proteinase K at RT for 10 min (20ug/ml in 500mM Tris-HCl pH 7.5, 1mM EDTA pH 8.0, 5mM CaCl2, 0.5% Triton X-100 pH 7.5). Slides were quenched with 3% H_2_O_2(aq)_ and underwent endogenous biotin blocking with avidin and biotin (Biolegend, SIG-31126), with 3 x PBS washes for 5 min inbetween. Slides were then washed 3 x PBS for 5 min and then protein blocked for 30 mins in PBS, 0.5% casein, 0.5% BSA, 0.1% sodium azide (Abcam, ab64226). In the case of mouse primary antibodies, slides were incubated with Mouse IgG Blocking Reagent for 1 h, washed in PBS and then equilibrated in mouse-on-mouse (MOM) diluent (MOM staining kit, Vector laboratories BMK-2202) for 5 mins. Slides were then incubated with primary antibody solution, typically diluted 1:50-1:400 depending on antibody, in 1 % BSA / PBS or MOM diluent, overnight at 4°C. Slides were washed 3 x PBS for 5 min and biotinylated secondary antibody (Biotinylated goat anti-rat/mouse/rabbit IgG, Vector Labs) diluted 1:400 in block (Abcam, ab64226) was added and incubated for 30 min before 3 x 5 min PBS washes. Streptavidin-HRP conjugate (VECTASTAIN Elite ABC reagent, Vector labs, PK-7100) was then added to each sample for 30 min at RT before 3 x PBS washes. 3,3’-Diaminobenzidine (DAB) solution (Abcam, ab64238) was applied for a maximum of 10 min. Counterstaining was performed with Harris Haematoxylin with bluing in saturated lithium carbonate solution. Samples were washed with running water and then dehydrated in graded ethanol solutions (30%, 50%, 80%) for 1 min each and then incubated 2 x 5 min in 100% ethanol before 1 min of xylene twice. Coverslips were then mounted using DPX (CellPath, SEA-1304-00A). Slides were then scanned using a Nanozoomer slide scanner (Hamamatsu Photonics) and viewed using NDP.view 2 software. Quantifications of macrophage abundance were performed using the positive cell detection function in QuPath software v0.5.0 using predefined RGB parameters for DAB detection. Positive cells were normalised to pancreatic area (mm^2^). H-scoring was performed using the positive cell detection function in QuPath software v0.3.1. Only cell expansion (15 μm), scoring (Cytoplasm: DAB OD mean or max) and thresholds (0.25, 0.5, 0.75) were changed, while the rest of the parameters were left as default.

### Caerulein-mediated injury

Mice were starved overnight then given 0.1 mg/kg buprenorphine (Vetergesic® 0.3 mg/ml solution) subcutaneously 30 min prior to the first injection. Mice then received six intraperitoneal injections of 80 µg/kg Caerulein Ammonium Salt (Bachem 4030451) in PBS or the equivalent volume of PBS, at hourly intervals.

### Acinar culture

Intact pancreatic acini were extracted based on a hybrid protocol adapted from published protocols (Gout et al., 2013; Geron et al., 2014). Briefly, pancreata from 10-week old mice were dissected and washed with cold Hanks’ Buffered Saline Solution (HBSS, Lonza, BE10-527F), and diced into 1-3 mm^3^ fragments in cold HBSS before centrifugation at 450 x g for 5 min at 4 °C. Supernatant was discarded to remove cell debris and pancreatic fragments were resuspended in 10 ml collagenase IA solution: HBSS, 10 mM HEPES pH 7.2, 200 U/ml collagenase IA (Sigma-Aldrich, 2674), 0.25 mg/ml of trypsin inhibitor (Sigma-Aldrich, T6522). The suspension was incubated at 37 °C for up to 20 min, with mechanical dissociation via periodic pipetting. When digestion to acini was evident, as monitored by phase contrast microscopy, digestion was stopped by adding 10 ml of cold wash (HBSS, 5% FBS, 10 mM HEPES pH 7.2). After initial pelleting at 150 x g for 4 min at 4 °C, samples were washed and recentrifuged three times. The resultant pellet was resuspended in 7 ml of warm Waymouth’s medium (Gibco, 31220023) - supplemented with 1 % Penicillin/Streptomycin, 0.25 mg/ml trypsin inhibitor (Sigma-Aldrich, T652) and 25 ng/ml of EGF (Invitrogen, PHG0311L) - then 100 μm filtered. Acini were settled by gravity from the filtered suspension and then washed 3 x Waymouth’s medium. Acini were then seeded in 6-well plates and cultured in Waymouth’s medium at 37 °C in 5 % CO_2_ overnight, before pelleting and extraction of an SDS-soluble fraction via resuspension and brief sonication in SDS lysis buffer (4% SDS, 150mM NaCl, 50 mM Tris-HCl pH 7.5). Samples were centrifuged at 17 000 x g for 15 min and supernatant was collected. The insoluble pellet was washed with PBS and frozen prior to mass spectrometric analysis.

### Mass spectrometry

To precipitate detergent-soluble samples and facilitate SDS removal, trichloroacetic acid (TCA) was added to 20 % w/v and samples incubated at −20 °C for 1 h then centrifuged at 17 000 x g for 10 min. The pellet was resuspended in 1 mL acetone. Samples were then sonicated to loosen and wash the pellet, then incubated at −20 °C for 10 mins, then centrifuged at 17 000 x g for 10 min. This acetone wash was repeated thrice and then pellets were air dried. Alternatively, detergent-insoluble samples (pellets) were washed in PBS and sonicated to remove residual SDS, prior to centrifugation at 17 000 x g for 10 min. This wash was repeated thrice. Washed pellets of both detergent-soluble and -insoluble fractions were then resolubilised in 200 mM Tris pH 8.5, 6 M guanidium chloride, 20 mM 2-chloroacetamide, 10 mM tris(2-carboxyethyl) phosphine. Samples were vortexed and spun down before incubation at 95 °C for 5 min. Cooled samples were centrifuged briefly before dilution 1:1 in mass spectrometry (MS) grade water (Supelco, 1.15333) containing 0.5 μg/ml of lysyl endopeptidase (Lys-C, Waiko, 121-05063). Samples incubated at 37 °C in a thermomixer at 900 rpm overnight. Samples were then diluted 1:2 in MS grade H2O and 0.5 μg of MS-grade trypsin (Thermo scientific, 90058) added. Samples were incubated at 37 °C in a thermomixer at 900 rpm overnight. Stage tips were prepared (1mm diameter circles of C18 membrane inserted into P200 pipette tips). Membranes were activated with 15 μL MS grade methanol and liquid removed by centrifugation for 2 min at 200 x g. Membranes were equilibrated with 50 μL 0.1% trifluoroacetic acid (TFA) and centrifuged for 3 min at 300 x g. 20 μL of 10 % TFA was added to samples, which were cleared by centrifugation for 5 min at 17 000 x g, then 300 μL was loaded onto membranes. Membranes were centrifuged at 500 x g for 5 min, washed with 50 μL 0.1% TFA and centrifuged at 500 x g. After repeating the washing step, membranes were eluted into 96-well plates with 40 μL of 50% acetonitrile (ACN), 0.05% TFA, centrifuging at 300 g for 3 min. Plates were then vacuum centrifuged for 30 minutes at 45 °C to remove ACN. 15 μL of 0.1% TFA was added, peptide concentration was measured using a Nanodrop 2000 Spectrophotometer, and samples were diluted in 0.1% TFA to a final concentration of 0.2 μg/ml. LC-MS/MS. 1 µg of de-salted peptides were loaded onto a 25 cm emitter (Odyssey, IonOptiks, Australia) using a RSLC-nano uHPLC systems connected to a Fusion Lumos mass spectrometer (both Thermo, UK). Peptides were separated by a 70 min linear gradient from 5% to 30% acetonitrile, 0.05% acetic acid. The mass spectrometer was operated in DIA mode, acquiring a MS 350-1650 Da at 120k resolution followed by MS/MS on 45 windows with 0.5 Da overlap (200-2000 Da) at 30k with a NCE setting of 27. The raw data files underwent analysis and quantification utilizing the DIA-NN software using the default settings. A mouse proteome FASTA file was employed to compare the calculated peptides. For the analysis, the precursor m/z range was defined from 350 to 1650, while the fragment ion m/z range was set from 200 to 2000. Double-pass mode (High Precision) was enabled. Upon completion of the analysis, TSV (tab-separated values) files were generated from the processed data for subsequent analysis and interpretation. Data clean up, statistical analyses and visualisation were performed using Excel and GraphPad Prism.

### Immunoprecipitation

PDAC-V5, PDAC CCPG1-V5 and HEK293FT GFP-REG3A cells were transfected in 10 cm dishes in antibiotic-free culture medium using 6 μg plasmid DNA complexed with Lipofectamine 2000 (ThermoFisher, 11668019) as per manufacturer’s instructions (PDAC cells with pEGFP-REG3 constructs, HEK293FT cells with pdcDNA myc_(6)_-CCPG1 constructs or EV control). The following day, co-immunoprecipitation was performed using V5-Trap Magnetic Agarose beads (PDAC; Proteintech, v5ta) or myc-Trap Magnetic Agarose beads (HEK293FT; Proteintech, ytma). Briefly, cells were washed with ice-cold PBS and lysed in 700 μl NP40 lysis buffer (50mM Tris-HCl, pH 7.5, 150 mM NaCl, 0.5% IGEPAL, 2 mM activated sodium orthovanadate, 20mM sodium fluoride, 10 mM sodium pyrophosphate, 1mM PMSF and Complete Protease Inhibitor (Roche 11697498001). Cells were incubated for 10 min with shaking and lysates were transferred into 1.5 ml microcentrifuge tubes and resuspended 5 times by pipetting. Lysates were incubated 10 min before being cleared by centrifugation at 13 000 x g for 10 min at 4 °C, and input samples removed. Magnetic beads were washed three times in NP40 lysis buffer then rotated with supernatants for 2 h at 4°C. Beads were washed with NP40 lysis buffer three times, then washed once with 50mM Tris-HCl, pH 7.5, 150 mM NaCl. Beads and input samples were boiled in Laemmli buffer prior to immunoblotting.

### Immunofluorescence

Slides were prepared as for immunohistochemistry as described herein, through to antigen retrieval in sodium citrate. Tissue was then permeabilised in 0.1% Triton X-100 in PBS for 10 min and blocked with protein block (5% goat serum, 0.3% Triton X-100 in PBS) for 30 min at RT. After three washes with 0.1% Triton-X-100 in PBS for 2 minutes, tissues were incubated with endogenous mouse IgG blocking (0.13 mg/ml AffiniPure Fab Fragment (Jackson ImmunoResearch, 115-007-003) in PBS) for 1 hour at RT. Primary antibodies were diluted in protein block (Abcam, ab64226) and incubated overnight at 4 °C. After 3 x 5 min PBS washes, slides were then incubated with secondary antibody (1:200 in PBS) for 1 h at RT (AlexaFlour488 anti-rabbit and AlexaFluor594 anti-mouse, for staining mouse tissue; AlexaFluor594 anti-rabbit and AlexaFluor647 anti-mouse, for staining human tissue; Invitrogen, A11034, A11005, A11037, A-21235, respectively). Pan-cytokeratin staining on human samples involved a subsequent incubation in pan-Cytokeratin-AlexaFluor532 (1:200 in PBS; NBP2-33200AF532, Novus Biologicals) for 1.5 h at RT. Slides were then washed 2 x PBS for 5 min, incubated with 100 μl of DAPI (1 μg/ml in PBS) for 10 min and washed again. Samples were mounted with Dako mounting media and a glass coverslip, sealed with nail varnish.

### Confocal microscopy

Generally, images were captured with an Olympus FV3000 Confocal Laser Scanning Microscope using Fluoview FV31S-SW (version 2.4.1.198) software and an Olympus UPlanSApo 60× 1.35NA Oil objective. Images were further analysed in ImageJ 2.9.0. In the case of human sample immunofluorescence, images were captured on a Stellaris 8 confocal microscope using LAS X software (Leica Microsystems UK). The microscope consists of a DMi8 inverted microscope and is equipped with a white light laser, enabling free tuning of the excitation wavelength between 440-790nm, and a 405nm diode laser for UV. Light detection is via two HyD S (Silicone Multi-Pixel Photon Counter) & two HyD X (GaAsP Hybrid) detectors. Images were captured using 60x and 100x objective lenses. Images were further analysed in ImageJ 2.9.0 (NIH, Bethesda, USA).

### Super-resolution microscopy

Tissue sections were processed for immunofluorescence using the protocol outline hereinand images captured using Super Resolution Optical Photon Reassignment Microscopy on a Nikon Eclipse Ti2 inverted microscope stand, using a Yokogawa CSU-W1 spinning disk unit with 50 um and SoRa disks (Nikon Instruments). 405, 488 and 561nm laser lines were directed in alternating pairs through a 100x 1.35NA Silicone immersion lens (Nikon Instruments) onto the sample and back scattered light was separated using a Di01-T405/488/568/647 dichroic mirror (Semrock) and 447/60, 525/60 and 641/75 emission filters. Images were captured using CMOS dual Teledyne Photometrics Prime 95B cameras (Teledyne Photometrics). Image z-step size was set to 0.1 μm and images were acquired, denoised and deconvolved using the Nikon NIS-Elements Advanced Research (Ver 5.30.05) software. Images were further analysed in Imaris 10.0.1 (Oxford Instruments).

### Immunogold transmission electron microscopy

Mice were fixed by intracardiac perfusion with 10 ml of fresh EM-grade 0.1% glutaraldehyde/ 2% paraformaldehyde in 0.1 M Sorensen’s phosphate buffer (PB) at 2 ml / min. Pancreas was dissected and diced into 1 mm^3^ fragments in fresh fixative, then incubated for 4 h at RT. Fixed tissue was washed 3 x in Sorensen’s PB. Samples were dehydrated in increasing concentrations of ethanol (50%, 70%, 90%, 100%, 100%, 100%) for 15 min followed by 2 x 10 min incubations in propylene oxide. Samples were incubated in LR white resin overnight, changed into the same resin twice more and baked at 60 °C in an oven overnight. 60 nm thick sections were cut from the cured resin and placed on a gold grid for immunolabelling. Grids were incubated in PBS-Tween 20 for 10 min (0.1% Tween 20 in PBS), then blocked in (2.5% BSA and 2.5% Goat serum in PBS with 0.05% Tween 20) for 30 min, then stained in primary antibody in Ig diluent (1% BSA in PBS) (1:10 REG3A/B, 1:50 ZG16) for 2 h, before washing in PBS-Tween 20 and incubation in secondary antibodies (α-Mouse IgG Gold (Sigma-Aldrich, G7652) 1:20 in Ig diluent). Grids were then washed in PBS and then ddH_2_O, air dried and counterstained with uranyl acetate and lead citrate and viewed on a JEOL JEM-1400 Plus electron microscope (JEOL), collecting images on a GATAN OneView camera (Gatan).

### Human tissue

Normal pancreatic tissue was from an anonymous male donor died aged 30 with no adverse medical history, for whom an acceptable recipient could not be found. Use in this study was with full ethical approval (Scotland A Research Ethics Committee 08/MRE00/14). Tumor-adjacent normal pancreatic tissue was from a commercially available tissue microarray (AMSBio, HPanA060CS02).

### mRNA-Seq

RNA integrity number (RIN) was measured using a Tapestation 4200 (Agilent). Selected samples (RIN > 5.7) were subject to poly(A)-enrichment, mRNA library preparation and sequencing by Novogene (Cambridge, UK). Data obtained was processed using the nf-core/rnaseq pipeline v3.12 (Ewels et al., 2020) and the pipeline was executed with Nextflow v23.04.2 (Di Tommaso et al., 2017). FASTQ files were merged, trimmed to remove adapters and low-quality regions with Trim Galore 3.4, aligned using STAR 2.6.1d (Dobin et al., 2013) and quantified using Salmon 1.10.1 (Patro et al., 2017). QC report was obtained using MultiQC v1.14 (Ewels et al., 2016). Subsequent analyses were performed in R v4.2.2. Counts were normalised using variance-stabilizing transformation (VST) and principal component analysis (PCA) was performed. Differentially expressed genes were identified with Deseq2 v1.38.3 (Love et al., 2014) using a generalised linear model with negative binomial distribution, with genotype as the main factor and the Wald test. Log2 fold values were shrunk using the apeglm v1.20.0 method (Zhu et al., 2019). Genes were annotated using the R packages biomaRt v2.54.1 and org.Mm.eg.db v.3.16.0. Heatmaps were created using the R packages pheatmap 1.0.12 and values presente in GraphPad. Gene Set Enrichment Analysis (GSEA) was performed in GSEA v4.3.2 (Subramanian et al., 2005) using all mouse annotated gene sets from MSigDB (Castanza et al., 2023) except for the “M1-positional gene sets” (Supplemental Table 3) and data was imported into R. Data was plotted for visualisation using the R package ggplot2.

### Spatial transcriptomics

Slides were prepared following the NanoString GeoMx DSP Manual RNA Assay Slide Preparation protocol (NanoString technologies, WA, USA). Briefly, 2 x 5-μm-thick FFPE tissue sections were placed in the scan area of each slide. Slides were dried at 60 °C for 60 min prior to dewaxing. Slides were immersed three times in xylene for 5 min, twice in 100% EtOH for 5 min, once in 95% EtOH for 5 min and 1 minute in PBS. Antigen retrieval was performed in IHC Antigen Retrieval Solution (Invitrogen, 00-4956-58) for 15 min at 100 °C. Slides were then incubated for 5 min in PBS at RT and incubated with 1 μg/ml Proteinase K solution (Invitrogen, AM2546) in PBS. Slides were then incubated in 10% Neutral Buffered Formalin (NBF, Cell Path, BAF-0100-01A) for 5 min, twice in Stop Buffer (0.1 M Tris, 0.1 M glycine in DEPC-treated water) for 5 min and 1 x PBS. Slides were moved to a hybridisation chamber and incubated with buffer R containing the Whole Transcriptome Atlas Probe Mix (Nanostring, 999064) overnight at 37 °C. Coverslips were removed by immersion in 2X SSC (Sigma-Aldrich, S6639) and slides washed 2 x stringent wash solution (2X SSC, 50% formamide) for 25 min at 37 °C, and twice in 2x SSC for 2 min at RT. Slides were transferred to a humidity chamber and incubated with Serum-free protein block (Dako, X0909) for 30 min at RT. Slides were incubated with primary antibodies diluted in protein block (REG3A/B 1:50, Amylase 1:300) for 1 h at RT. After 2 x 5 min wash in 2x SSC, slides were incubated with secondary antibodies diluted 1:400 in protein block for 30 min. Slides were washed 2 x 5 min with 2x SCC and incubated for 30 min at RT with Goat α-Mouse AlexaFluor 594 (Invitrogen, A-11032) and Goat α-Rabbit AlexaFluor 647 (Invitrogen, A-21245) (1:400 in protein block). Slides were then washed 2 x 5 min in 2 x SCC before incubation with PanCK AF532 (1:100 Novus Biologicals, NBP2-33200AF532), SYTO 13 (1:20 000, Invitrogen, S7575) in protein block for 1 h at RT. Slides were washed twice in 2X SSC for 5 min, then scanned with GeoMx Digital Spatial Profiler (GeoMX DSP) (Nanostring). Once scanned, slides were visualised in the GeoMx DPS Scan Workspace and regions of interest (ROIs) were selected.. To separate the different cell populations within the ROIs, segments were defined by choosing which antibodies describe each population. For each ROI, intensity thresholds were manually adjusted and after all segments were created, non-specific segments were deleted. A total of 95 segmented samples from multiple ROIs were collected in the 96-well DPS collection plate and sent to the Genetics Core for library preparation and sequencing on an Illumina NextSeq 2000 platform (Illumina Inc, 20038898). FASTQ files were transformed into digital count conversion (DCC) files using the Nanostring GeoMx NGS pipeline and uploaded to the GeoMX DSP for analysis. Samples were analysed in the GeoMX DSP Data Analysis Suite following the manufacturer’s protocols. Briefly, after counts were uploaded, biological probe QC removed poorly performing probes (≤10%). Targets were filtered and only those with expression > limit of quantitation (LOQ) in at least 5% of the segments were retained. Data was normalised using Q3 normalisation and PCA was performed (GeoScript Hub). Differentially-expressed genes were selected from pair-wise t-test comparison of acinar populations (not ADM), with a criterion of inclusion of log fold-change magnitude of > 1 and p-value < 0.05. Normalised data were exported and heatmaps were created using the R package pheatmap with scaled values across each row, then values from heatmaps presented in Graphpad. Gene Set Enrichment Analysis (GSEA) was performed in GSEA v4.3.2 (Broad Institute) and data was imported into R. Gene set variation analysis (GSVA) was performed using the R package GSVA v1.46.0 (Hanzelmann et al., 2013). Data were plotted for visualisation using the R package ggplot2.

### REG3B expression assays

Delivery of rAAV expressing *Reg3b* isoforms or *luciferase-V5* negative control to mouse pancreas was performed via a single intraperitoneal (i.p.) injection of neat virus preparation (20 ml/kg, 2 x 10^9^ viral genome / μL). Mice were monitored to ensure full recovery prior to return to general husbandry. Mice were terminated at 5 weeks post-injection by a rising concentration of carbon dioxide. Plasma was obtained by extraction of blood via cardiac puncture using a heparinised syringe followed by centrifugation at 1500 g for 10 min. Equal volumes of plasma were boiled in Laemmli buffer prior to immunoblot analysis. Pancreas was dissected and subjected to standard SDS protein fractionation, RNA extraction, and formalin fixation & paraffin embedding as described elsewhere herein. In the case of HEK293FT cells, transfection with empty and *Reg3b* mammalian expression vectors was using Lipofectamine 2000, according to manufacturer’s instructions. Conditioned medium was collected and boiled in Laemmli buffer prior to immunoblot and PBS-washed cells were subject to detergent soluble/insoluble fractionation as described herein for pancreatic tissues.

### Plasmids

Plasmids are tabulated below. Detailed maps available from authors upon request.

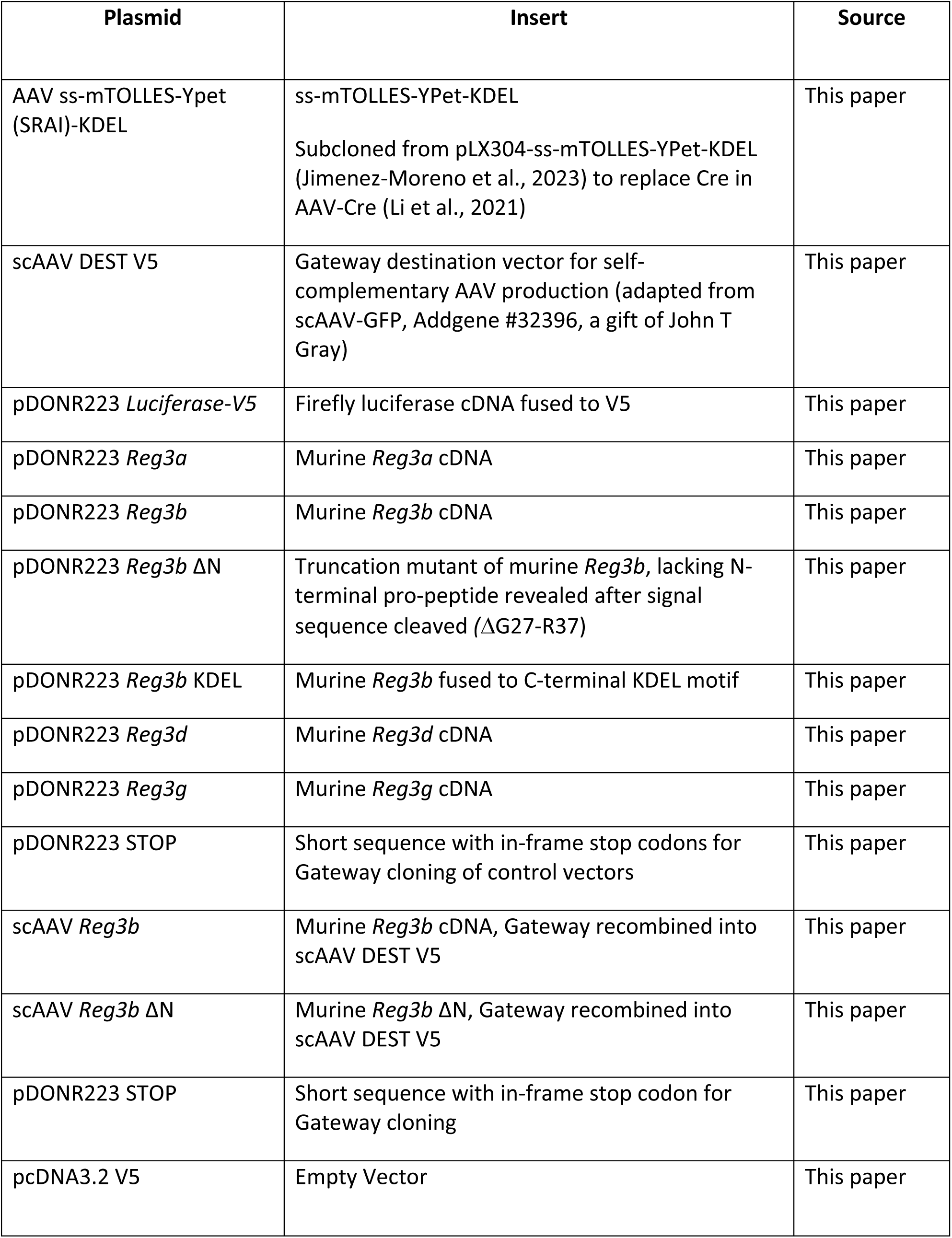
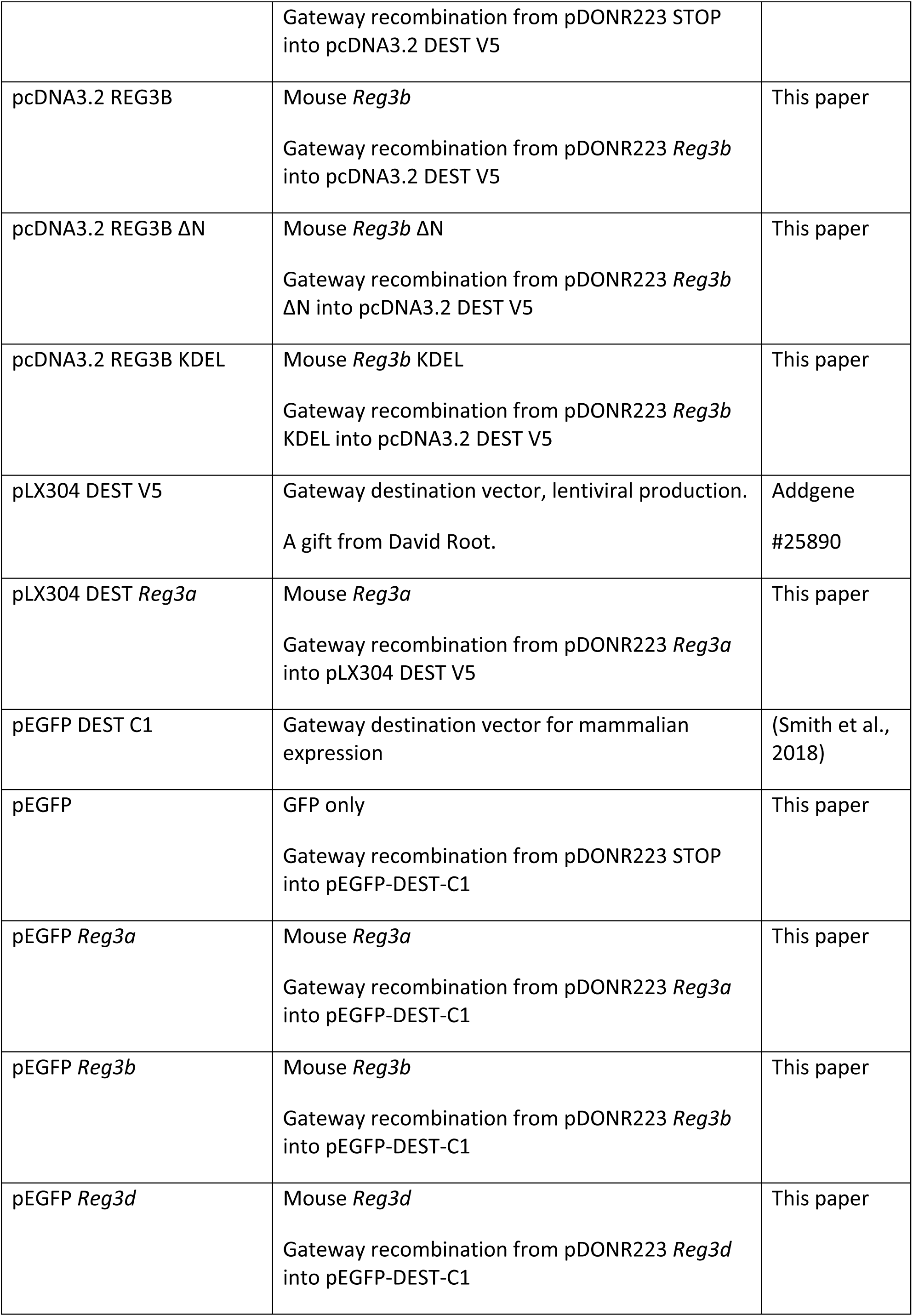
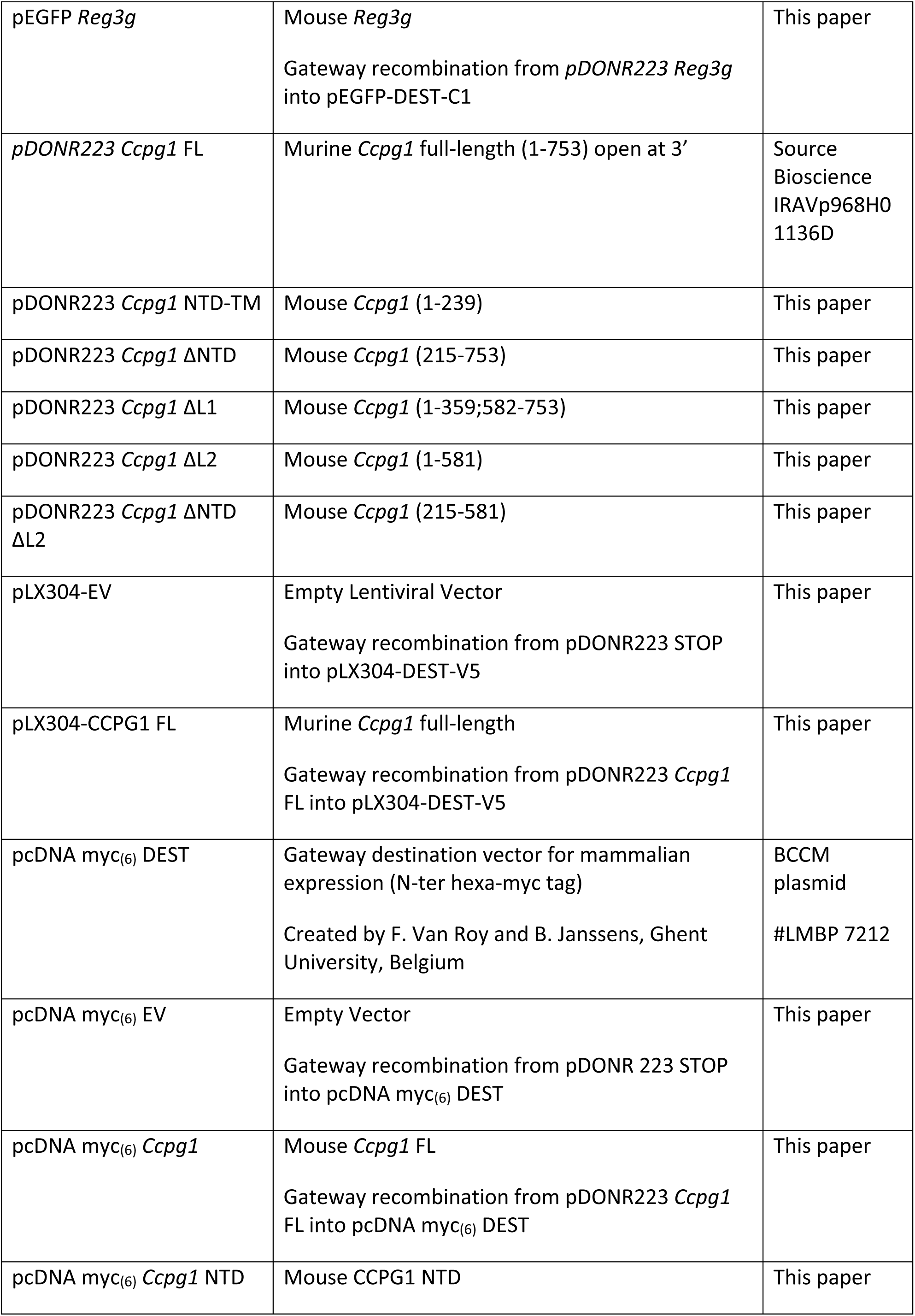
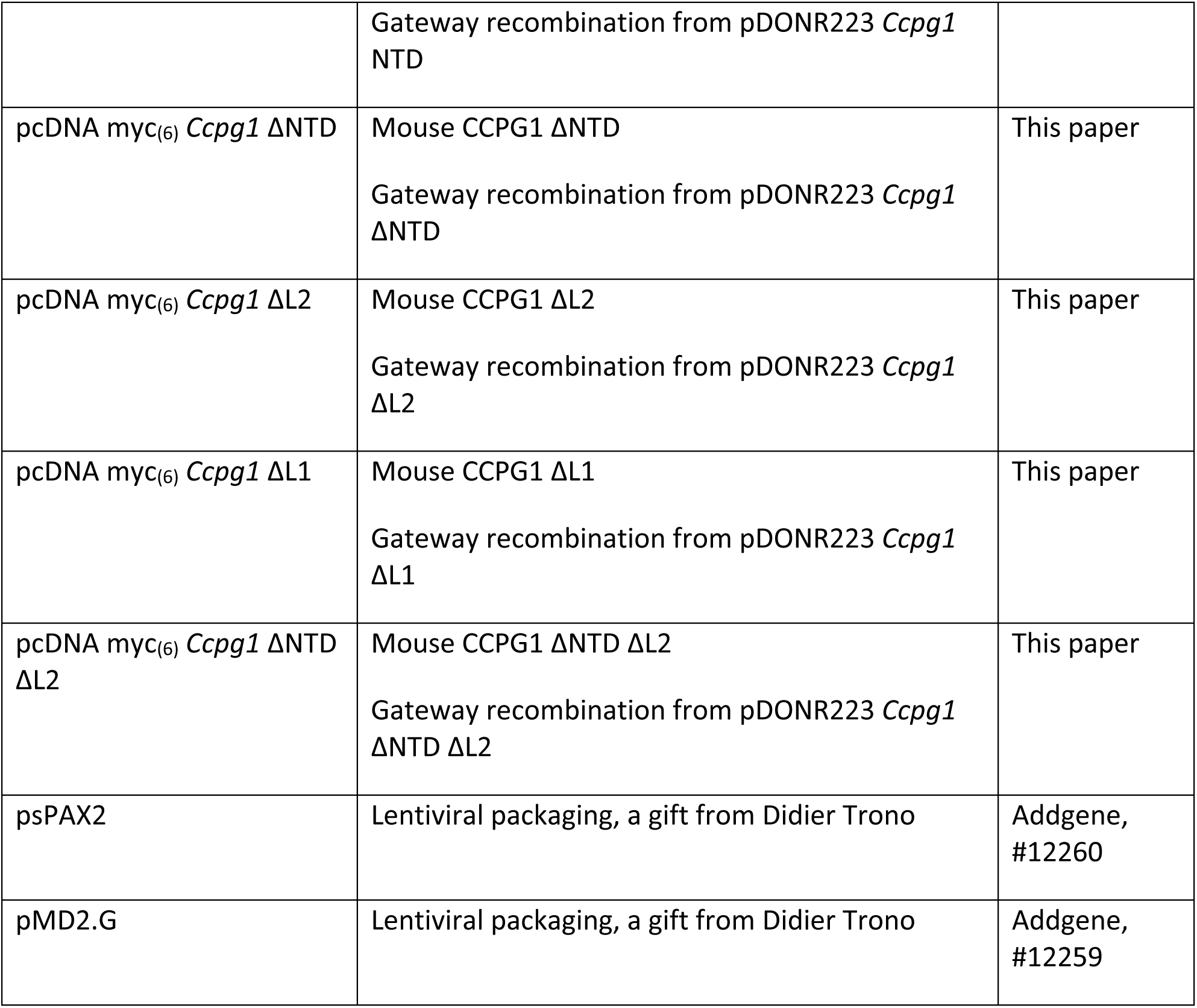

### Oligonucleotides

qRT-PCR primers are tabulated below

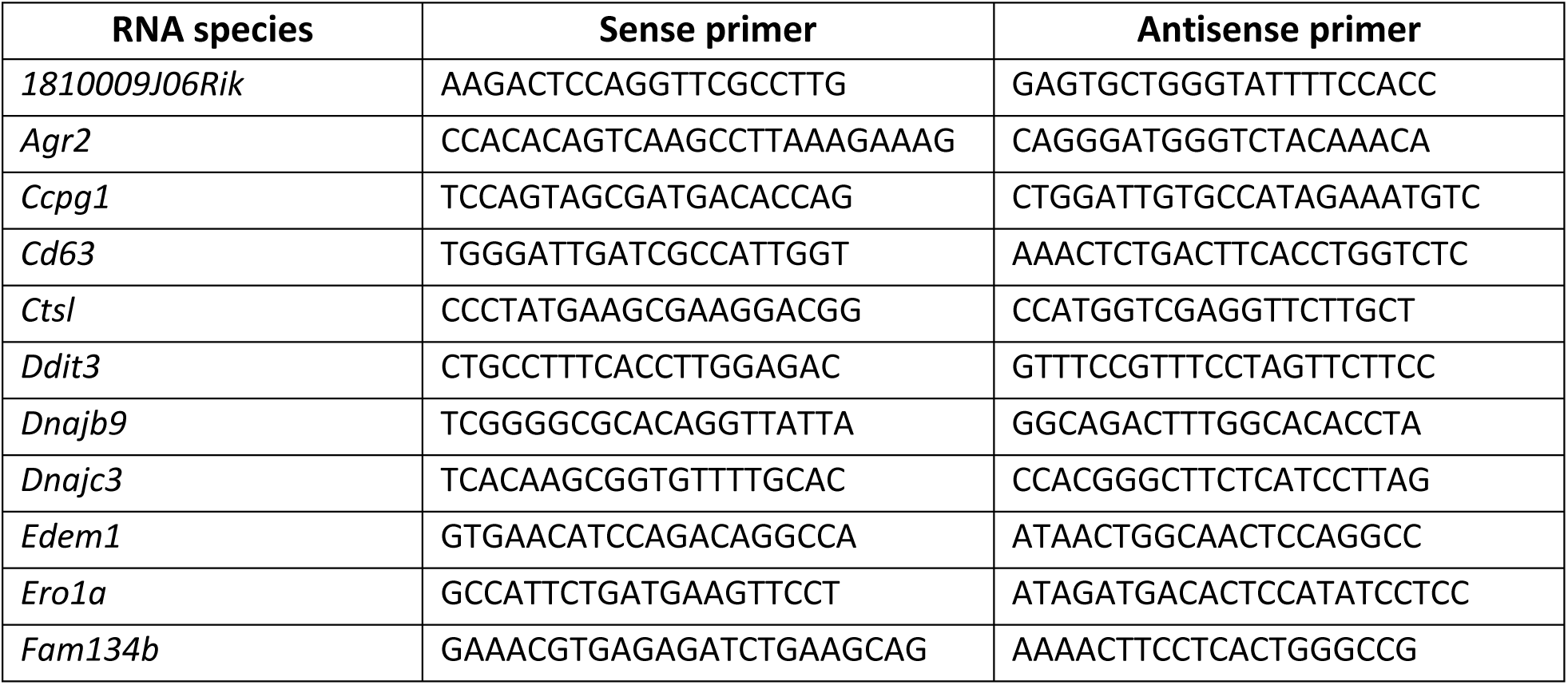
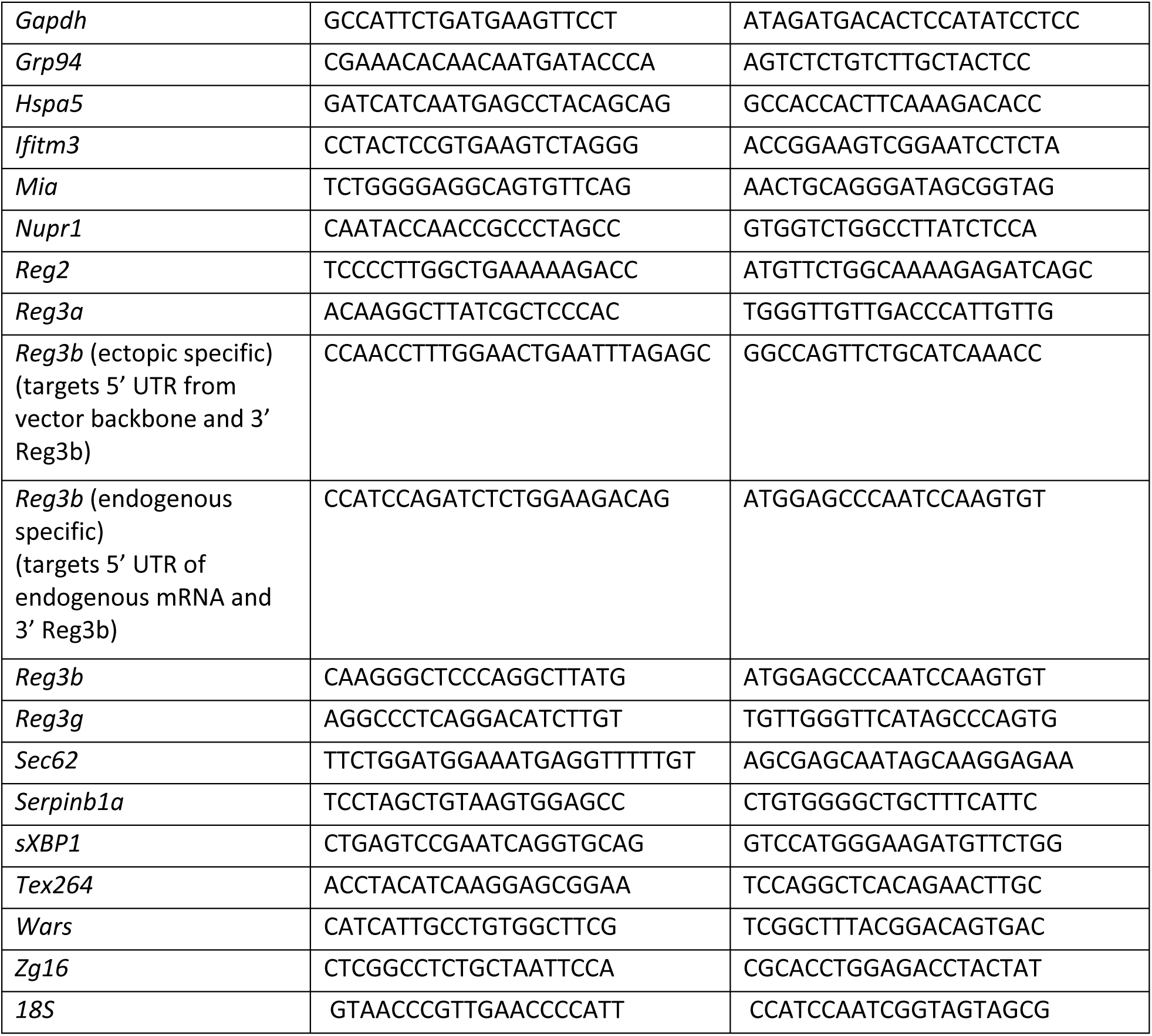

### Antibodies

Antibodies used in this study are tabulated below along with use: IB = immunoblot, IHC = immunohistochemistry, IF = immunofluorescence, SP = immunofluorescence for spatial transcriptomics, IG = immunogold TEM

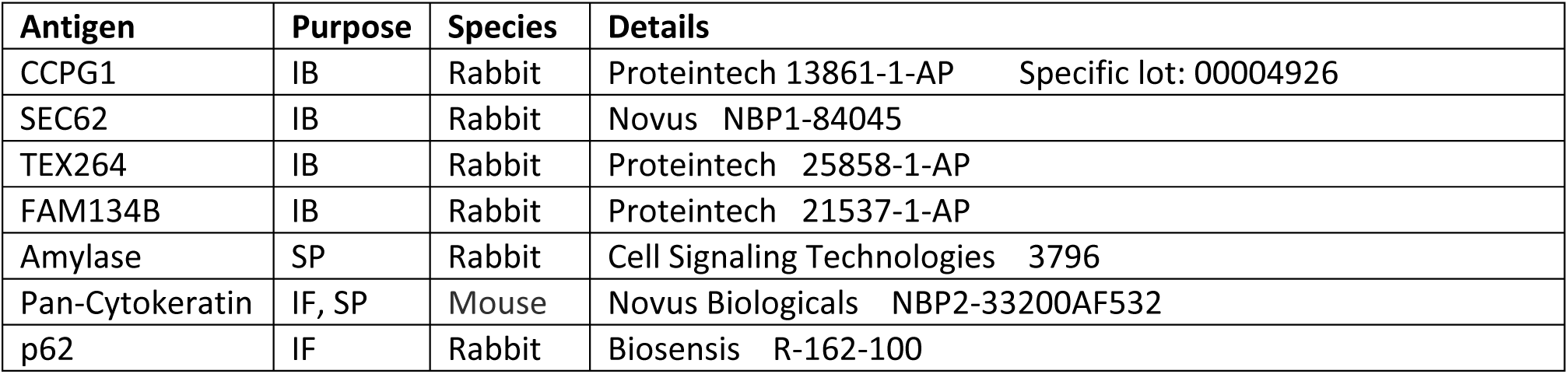
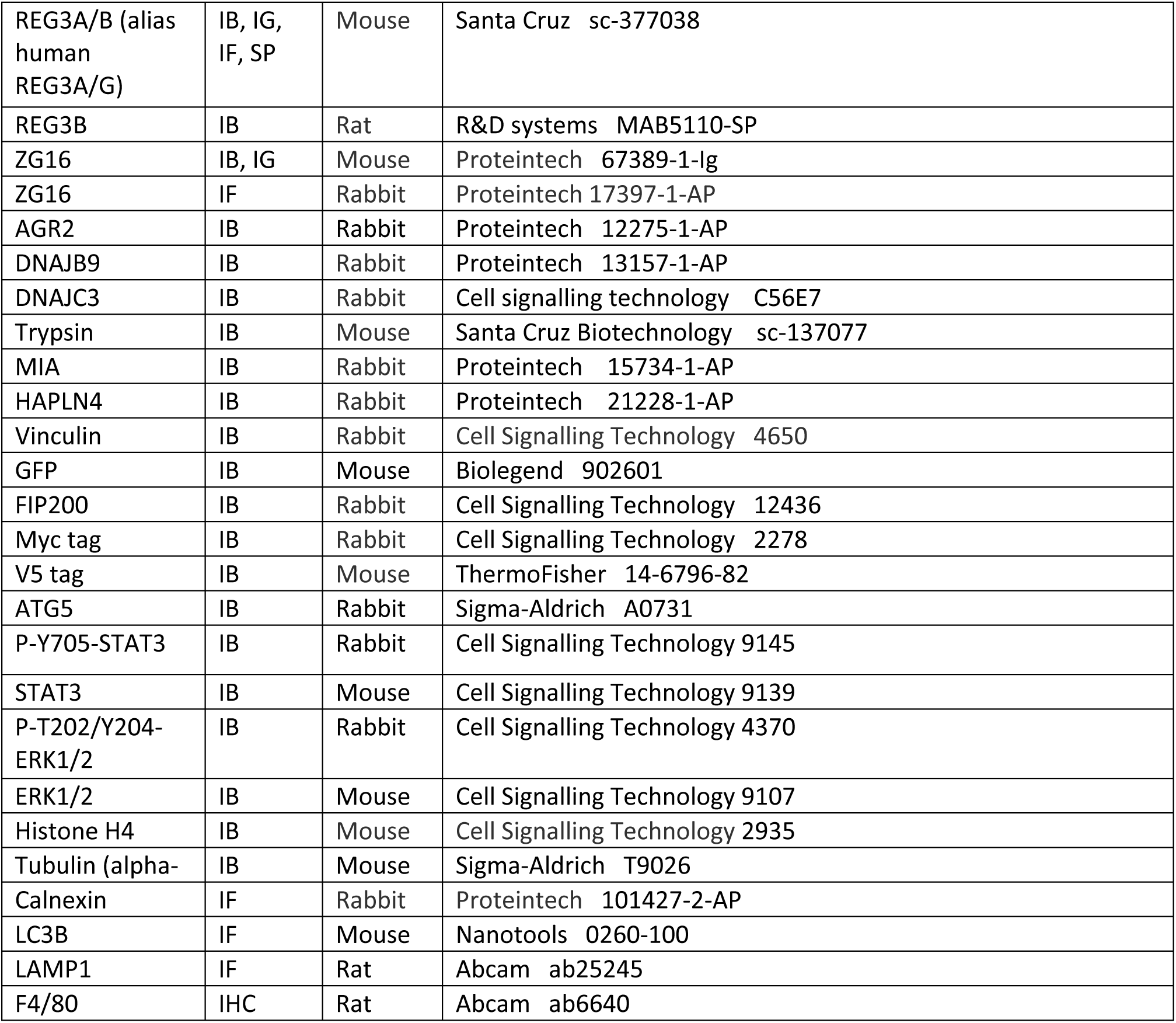

### Code availability

Standard R workflows for bioinformatic analyses were used as described elsewhere in these Methods and are available from the authors upon request.

### Statistics

Unless stated otherwise, all statistical tests were performed as described in figure legends using GraphPad v10. Where applicable, all tests were two-tailed. Grubb’s outlier tests were performed prior to testing to remove up to one outlier. Where individual values within an independent replicate (or pair of sibling mice) is normalised to control, appropriate tests that deal with this such as the 1-sample t-test were employed. Multiple testing correction on p-values was performed as described in individual figure legends.

### Data set deposition

The mass spectrometry proteomics data have been deposited to the ProteomeXchange Consortium via the PRIDE partner repository (Perez-Riverol et al., 2022) with the dataset identifier PXD055161. Reviewer login details: Username: and password: KU2MEpU5GUMy. mRNA-Seq data are available at NCBI GEO (https://www.ncbi.nlm.nih.gov/geo/query/acc.cgi?acc=GSE274978, GSE274978) (password for review: gxcfgqguprghfez). Spatial transcriptomic data are available at NCBI GEO (https://www.ncbi.nlm.nih.gov/geo/query/acc.cgi?acc=GSE274979, GSE274979) (password for review: ypkvioiqzhsvzqd).

## Supplemental information legends

Tables S1-S4. Excel files containing data too large to fit in a PDF.

**Supplementary Table 1**

Mass spectrometry data from acini isolated from *KC* and *KC Ccpg1^ΔPANC^*mice (detergent-soluble and -insoluble extracts). Reporting imputed LFQ values, fold-change and p-values.

**Supplementary Table 2**

Processed mRNA-Seq data from bulk pancreas of *KC Ccpg1^+/+^*and *KC Ccpg1^ΔPANC^*mice. Reporting normalised reads and derived statistics (means, standard deviation and mean fold change).

**Supplementary Table 3**

Bespoke gene lists employed in GSEA and GSVA analyses to probe for pancreatic injury and for UPR, alongside MSigDB collections used alongside in the GSEA analyses.

**Supplementary Table 4**

Spatial transcriptomic data across *KC* mouse pancreata (four mice).

## Notes

https://www.ncbi.nlm.nih.gov/geo/query/acc.cgi?acc=GSE274978

https://www.ncbi.nlm.nih.gov/geo/query/acc.cgi?acc=GSE274979

### Competing Interest Statement

The authors have declared no competing interest.

## References

Alonso-Curbelo, D., Ho, Y.J., Burdziak, C., Maag, J.L.V., Morris, J.P.t., Chandwani, R., Chen, H.A., Tsanov, K.M., Barriga, F.M., Luan, W., et al. (2021). A gene-environment-induced epigenetic program initiates tumorigenesis. Nature 590, 642–648.

Aman, Y., Schmauck-Medina, T., Hansen, M., Morimoto, R.I., Simon, A.K., Bjedov, I., Palikaras, K., Simonsen, A., Johansen, T., Tavernarakis, N., et al. (2021). Autophagy in healthy aging and disease. Nat Aging 1, 634–650.

An, H., Ordureau, A., Paulo, J.A., Shoemaker, C.J., Denic, V., and Harper, J.W. (2019). TEX264 Is an Endoplasmic Reticulum-Resident ATG8-Interacting Protein Critical for ER Remodeling during Nutrient Stress. Mol Cell 74, 891–908 e810.

Ardito, C.M., Gruner, B.M., Takeuchi, K.K., Lubeseder-Martellato, C., Teichmann, N., Mazur, P.K., Delgiorno, K.E., Carpenter, E.S., Halbrook, C.J., Hall, J.C., et al. (2012). EGF receptor is required for KRAS-induced pancreatic tumorigenesis. Cancer Cell 22, 304–317.

Aurnhammer, C., Haase, M., Muether, N., Hausl, M., Rauschhuber, C., Huber, I., Nitschko, H., Busch, U., Sing, A., Ehrhardt, A., and Baiker, A. (2012). Universal real-time PCR for the detection and quantification of adeno-associated virus serotype 2-derived inverted terminal repeat sequences. Hum Gene Ther Methods 23, 18–28.

Bailey, J.M., Hendley, A.M., Lafaro, K.J., Pruski, M.A., Jones, N.C., Alsina, J., Younes, M., Maitra, A., McAllister, F., Iacobuzio-Donahue, C.A., and Leach, S.D. (2016). p53 mutations cooperate with oncogenic Kras to promote adenocarcinoma from pancreatic ductal cells. Oncogene 35, 4282–4288.

Braxton, A.M., Kiemen, A.L., Grahn, M.P., Forjaz, A., Parksong, J., Mahesh Babu, J., Lai, J., Zheng, L., Niknafs, N., Jiang, L., et al. (2024). 3D genomic mapping reveals multifocality of human pancreatic precancers. Nature 629, 679–687.

Burdziak, C., Alonso-Curbelo, D., Walle, T., Reyes, J., Barriga, F.M., Haviv, D., Xie, Y., Zhao, Z., Zhao, C.J., Chen, H.A., et al. (2023). Epigenetic plasticity cooperates with cell-cell interactions to direct pancreatic tumorigenesis. Science 380, eadd5327.

Castanza, A.S., Recla, J.M., Eby, D., Thorvaldsdottir, H., Bult, C.J., and Mesirov, J.P. (2023). Extending support for mouse data in the Molecular Signatures Database (MSigDB). Nat Methods 20, 1619–1620.

Chen, M., Maeng, K., Nawab, A., Francois, R.A., Bray, J.K., Reinhard, M.K., Boye, S.L., Hauswirth, W.W., Kaye, F.J., Aslanidi, G., et al. (2017). Efficient Gene Delivery and Expression in Pancreas and Pancreatic Tumors by Capsid-Optimized AAV8 Vectors. Hum Gene Ther Methods 28, 49–59.

Chen, Z., Downing, S., and Tzanakakis, E.S. (2019). Four Decades After the Discovery of Regenerating Islet-Derived (Reg) Proteins: Current Understanding and Challenges. Front Cell Dev Biol 7, 235.

Chino, H., Hatta, T., Natsume, T., and Mizushima, N. (2019). Intrinsically Disordered Protein TEX264 Mediates ER-phagy. Mol Cell 74, 909–921 e906.

Chondronasiou, D., Martinez de Villarreal, J., Melendez, E., Lynch, C.J., Pozo, N.D., Kovatcheva, M., Aguilera, M., Prats, N., Real, F.X., and Serrano, M. (2022). Deciphering the roadmap of in vivo reprogramming toward pluripotency. Stem Cell Reports 17, 2501–2517.

Cui Zhou, D., Jayasinghe, R.G., Chen, S., Herndon, J.M., Iglesia, M.D., Navale, P., Wendl, M.C., Caravan, W., Sato, K., Storrs, E., et al. (2022). Spatially restricted drivers and transitional cell populations cooperate with the microenvironment in untreated and chemo-resistant pancreatic cancer. Nat Genet 54, 1390–1405.

Del Poggetto, E., Ho, I.L., Balestrieri, C., Yen, E.Y., Zhang, S., Citron, F., Shah, R., Corti, D., Diaferia, G.R., Li, C.Y., et al. (2021). Epithelial memory of inflammation limits tissue damage while promoting pancreatic tumorigenesis. Science 373, eabj0486.

Di Tommaso, P., Chatzou, M., Floden, E.W., Barja, P.P., Palumbo, E., and Notredame, C. (2017). Nextflow enables reproducible computational workflows. Nat Biotechnol 35, 316–319.

Dobin, A., Davis, C.A., Schlesinger, F., Drenkow, J., Zaleski, C., Jha, S., Batut, P., Chaisson, M., and Gingeras, T.R. (2013). STAR: ultrafast universal RNA-seq aligner. Bioinformatics 29, 15–21.

Dumartin, L., Alrawashdeh, W., Trabulo, S.M., Radon, T.P., Steiger, K., Feakins, R.M., di Magliano, M.P., Heeschen, C., Esposito, I., Lemoine, N.R., and Crnogorac-Jurcevic, T. (2017). ER stress protein AGR2 precedes and is involved in the regulation of pancreatic cancer initiation. Oncogene 36, 3094–3103.

Ewels, P., Magnusson, M., Lundin, S., and Kaller, M. (2016). MultiQC: summarize analysis results for multiple tools and samples in a single report. Bioinformatics 32, 3047–3048.

Ewels, P.A., Peltzer, A., Fillinger, S., Patel, H., Alneberg, J., Wilm, A., Garcia, M.U., Di Tommaso, P., and Nahnsen, S. (2020). The nf-core framework for community-curated bioinformatics pipelines. Nat Biotechnol 38, 276–278.

Falvo, D.J., Grimont, A., Zumbo, P., Fall, W.B., Yang, J.L., Osterhoudt, A., Pan, G., Rendeiro, A.F., Meng, Y., Wilkinson, J.E., et al. (2023). A reversible epigenetic memory of inflammatory injury controls lineage plasticity and tumor initiation in the mouse pancreas. Dev Cell 58, 2959–2973 e2957.

Ferreira, R.M.M., Sancho, R., Messal, H.A., Nye, E., Spencer-Dene, B., Stone, R.K., Stamp, G., Rosewell, I., Quaglia, A., and Behrens, A. (2017). Duct- and Acinar-Derived Pancreatic Ductal Adenocarcinomas Show Distinct Tumor Progression and Marker Expression. Cell Rep 21, 966–978.

Fumagalli, F., Noack, J., Bergmann, T.J., Cebollero, E., Pisoni, G.B., Fasana, E., Fregno, I., Galli, C., Loi, M., Solda, T., et al. (2016). Translocon component Sec62 acts in endoplasmic reticulum turnover during stress recovery. Nat Cell Biol 18, 1173–1184.

Geron, E., Schejter, E.D., and Shilo, B.Z. (2014). Assessing the secretory capacity of pancreatic acinar cells. J Vis Exp.

Gidekel Friedlander, S.Y., Chu, G.C., Snyder, E.L., Girnius, N., Dibelius, G., Crowley, D., Vasile, E., DePinho, R.A., and Jacks, T. (2009). Context-dependent transformation of adult pancreatic cells by oncogenic K-Ras. Cancer Cell 16, 379–389.

Gopalan, V., Singh, A., Rashidi Mehrabadi, F., Wang, L., Ruppin, E., Arda, H.E., and Hannenhalli, S. (2021). A Transcriptionally Distinct Subpopulation of Healthy Acinar Cells Exhibit Features of Pancreatic Progenitors and PDAC. Cancer Res 81, 3958–3970.

Gopinathan, A., Morton, J.P., Jodrell, D.I., and Sansom, O.J. (2015). GEMMs as preclinical models for testing pancreatic cancer therapies. Dis Model Mech 8, 1185–1200.

Gout, J., Pommier, R.M., Vincent, D.F., Kaniewski, B., Martel, S., Valcourt, U., and Bartholin, L. (2013). Isolation and culture of mouse primary pancreatic acinar cells. J Vis Exp.

Graf, R., Schiesser, M., Scheele, G.A., Marquardt, K., Frick, T.W., Ammann, R.W., and Bimmler, D. (2001). A family of 16-kDa pancreatic secretory stress proteins form highly organized fibrillar structures upon tryptic activation. J Biol Chem 276, 21028–21038.

Grumati, P., Morozzi, G., Holper, S., Mari, M., Harwardt, M.I., Yan, R., Muller, S., Reggiori, F., Heilemann, M., and Dikic, I. (2017). Full length RTN3 regulates turnover of tubular endoplasmic reticulum via selective autophagy. Elife 6.

Guerra, C., Collado, M., Navas, C., Schuhmacher, A.J., Hernandez-Porras, I., Canamero, M., Rodriguez-Justo, M., Serrano, M., and Barbacid, M. (2011). Pancreatitis-induced inflammation contributes to pancreatic cancer by inhibiting oncogene-induced senescence. Cancer Cell 19, 728–739.

Guerra, C., Schuhmacher, A.J., Canamero, M., Grippo, P.J., Verdaguer, L., Perez-Gallego, L., Dubus, P., Sandgren, E.P., and Barbacid, M. (2007). Chronic pancreatitis is essential for induction of pancreatic ductal adenocarcinoma by K-Ras oncogenes in adult mice. Cancer Cell 11, 291–302.

Guo, J.Y., Chen, H.Y., Mathew, R., Fan, J., Strohecker, A.M., Karsli-Uzunbas, G., Kamphorst, J.J., Chen, G., Lemons, J.M., Karantza, V., et al. (2011). Activated Ras requires autophagy to maintain oxidative metabolism and tumorigenesis. Genes Dev 25, 460–470.

Habbe, N., Shi, G., Meguid, R.A., Fendrich, V., Esni, F., Chen, H., Feldmann, G., Stoffers, D.A., Konieczny, S.F., Leach, S.D., and Maitra, A. (2008). Spontaneous induction of murine pancreatic intraepithelial neoplasia (mPanIN) by acinar cell targeting of oncogenic Kras in adult mice. Proc Natl Acad Sci U S A 105, 18913–18918.

Hanzelmann, S., Castelo, R., and Guinney, J. (2013). GSVA: gene set variation analysis for microarray and RNA-seq data. BMC Bioinformatics 14, 7.

Hetz, C. (2012). The unfolded protein response: controlling cell fate decisions under ER stress and beyond. Nat Rev Mol Cell Biol 13, 89–102.

Hingorani, S.R., Petricoin, E.F., Maitra, A., Rajapakse, V., King, C., Jacobetz, M.A., Ross, S., Conrads, T.P., Veenstra, T.D., Hitt, B.A., et al. (2003). Preinvasive and invasive ductal pancreatic cancer and its early detection in the mouse. Cancer Cell 4, 437–450.

Ishii, S., Chino, H., Ode, K.L., Kurikawa, Y., Ueda, H.R., Matsuura, A., Mizushima, N., and Itakura, E. (2023). CCPG1 recognizes endoplasmic reticulum luminal proteins for selective ER-phagy. Mol Biol Cell 34, ar29.

Jimenez-Moreno, N., Salomo-Coll, C., Murphy, L.C., and Wilkinson, S. (2023). Signal-Retaining Autophagy Indicator as a Quantitative Imaging Method for ER-Phagy. Cells 12.

Katayama, H., Hama, H., Nagasawa, K., Kurokawa, H., Sugiyama, M., Ando, R., Funata, M., Yoshida, N., Homma, M., Nishimura, T., et al. (2020). Visualizing and Modulating Mitophagy for Therapeutic Studies of Neurodegeneration. Cell 181, 1176–1187 e1116.

Khaminets, A., Heinrich, T., Mari, M., Grumati, P., Huebner, A.K., Akutsu, M., Liebmann, L., Stolz, A., Nietzsche, S., Koch, N., et al. (2015). Regulation of endoplasmic reticulum turnover by selective autophagy. Nature 522, 354–358.

Kirkegard, J., Mortensen, F.V., and Cronin-Fenton, D. (2017). Chronic Pancreatitis and Pancreatic Cancer Risk: A Systematic Review and Meta-analysis. Am J Gastroenterol 112, 1366–1372.

Klaips, C.L., Jayaraj, G.G., and Hartl, F.U. (2018). Pathways of cellular proteostasis in aging and disease. J Cell Biol 217, 51–63.

Klomp, J.A., Klomp, J.E., Stalnecker, C.A., Bryant, K.L., Edwards, A.C., Drizyte-Miller, K., Hibshman, P.S., Diehl, J.N., Lee, Y.S., Morales, A.J., et al. (2024). Defining the KRAS- and ERK-dependent transcriptome in KRAS-mutant cancers. Science 384, eadk0775.

Kopp, J.L., von Figura, G., Mayes, E., Liu, F.F., Dubois, C.L., Morris, J.P.t., Pan, F.C., Akiyama, H., Wright, C.V., Jensen, K., et al. (2012). Identification of Sox9-dependent acinar-to-ductal reprogramming as the principal mechanism for initiation of pancreatic ductal adenocarcinoma. Cancer Cell 22, 737–750.

Lee, A.Y.L., Dubois, C.L., Sarai, K., Zarei, S., Schaeffer, D.F., Sander, M., and Kopp, J.L. (2019). Cell of origin affects tumour development and phenotype in pancreatic ductal adenocarcinoma. Gut 68, 487–498.

Lee, E., Jung, Y.J., Park, Y.R., Lim, S., Choi, Y.J., Lee, S.Y., Kim, C.H., Mun, J.Y., and Chung, W.S. (2022). A distinct astrocyte subtype in the aging mouse brain characterized by impaired protein homeostasis. Nat Aging 2, 726–741.

Li, Y., He, Y., Peng, J., Su, Z., Li, Z., Zhang, B., Ma, J., Zhuo, M., Zou, D., Liu, X., et al. (2021). Mutant Kras co-opts a proto-oncogenic enhancer network in inflammation-induced metaplastic progenitor cells to initiate pancreatic cancer. Nat Cancer 2, 49–65.

Lock, R., Roy, S., Kenific, C.M., Su, J.S., Salas, E., Ronen, S.M., and Debnath, J. (2011). Autophagy facilitates glycolysis during Ras-mediated oncogenic transformation. Mol Biol Cell 22, 165–178.

Loncle, C., Bonjoch, L., Folch-Puy, E., Lopez-Millan, M.B., Lac, S., Molejon, M.I., Chuluyan, E., Cordelier, P., Dubus, P., Lomberk, G., et al. (2015). IL17 Functions through the Novel REG3beta-JAK2-STAT3 Inflammatory Pathway to Promote the Transition from Chronic Pancreatitis to Pancreatic Cancer. Cancer Res 75, 4852–4862.

Love, M.I., Huber, W., and Anders, S. (2014). Moderated estimation of fold change and dispersion for RNA-seq data with DESeq2. Genome Biol 15, 550.

Ma, Z., Lytle, N.K., Chen, B., Jyotsana, N., Novak, S.W., Cho, C.J., Caplan, L., Ben-Levy, O., Neininger, A.C., Burnette, D.T., et al. (2022). Single-Cell Transcriptomics Reveals a Conserved Metaplasia Program in Pancreatic Injury. Gastroenterology 162, 604–620 e620.

Martincorena, I., Roshan, A., Gerstung, M., Ellis, P., Van Loo, P., McLaren, S., Wedge, D.C., Fullam, A., Alexandrov, L.B., Tubio, J.M., et al. (2015). Tumor evolution. High burden and pervasive positive selection of somatic mutations in normal human skin. Science 348, 880–886.

McWilliams, T.G., and Ganley, I.G. (2019). Investigating Mitophagy and Mitochondrial Morphology In Vivo Using mito-QC: A Comprehensive Guide. Methods Mol Biol 1880, 621–642.

McWilliams, T.G., Prescott, A.R., Montava-Garriga, L., Ball, G., Singh, F., Barini, E., Muqit, M.M.K., Brooks, S.P., and Ganley, I.G. (2018). Basal Mitophagy Occurs Independently of PINK1 in Mouse Tissues of High Metabolic Demand. Cell Metab 27, 439–449 e435.

Montava-Garriga, L., Singh, F., Ball, G., and Ganley, I.G. (2020). Semi-automated quantitation of mitophagy in cells and tissues. Mech Ageing Dev 185, 111196.

Morris, J.P.t., Cano, D.A., Sekine, S., Wang, S.C., and Hebrok, M. (2010). Beta-catenin blocks Kras-dependent reprogramming of acini into pancreatic cancer precursor lesions in mice. J Clin Invest 120, 508–520.

Morton, J.P., Timpson, P., Karim, S.A., Ridgway, R.A., Athineos, D., Doyle, B., Jamieson, N.B., Oien, K.A., Lowy, A.M., Brunton, V.G., et al. (2010). Mutant p53 drives metastasis and overcomes growth arrest/senescence in pancreatic cancer. Proc Natl Acad Sci U S A 107, 246–251.

Mukherjee, S., Partch, C.L., Lehotzky, R.E., Whitham, C.V., Chu, H., Bevins, C.L., Gardner, K.H., and Hooper, L.V. (2009). Regulation of C-type lectin antimicrobial activity by a flexible N-terminal prosegment. J Biol Chem 284, 4881–4888.

Mukherjee, S., Zheng, H., Derebe, M.G., Callenberg, K.M., Partch, C.L., Rollins, D., Propheter, D.C., Rizo, J., Grabe, M., Jiang, Q.X., and Hooper, L.V. (2014). Antibacterial membrane attack by a pore-forming intestinal C-type lectin. Nature 505, 103–107.

Muraro, M.J., Dharmadhikari, G., Grun, D., Groen, N., Dielen, T., Jansen, E., van Gurp, L., Engelse, M.A., Carlotti, F., de Koning, E.J., and van Oudenaarden, A. (2016). A Single-Cell Transcriptome Atlas of the Human Pancreas. Cell Syst 3, 385–394 e383.

Navas, C., Hernandez-Porras, I., Schuhmacher, A.J., Sibilia, M., Guerra, C., and Barbacid, M. (2012). EGF receptor signaling is essential for k-ras oncogene-driven pancreatic ductal adenocarcinoma. Cancer Cell 22, 318–330.

Neuhofer, P., Roake, C.M., Kim, S.J., Lu, R.J., West, R.B., Charville, G.W., and Artandi, S.E. (2021). Acinar cell clonal expansion in pancreas homeostasis and carcinogenesis. Nature 597, 715–719.

Nthiga, T.M., Kumar Shrestha, B., Sjottem, E., Bruun, J.A., Bowitz Larsen, K., Bhujabal, Z., Lamark, T., and Johansen, T. (2020). CALCOCO1 acts with VAMP-associated proteins to mediate ER-phagy. EMBO J 39, e103649.

Patro, R., Duggal, G., Love, M.I., Irizarry, R.A., and Kingsford, C. (2017). Salmon provides fast and bias-aware quantification of transcript expression. Nat Methods 14, 417–419.

Perera, R.M., Stoykova, S., Nicolay, B.N., Ross, K.N., Fitamant, J., Boukhali, M., Lengrand, J., Deshpande, V., Selig, M.K., Ferrone, C.R., et al. (2015). Transcriptional control of autophagy-lysosome function drives pancreatic cancer metabolism. Nature 524, 361–365.

Perez-Riverol, Y., Bai, J., Bandla, C., Garcia-Seisdedos, D., Hewapathirana, S., Kamatchinathan, S., Kundu, D.J., Prakash, A., Frericks-Zipper, A., Eisenacher, M., et al. (2022). The PRIDE database resources in 2022: a hub for mass spectrometry-based proteomics evidences. Nucleic Acids Res 50, D543–D552.

Rosenfeldt, M.T., O’Prey, J., Morton, J.P., Nixon, C., MacKay, G., Mrowinska, A., Au, A., Rai, T.S., Zheng, L., Ridgway, R., et al. (2013). p53 status determines the role of autophagy in pancreatic tumour development. Nature 504, 296–300.

Schlesinger, Y., Yosefov-Levi, O., Kolodkin-Gal, D., Granit, R.Z., Peters, L., Kalifa, R., Xia, L., Nasereddin, A., Shiff, I., Amran, O., et al. (2020). Single-cell transcriptomes of pancreatic preinvasive lesions and cancer reveal acinar metaplastic cells’ heterogeneity. Nat Commun 11, 4516.

Smith, M., Salomo-Coll, C., Muir, M., and Wilkinson, S. (2022). Analysis of Pancreatic Acinar Protein Solubility in Autophagy-Deficient Mice. Methods Mol Biol 2445, 243–253.

Smith, M.D., Harley, M.E., Kemp, A.J., Wills, J., Lee, M., Arends, M., von Kriegsheim, A., Behrends, C., and Wilkinson, S. (2018). CCPG1 Is a Non-canonical Autophagy Cargo Receptor Essential for ER-Phagy and Pancreatic ER Proteostasis. Dev Cell 44, 217–232 e211.

Stephani, M., Picchianti, L., Gajic, A., Beveridge, R., Skarwan, E., Sanchez de Medina Hernandez, V., Mohseni, A., Clavel, M., Zeng, Y., Naumann, C., et al. (2020). A cross-kingdom conserved ER-phagy receptor maintains endoplasmic reticulum homeostasis during stress. Elife 9.

Storz, P. (2017). Acinar cell plasticity and development of pancreatic ductal adenocarcinoma. Nat Rev Gastroenterol Hepatol 14, 296–304.

Subramanian, A., Tamayo, P., Mootha, V.K., Mukherjee, S., Ebert, B.L., Gillette, M.A., Paulovich, A., Pomeroy, S.L., Golub, T.R., Lander, E.S., and Mesirov, J.P. (2005). Gene set enrichment analysis: a knowledge-based approach for interpreting genome-wide expression profiles. Proc Natl Acad Sci U S A 102, 15545–15550.

Tosti, L., Hang, Y., Debnath, O., Tiesmeyer, S., Trefzer, T., Steiger, K., Ten, F.W., Lukassen, S., Ballke, S., Kuhl, A.A., et al. (2021). Single-Nucleus and In Situ RNA-Sequencing Reveal Cell Topographies in the Human Pancreas. Gastroenterology 160, 1330–1344 e1311.

Vargas, J.N.S., Hamasaki, M., Kawabata, T., Youle, R.J., and Yoshimori, T. (2023). The mechanisms and roles of selective autophagy in mammals. Nat Rev Mol Cell Biol 24, 167–185.

Yang, A., Rajeshkumar, N.V., Wang, X., Yabuuchi, S., Alexander, B.M., Chu, G.C., Von Hoff, D.D., Maitra, A., and Kimmelman, A.C. (2014). Autophagy is critical for pancreatic tumor growth and progression in tumors with p53 alterations. Cancer Discov 4, 905–913.

Yang, S., Wang, X., Contino, G., Liesa, M., Sahin, E., Ying, H., Bause, A., Li, Y., Stommel, J.M., Dell’antonio, G., et al. (2011). Pancreatic cancers require autophagy for tumor growth. Genes Dev 25, 717–729.

Zhang, H., Corredor, A.L.G., Messina-Pacheco, J., Li, Q., Zogopoulos, G., Kaddour, N., Wang, Y., Shi, B.Y., Gregorieff, A., Liu, J.L., and Gao, Z.H. (2021). REG3A/REG3B promotes acinar to ductal metaplasia through binding to EXTL3 and activating the RAS-RAF-MEK-ERK signaling pathway. Commun Biol 4, 688.

