## Supplementary Figures for "Dysproteostasis primes pancreatic epithelial state changes in *KRAS*-mediated oncogenesis"

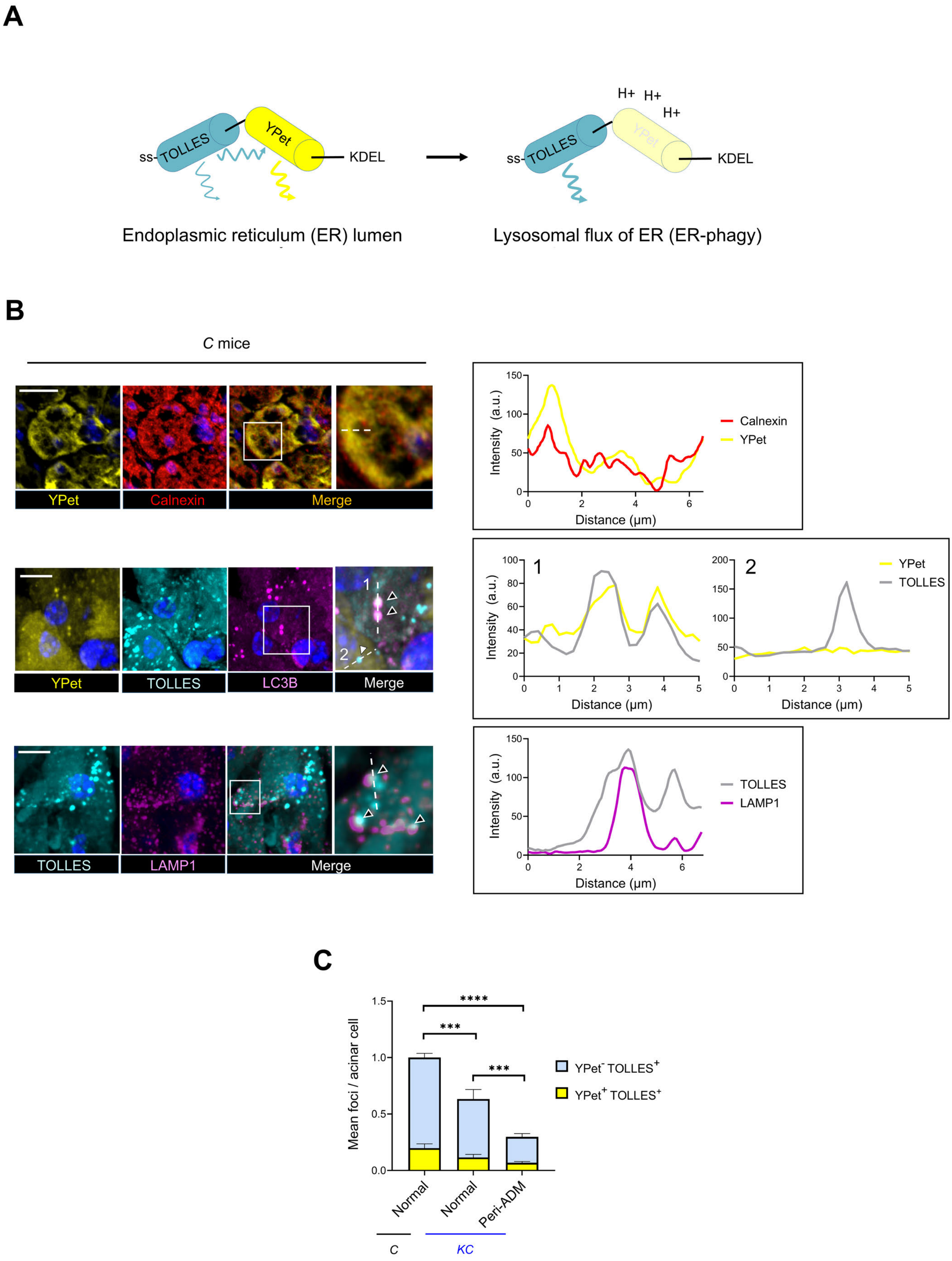


**Supplementary Figure 1, corresponds with Figure 1**

### Supplementary Figure 1, corresponds with Figure 1

**A)** Schematic of ER-phagy reporter fusion protein ss-TOLLES-YPet-KDEL. N-ter signal sequence (ss) and C-ter ER retention sequence (KDEL) target the protein to the ER lumen. YPet and TOLLES fluorescences are evident within the ER and within autophagosomes derived from the ER. Only TOLLES fluorescence is detected in autolysosomes derived frommic completion of ER-phagy; when ER containing autophagosomes fuse with lysosomes the low pH quenches YPet and additionally relives FRET-mediated partial suppression of the TOLLES signal by YPet.

**B)** 10-week-old mice were infected with rAAV expressing ss-TOLLES-YPet-KDEL. Here are shown representative spinning-disk confocal microscopy images of control *C* mouse pancreatic acini (*Pdx1-Cre* only). The reporter is localised to the ER (co-staining for Calnexin, top row). Puncta positive for both YPet and TOLLES are also observed (second row, black arrowheads, example 1), although more frequently TOLLES-only puncta are seen (second row, white arrowheads, example 2). The former co-stain for LC3B (example 1), suggesting their autophagosomal identity, whereas the latter are not LC3B positive (example 2). The latter TOLLES foci are found within the LAMP1 compartment, consistent with completion of ER-phagy within the lysosome (third row, black arrowheads). Dotted lines denote the paths of the individual fluorescence traces graphed on the right-hand side of the panel.

Scale bars = 20 μm.

**C)** Imaris image analysis was used to extract the number of YPet$^{+}$ TOLLES$^{+}$ and TOLLES-only foci in each acinar cell, from morphologically normal lobules or lobules harbouring sporadic ADM, shown here normalised to sum of both focus types in control animals (*C*). Proportional reductions in both YPet$^{+}$ TOLLES$^{+}$ foci (ER-containing autophagosomes) and TOLLES-only foci (ER-phagy endpoint lysosomes) in *KC* mice indicate that ER-phagy is inhibited at an early stage after *Kras* activation, prior to generation of ER-containing autophagosomes.(n = 5 mice, mean counts per acinar cell for each animal, > 4000 total cells per group, ± S.E.M., 2-way ANOVA and Holm-Šidák post-hoc tests, comparing YPet$^{-}$ TOLLES$^{+}$ foci, *** = $p \leq 0.001$, **** = $p \leq 0.0001$).

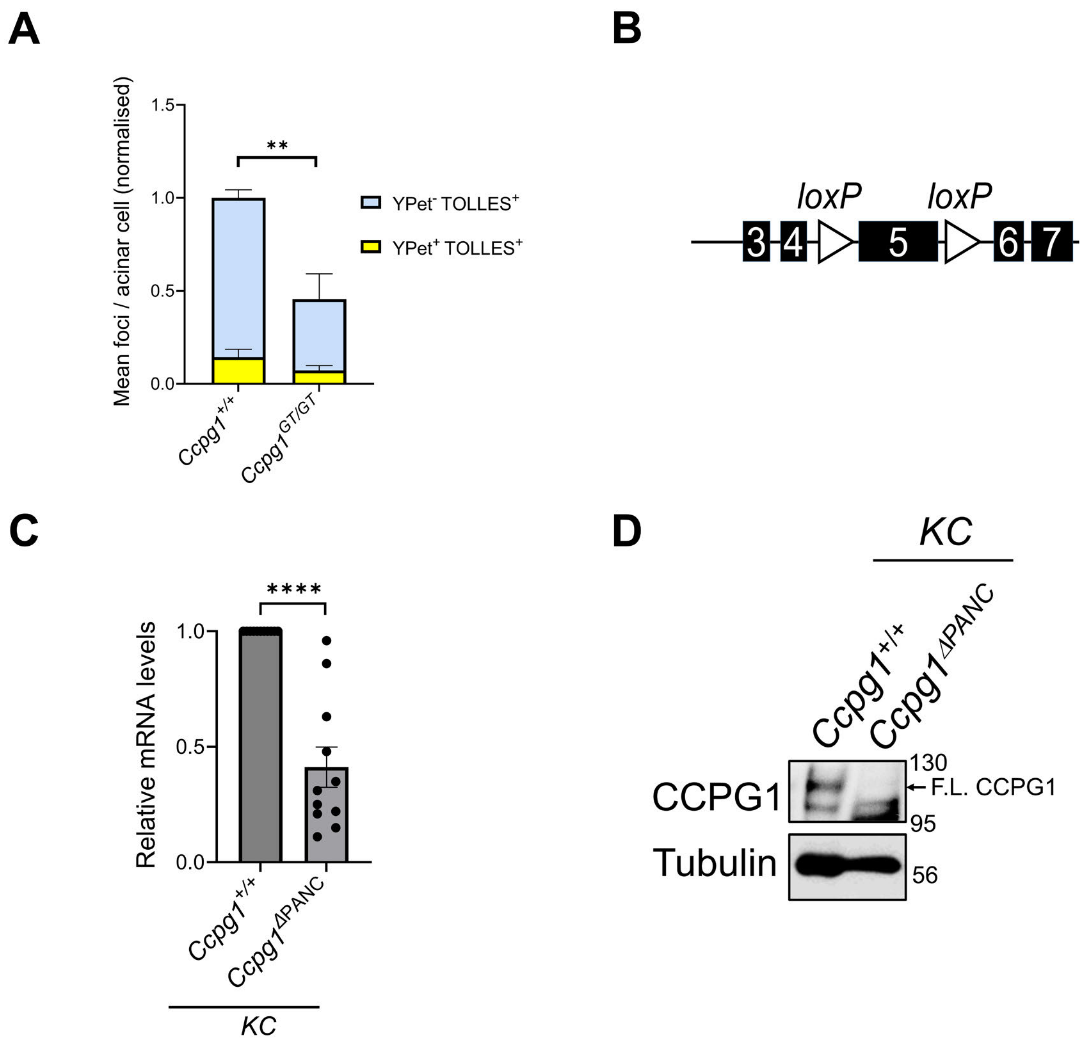


### Supplementary Figure 2, corresponds with Figure 2

**A)** Imaris image analysis was used to extract the number of double-fluorescent (YPet$^+$ TOLLES$^+$) and TOLLES-only (YPet$^-$ TOLLES$^+$) foci in acinar cells of control (*Ccpg1*$^{+/+}$) and germline *Ccpg1*-deficient mice (*Ccpg1*$^{GT/GT}$), two weeks after injection with AAV expressing the ER-phagy reporter ss-YPet-TOLLES-KDEL. Values are normalised to the sum of both focus types in the control animal group. Proportional reductions in both YPet-TOLLES foci (ER-containing autophagosomes) and TOLLES-only foci (ER-phagy endpoint lysosomes) in *Ccpg1*$^{GT/GT}$ mice indicate that ER-phagy is inhibited at an early stage prior to generation of ER-containing autophagosomes (n = 3 mice, mean counts per acinar cell for each animal, > 3300 total cells per group, ± S.E.M., 2-way ANOVA and Holm-Šidák post-hoc test, comparing YPet$^-$ TOLLES$^+$ foci, ** = p ≤ 0.01). **B)** Schematic of the *Ccpg1*$^{flox}$ conditional mouse allele. This permits pancreatic epithelial specific knockout of *Ccpg1* on the *KC* and *C* backgrounds (*Pdx1-Cre*). Exons 3 to 7 are depicted. **C-D)** Validation of *Ccpg1* loss in mice where *Ccpg1*$^{flox}$ is bred to homozygosity upon a *KC* background (generating *KC Ccpg1*$^{\Delta PANC}$ mice), in **C** via *Ccpg1* qRT-PCR (n = 11, ± S.E.M., 1-sample t-test, ****= p ≤ 0.0001) or **D** via immunoblot of whole pancreatic lysate (F.L. = full-length).

**A**

| Gene | Encoded protein |
|---|---|
| *Agr2* | Secreted |
| *Ang* | Secreted |
| *Cpn1* | Secreted |
| *Dnajb9* | ER co-chaperone |
| *Dnajc3* | ER co-chaperone |
| *Hapln4* | Secreted |
| *Igf1* | Secreted |
| *Il1rn* | Secreted |
| *Mia* | Secreted |
| *Pdgfa* | Secreted |
| *Pla2g1b* | Secreted |
| *Reg3a* | Secreted |
| *Reg3b* | Secreted |
| *Reg3d* | Secreted |
| *Rnase4* | Secreted |
| *Ttr* | Secreted |
| *Twsg1* | Secreted |
| *Zg16* | Secreted |

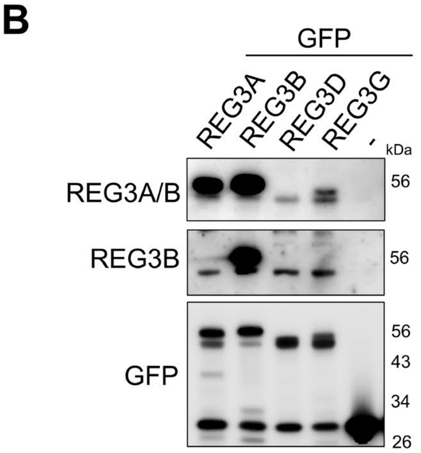


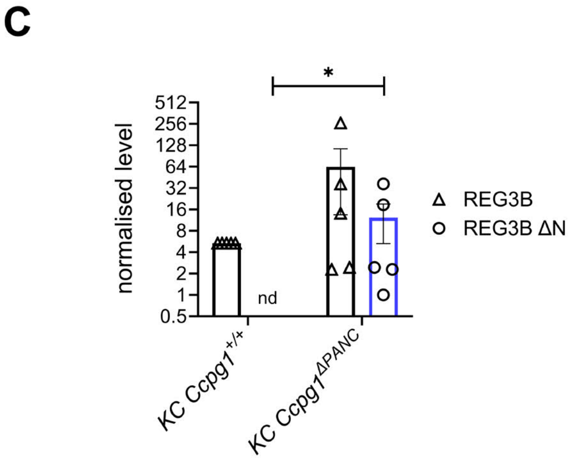


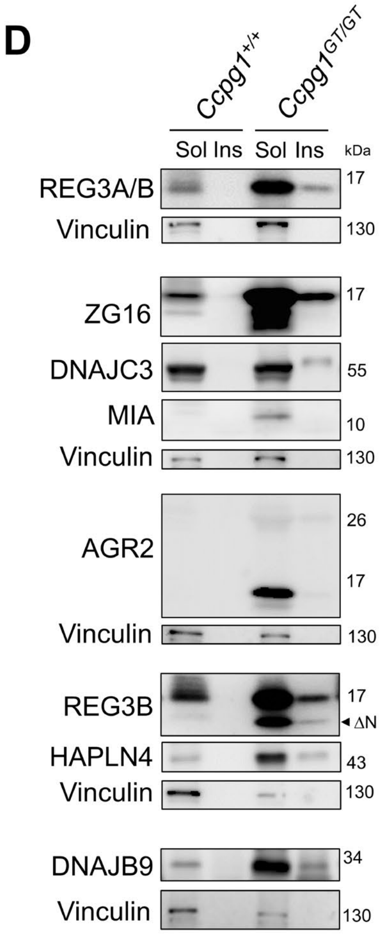


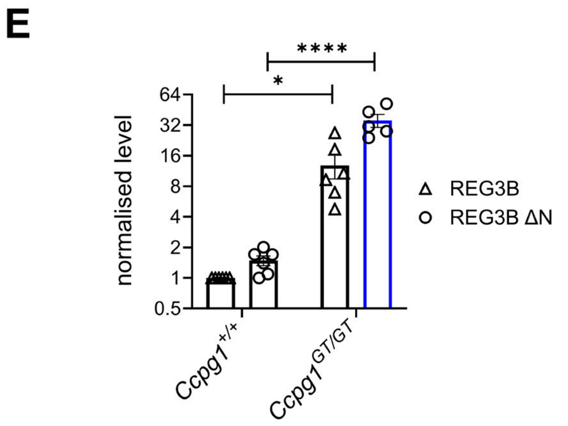


**Supplementary Figure 3, corresponds with Figure 3**

### Supplementary Figure 3, corresponds with Figure 3

**A)** Ontology of proteins detected in Figure 3A showing predicted compartmentalisation in the ER lumen.

**B)** Immunoblot validation of specificity of antibodies used in this study for detection of REG3 proteins, assessing recognition of C-terminally GFP-fused murine REG3 paralogues transfected into PDAC cells.

**C)** Quantification of full-length REG3B and the aggregation-prone, presumed N-terminally cleaved ΔN subspecies (ΔN), summated across detergent-soluble and insoluble fractions of whole pancreatic lysates from the *KC* $Ccpg1^{+/+}$ and *KC* $Ccpg1^{\Delta PANC}$ animals, as described in Fig. 3B-C (n = 5, ± S.E.M., normalised to REG3B in *KC* $Ccpg1^{+/+}$, nd = non-determined as there was no detectable REG3B in any replicate, 1-sample t-test versus hypothetical mean of 0, * = $p \leq 0.05$).

**D)** Representative immunoblot corresponding to Fig. 3E (sol/insol = detergent soluble/insoluble).

**E)** Quantification of full-length REG3B and aggregation-prone, presumed N-terminally cleaved ΔN subspecies (ΔN), summated across detergent-soluble and -insoluble fractions of whole pancreatic lysates from $Ccpg1^{+/+}$ and $Ccpg1^{GT/GT}$ mice, as described in Fig. 3E and Supp Fig. 3D (n = 6, ± S.E.M., normalised to REG3B in $Ccpg1^{+/+}$, 1-sample t-test for REG3B, Student's t-test for REG3BΔN, * = $p \leq 0.05$, **** = $p < 0.0001$).

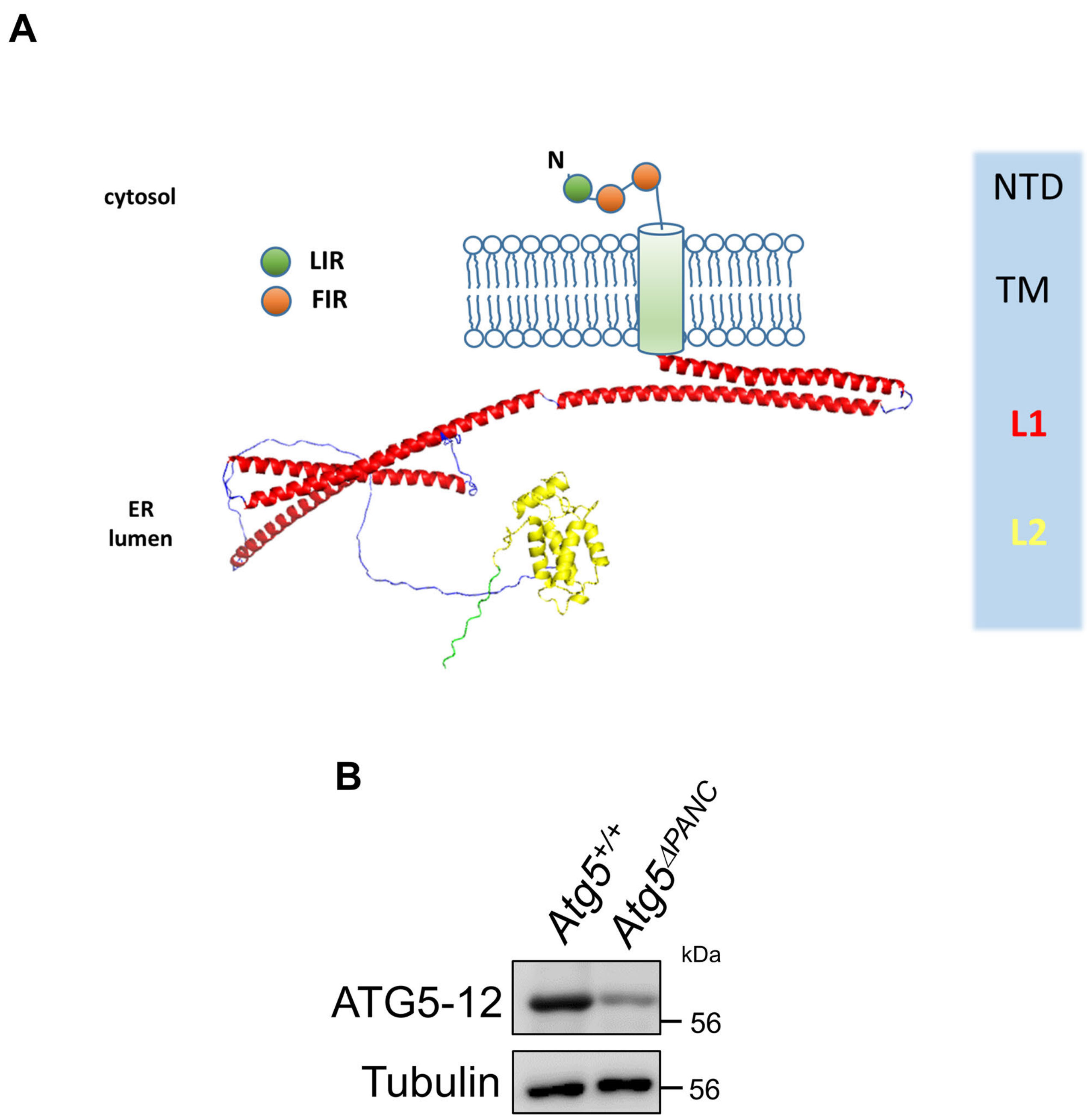


**Supplementary Figure 4, corresponds with Figure 4**

**A)** AlphaFold v2 tertiary structure prediction of the luminal C-terminal region of murine CCPG1 (adjoined to a schematic representation of the N-terminal intrinsically disordered domain and transmembrane region: NTD = N-terminal domain, TM = transmembrane region, L1/2 = ER luminal domains 1 and 2, LIR = LC3-interacting region and FIR = FIP200-interacting region, both required for ER-phagy). **B)** Immunoblot confirmation of reduced *Atg5* expression in pancreatic epithelia of 8-week-old *Atg5*$^{\Delta PANC}$ (*Pdx1-Cre Atg5*$^{flox/flox}$) mice.

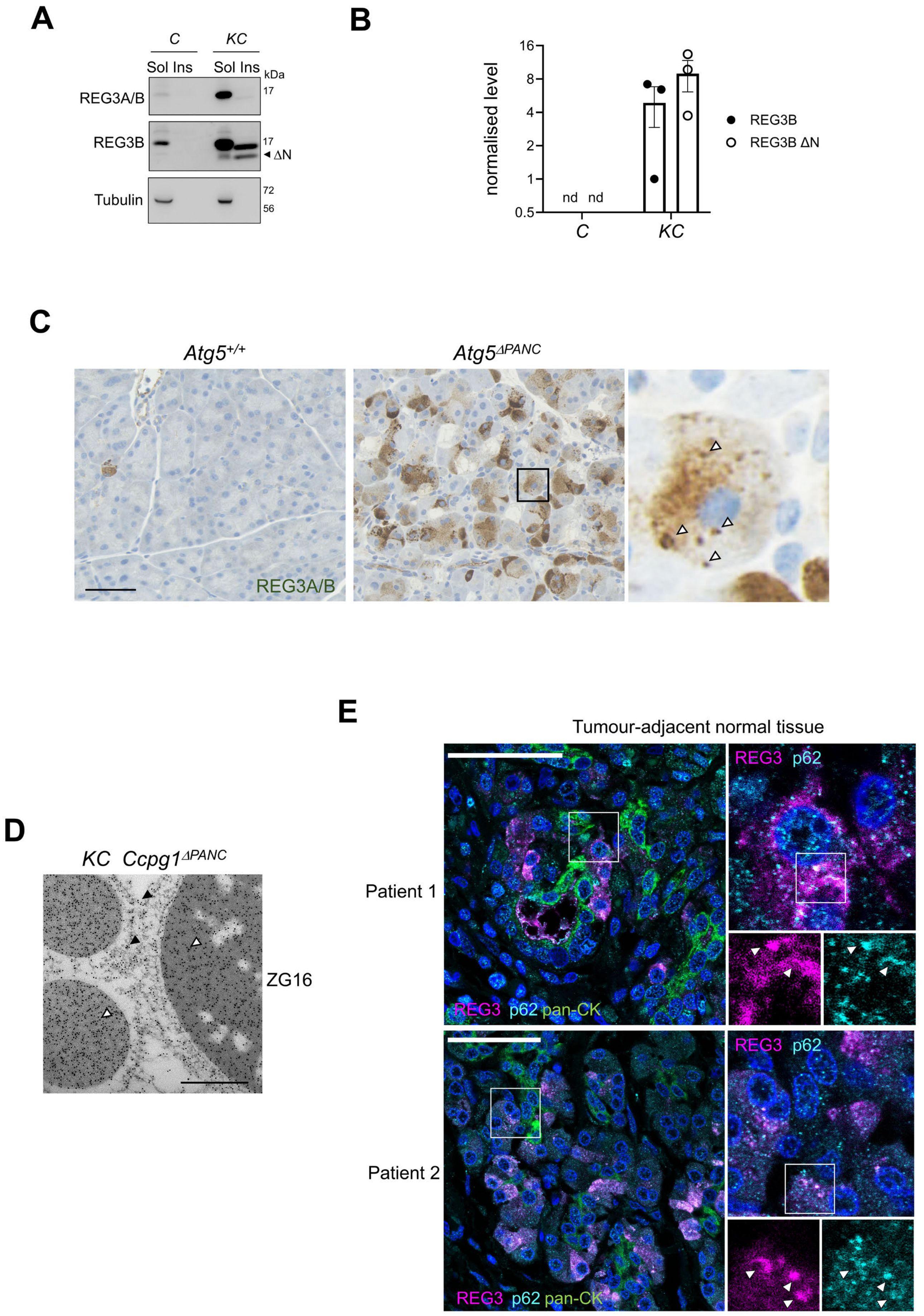


**Supplementary Figure 5, corresponds with Figure 5**

### Supplementary Figure 5, corresponds with Figure 5

**A-B)** Accumulation of REG3A/B protein in detergent-soluble and insoluble-fractions (NP40 Sol/Insol) from whole pancreas of 18-week-old *C* and *KC* mice, representative immunoblot shown in **A**, ΔN = N-terminally cleaved REG3B. Quantification in **B** of insoluble full-length REG3B and aggregation-prone ΔN subspecies (n = 3, ± S.E.M., nd = not determined due to no detectable signal in any replicate).

**C)** Immunohistochemistry for REG3A/B on formalin-fixed paraffin embedded (FFPE) pancreata from 4-week-old *Atg5*$^{+/+}$ (*Pdx1-Cre Atg5*$^{+/+}$) or *Atg5*$^{\Delta PANC}$ (*Pdx1-Cre Atg5*$^{flox/flox}$) mice. Arrowheads indicate exemplar focal signals. Scale bar = 500 μm.

**D)** Immunogold transmission electron microscopy of ZG16 in acinar cells within pancreata of 16-week-old *KC Ccpg1*$^{\Delta PANC}$ mice. Arrowheads indicate gold particles marking accumulation within granular aggregates (white arrowheads) or along the tubular ER network (black arrowheads). Scale bar = 500 nm.

**E)** Confocal immunofluorescence microscopy of ADM and surrounding acinar cells in PDAC-adjacent normal acinar tissue (panCK = pan-cytokeratin). Arrowheads indicate exemplar focal signals for REG3A/G (human orthologues of REG3A/B) and p62. At least one ADM-acinar protein aggregate region was detected in 8 of 22 x 2mm diameter tissue cores analysed. Scale bar = 50 μm.

**Supplementary Figure 6, corresponds with Figure 6 (bulk mRNA-Seq data)**

A

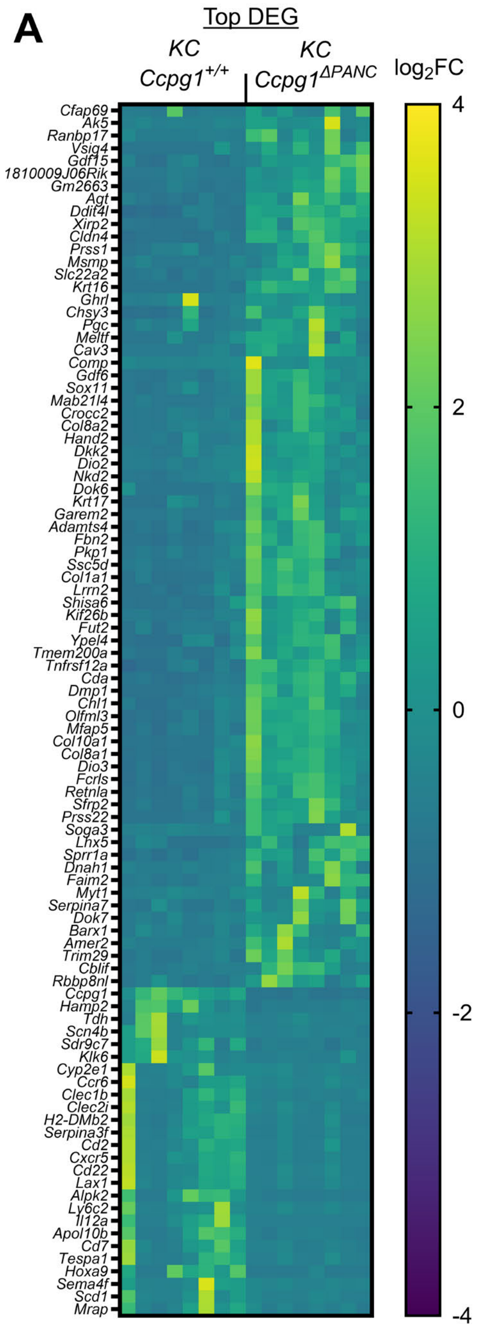


B

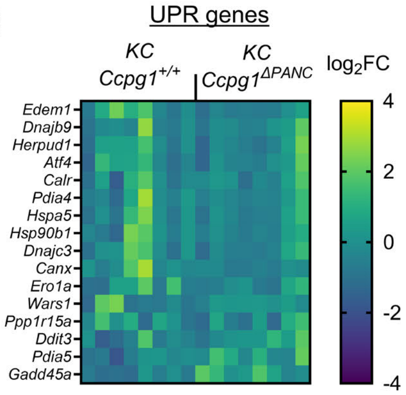


C

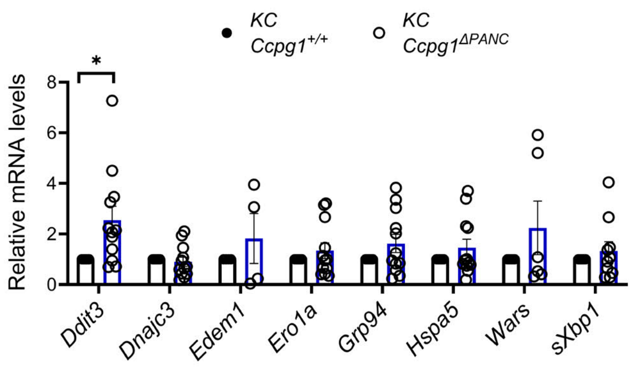


D

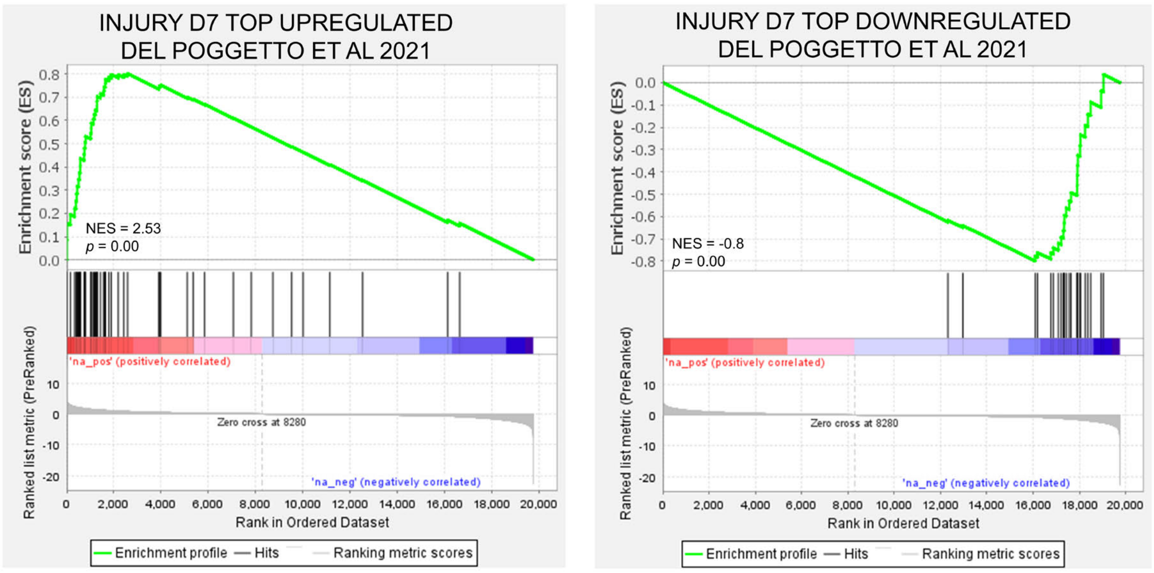

### Supplementary Figure 6, corresponds with Figure 6 (bulk mRNA-Seq data)

**A-B)** Heatmaps of all top differentially expressed genes (DEG), **A**, and UPR target genes, **B,** after bulk mRNA-Seq of whole pancreatic RNA from 10-week-old *KC Ccpg1*$^{+/+}$ and *KC Ccpg1*$^{\Delta PANC}$ mice. FC = fold change.

**C)** qRT-PCR analysis of s-*Xbp1* (spliced *Xbp1*) and UPR target genes in whole pancreatic RNA from 10-week-old *KC Ccpg1*$^{\Delta PANC}$ mice, normalising to *KC* controls (n = 4-12, 1-sample t-tests, * = $p \leq 0.05$, not shown= $p > 0.05$).

**D)** Example individual gene-set enrichment analysis (GSEA) scores corresponding to Fig. 6A, comparing whole pancreatic extracts of 10-week-old *KC Ccpg1*$^{\Delta PANC}$ with *KC Ccpg1*$^{+/+}$ mice. These examples are, specifically, d7 post-injury signatures curated from (Del Poggetto et al., 2021). NES = normalised enrichment score.

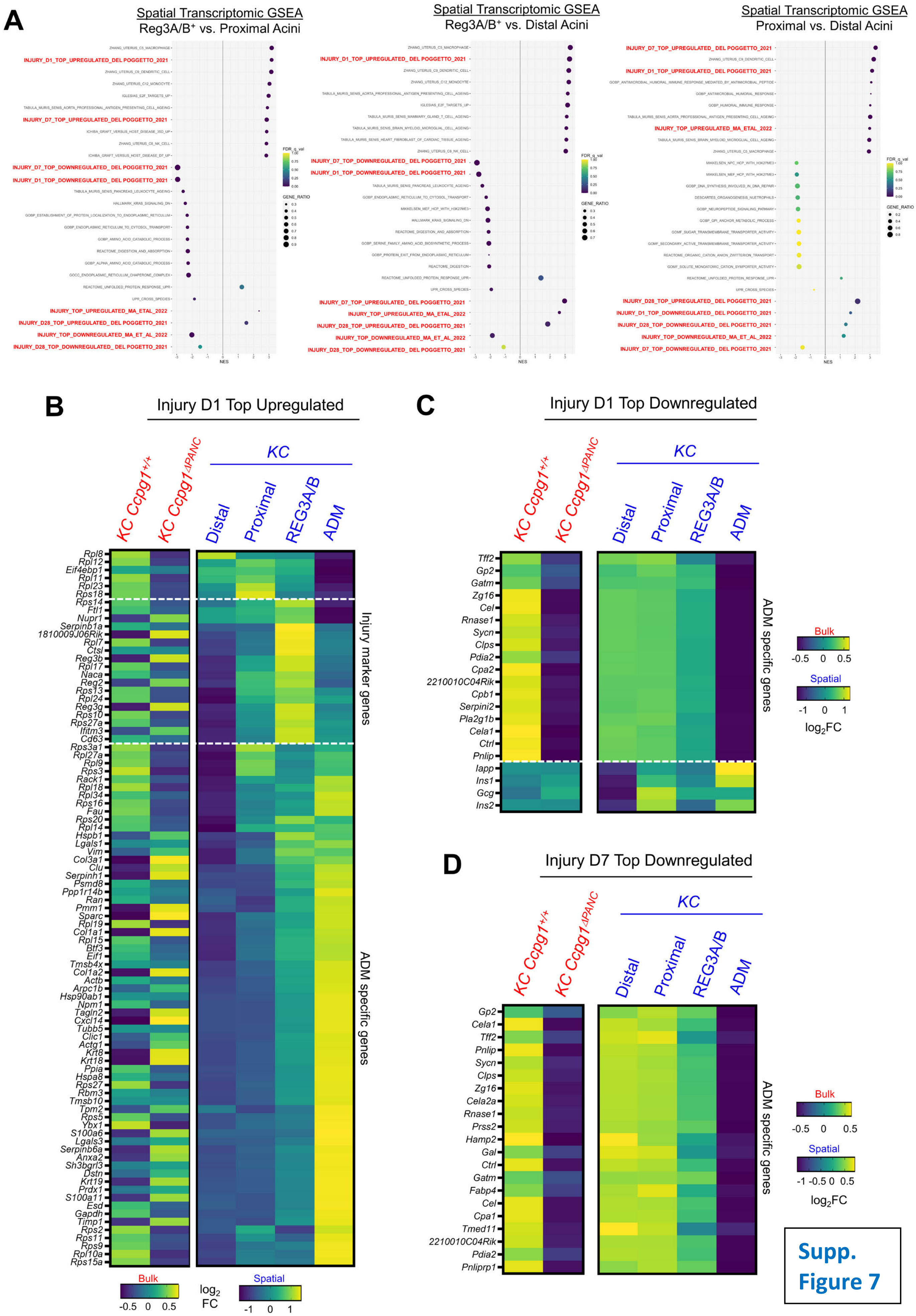


Supp. Figure 7

#### Supplementary Figure 7, corresponds with Figure 6 (spatial transcriptomic data)

**A)** Summary of all gene-set enrichment analyses (GSEA) performed in analysis of spatial transcriptomic data described in Fig. 6C-E, performing pairwise comparisons of indicated cell populations. The top 20 signatures ordered by NES (normalised enrichment score) are ranked alongside UPR signatures and collated injury signatures (emboldened in red). FDR = false-discovery rate.

**B-D)** Heatmaps comparing transcript-by-transcript changes for additional injury gene signatures from (Del Poggetto et al., 2021) to that represented in Fig. 6F, both within bulk mRNA-Seq data from wild-type and *Ccpg1*-deficient *KC* mice (red column labels), and across the different cell populations isolated by spatial transcriptomics from *KC* mice (blue column labels).

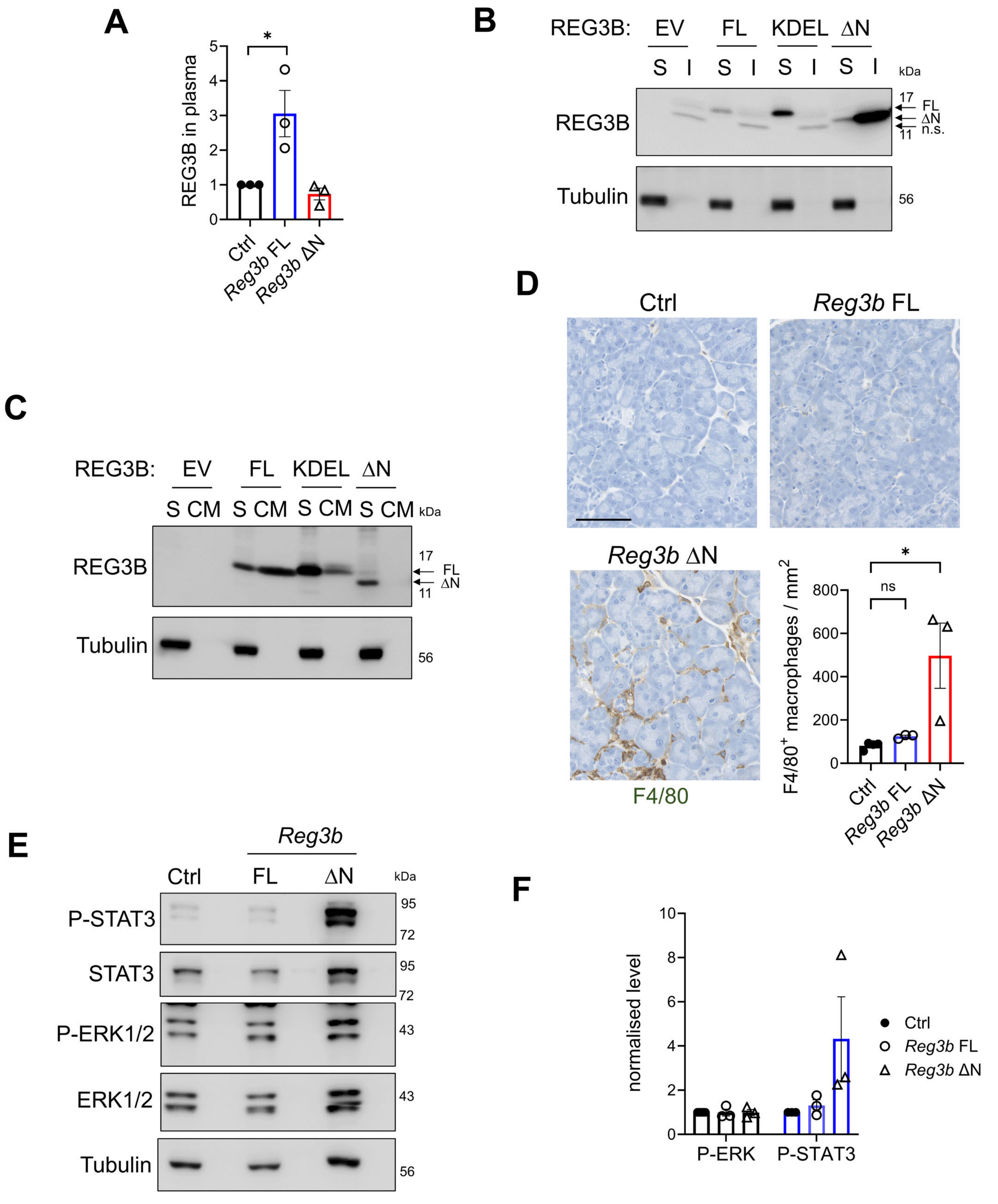


**Supplementary Figure 8, corresponds with Figure 7**

### Supplementary Figure 8, corresponds with Figure 7

**A)** Quantification of plasma REG3B blots from Fig. 7D (n = 3, ± S.E.M., 1-sample t-tests vs. Ctrl, * = $p \leq 0.05$, not shown = $p > 0.05$).

**B)** Immunoblotting for REG3B in whole HEK293FT cell detergent (NP40)-soluble (S) and -insoluble (I) extracts after transfection with empty vector (EV) or *Reg3b* forms: FL (full-length); KDEL (full-length fused to C-terminal KDEL ER retention signal to impair secretion; ΔN (N-terminally processed aggregation-prone). n.s. = non-specific band.

**C)** Immunoblot of conditioned culture medium from HEK293FT transfected in **B** (S = detergent-soluble fraction from **B** for reference, CM = conditioned medium).

**D)** $F4/80^+$ immunohistochemistry to detect macrophages in FFPE sections from the mice described in Figure 7, alongside quantifications performed on randomly-selected, morphologically-normal lobules (± S.E.M., 1-way ANOVA and Holm-Šidák post-hoc tests vs. Ctrl, * = $p \leq 0.05$, ns = $p > 0.05$). Scale bar = 100 μm.

**E-F)** Immunoblot of whole pancreas lysate (detergent-soluble fraction) to assess JAK/STAT3 signalling and MAPK kinase (ERK1/2) signalling in mice described in Figure 7 (P-STAT3 = phospho-Y704-STAT3, P-ERK1/2 = phospho-T202/Y204 ERK1/2), representative blot in **E**, quantification in **F** (n = 3, ± S.E.M.).
